# Coiled-coil homo-oligomerization and disaggregase Hsp104 act in parallel to stabilize orphan septins

**DOI:** 10.64898/2026.03.09.710472

**Authors:** Italo A Cavini, Randi M Yeager, Alejandra Velasquez, Andressa P A Pinto, Ana Paula U Araujo, Richard C Garratt, Michael A McMurray

## Abstract

Multiple septin family proteins co-assemble with strict subunit stoichiometry into hetero-oligomers. In the absence of native septin partners, purified septins aggregate in vitro, and “orphan” septins are found in pathological aggregates associated with neurodegenerative diseases. Cytosolic chaperones bind the septin GTPase domain to promote on-pathway septin folding but it was unclear how cells manage orphan septins to maintain subunit stoichiometry. Most septins have C-terminal domains (CTDs) that form heteromeric coiled coils within or between septin complexes. Here we present evidence that orphan yeast septins are protected from proteasomal degradation by forming transient coiled-coil homodimers and trimers and, in parallel, by the disaggregase chaperone Hsp104. Septins unable to undergo CTD-mediated homo-oligomerization require Hsp104 to accumulate to super-stoichiometric levels. We show that septin-encoding mRNAs differ in copy number per cell and subcellular localization, creating opportunities for transient subunit imbalances. These findings reveal a novel role for coiled coils and the proteostasis machinery in the fidelity of higher-order septin assembly.

## INTRODUCTION

The septin family of eukaryotic filament-forming proteins function in cellular morphogenesis, intracellular trafficking, and cytokinesis, among other processes (Mostowy and Cossart, 2012). In humans and budding yeast, filaments form via polymerization of rod-shaped septin hetero-octamers into which two molecules of each of four septin subunits assemble via alternating interfaces on opposite faces of a globular GTPase domain (Sirajuddin *et al*., 2007; Bertin *et al*., 2008; McMurray and Thorner, 2019; Mendonça *et al*., 2019; Soroor *et al*., 2021) (Fig. 1 A). Neuronal septins involved in synaptic plasticity have been found in protein aggregates associated with human neurodegenerative diseases (reviewed in (Marttinen *et al*., 2015; Alkhanjari *et al*., 2025)), and in the absence of native hetero-dimerization partners, human septins can form amyloid-like fibrils in vitro with properties indicative of cross-β sheet aggregation (Garcia *et al*., 2007; Pissuti Damalio *et al*., 2012; Kumagai *et al*., 2019). Indeed, a septin was the first identified substrate of Parkin, an E3 ubiquitin-protein ligase, mutations in which are responsible for early-onset Parkinson disease (Zhang *et al*., 2000). A model that emerges from these findings is that excess of a single septin protein creates a scenario in which unoccupied hetero-oligomerization interfaces mediate the assembly of dysfunctional homo-oligomers that can perturb normal septin function. However, we know few molecular details about how cells manage septin homo-oligomerization to maintain normal function.

**Figure 1.**
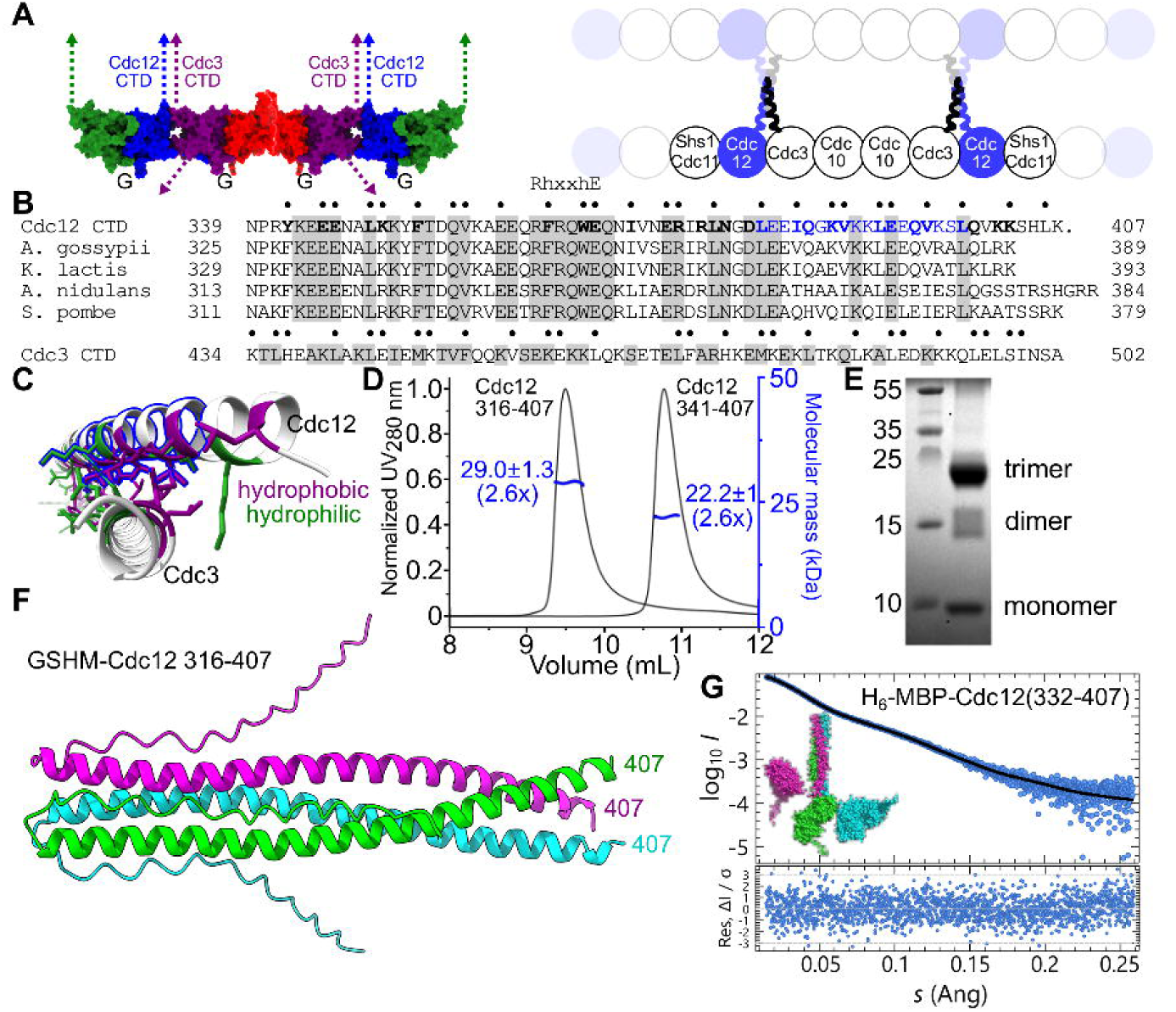
Homotrimerization of the Cdc12 C-terminal domain. (A) Schematic illustration of budding yeast septin hetero-octamer and filament organization focusing on putative roles for coiled coils formed by the C-terminal domains of Cdc3 and Cdc12. At left is a model of the septin hetero-octamer from *S. cerevisiae* based on the crystal structure of a Shs1–Cdc12–Cdc3–Cdc10 hetero-tetramer lacking the extended N- and C-terminal domains (PDB 8PFH (Grupp *et al*., 2024)). The missing domains are illustrated as dashed arrows. “CTD”, C-terminal domain; “G”, so-called G interface between septins. The cartoon at right shows end-on-end hetero-octamer polymerization and also the putative four-helix coiled coils that have been proposed to mediate lateral pairing of adjacent filaments (Bertin *et al*., 2008). (B) Sequence alignment of the Cdc12 CTD from *S. cerevisiae* with homologs from other fungal species, with the sequence of the Cdc3 CTD shown below. Gray shading indicates identity among all five species (other Cdc3 sequences are not shown). Bullet points above residues indicate those that are predicted to make contacts in a heterodimeric Cdc3–Cdc12 coiled coil. Bold residues are predicted to make contacts in a homotrimeric Cdc12 coiled coil. The “RhxxhE” motif known to mediate homotrimerization in non-septin coiled coils (Kammerer *et al*., 2005) is also indicated above the sequence in Cdc12 that matches this motif. Blue residues comprise the amphipathic helix that is necessary and sufficient for the curvature preference of membrane binding by yeast hetero-octamers (Cannon *et al*., 2019; Woods *et al*., 2021). (C) The heterodimeric Cdc3–Cdc12 coiled coil as predicted by AlphaFold3, showing Cdc3 residues 434-503 and Cdc12 residues 339-406 and the hydrophobic (isoleucine, leucine, methionine, phenylalanine, tryptophan, and valine, in purple) and charged/hydrophilic (arginine, aspartate, glutamate, lysine, in green) side chains that make predicted contacts. The blue outline indicates the residues of the Cdc12 amphipathic helix that are predicted to interact with the inner leaflet of the plasma membrane. (D) SEC-MALS analysis of purified Cdc12 CTD at 300 µM. Both purified versions also include the residual thrombin cleavage sequence GSHM at the N terminus. Blue values show MALS-derived molecular mass and, in parenthesis, the relative mass compared to the monomer. SEC indicates non-resolved peaks corresponding to a mixture of dimers and trimers. Buffer employed was 50 mM Tris, 300 mM NaCl, pH 8.0 and protein concentration at injection was 0.3 mM. (E) SDS-PAGE analysis of the smaller Cdc12 CTD fragment (341-407, 9.3 kDa monomeric mass) following crosslinking with BS3. Left lane contains molecular weight standards (Promega Broad Range Marker); values indicate band masses. Labels indicate presumed identity of the Cdc12 CTD bands. (F) AlphaFold3 prediction of trimers of the Cdc12 CTD (residues 316-407) where residue 407 is labeled, and including the four residues (GSHM) left from protease cleavage during purification. (G) EOM-NNLSJOE fit and residuals of the SAXS dataset representing the top of the SEC elution peak of MBP-Cdc12CC+8 using parallel dimers and trimers. Inset, model of the most abundant species (27%).

In addition to the GTPase domain, most septins have extended CTDs with the propensity to form coiled coils (reviewed in (Cavini *et al*., 2021)). Septin coiled-coil-forming sequences are the targets of viral proteases that perturb cellular function (Li *et al*., 2019; Lee *et al*., 2023), but it remains unclear exactly how septin coiled coil assembly fits into the mechanism of septin rod and filament assembly (Sala *et al*., 2016). For example, in addition to parallel hetero-dimeric coiled coils, it has been proposed that the CTDs of the budding yeast septins Cdc3 and Cdc12 form anti-parallel four-helix coiled-coil bundles when filaments pair side-by-side (Bertin *et al*., 2008) (Fig. 1A). Both may be true: in the coiled coils formed by human septins both parallel and anti-parallel arrangements are energetically favorable and individual septins may switch between them (Leonardo *et al*., 2021).

Such “metastability” may be important in allowing dynamic changes in septin organization/assembly. Indeed, a short helical region of the CTD of Cdc12 (residues 382-399) has amphipathic properties that allow septin rods to sense membrane curvature (Cannon *et al*., 2019; Woods *et al*., 2021) (Fig. 1 A). The same hydrophobic Cdc12 residues predicted to be immersed in the inner leaflet of the plasma membrane are also predicted to be engaged in a heterodimeric coiled coil with the CTD of Cdc3 (Fig. 1 B,C). If the same Cdc12 molecule participates in both kinds of interactions, the Cdc12–Cdc3 coiled coil must undergo some degree of “unzipping” when a rod or filament binds a curved membrane, which metastability would facilitate. Unzipping would not destabilize septin rods or filaments, since Cdc3 and Cdc12 interact stably within hetero-octamers even when Cdc12 residues 318-407 are missing (Bertin *et al*., 2010). The CTD of septin Shs1 mediates interaction with the CTD of septin Cdc11 (Versele *et al*., 2004; Finnigan *et al*., 2015b) and also harbors its own curvature-sensing amphipathic helix (Woods *et al*., 2021) as well as a predicted coiled-coil-forming sequence that is sufficient to recruit a non-septin protein to septin filaments in vivo (Finnigan *et al*., 2015a, 2015b). Hence septin coiled-coil-forming sequences may engage in multiple distinct modes of interaction with other molecules before and/or after incorporation of a septin into a hetero-octameric rod, but such dynamic interactions have not yet been defined.

At steady-state, there are very few septin monomers or sub-octameric septin hetero-oligomers in wild-type budding yeast cells (Frazier *et al*., 1998; Vrabioiu *et al*., 2004; Farkasovsky *et al*., 2005; Bridges *et al*., 2014; Johnson *et al*., 2015) and in wild-type human cells septins exist solely as hetero-hexamers or -octamers (Sellin *et al*., 2011b, 2011a, 2014; Kuzmić *et al*., 2022). We proposed that Cdc12 normally acts as a slowly translated platform for efficient co-translational assembly of yeast septin hetero-octamers (Hassell *et al*., 2022). Since hetero-octamer assembly requires strict 2:2:2:2 stoichiometry of septin subunits, excess Cdc12 would complete translation with one or more interfaces left unengaged with a partner septin. Cytosolic chaperone proteins co-translationally bind nascent septins as the GTPase domain folds and compete transiently with partner septins to engage the G interface (Johnson *et al*., 2015; Stein *et al*., 2019; Denney *et al*., 2021; Hassell *et al*., 2022), which encompasses the GTP-binding pocket (Fig. 1 A) and includes a region with high predicted propensity for β aggregation (Pissuti Damalio *et al*., 2012; Hassell *et al*., 2022). Engaging the G interface in a native septin-septin heterodimer prevents human septin aggregation in vitro (Kumagai *et al*., 2019).

By experimentally overexpressing Cdc12, we showed that post-translational hetero-octamer assembly is possible but it requires cytosolic chaperones of the chaperonin and Hsp70 families (Hassell *et al*., 2022), presumably to maintain the excess Cdc12 in a conformation that is competent to properly associate with other septins once those molecules are synthesized. When those chaperones were mutated, excess Cdc12 persisted in cells but was unable to incorporate into septin filaments (Hassell *et al*., 2022). Notably, this model of chaperone-guided septin hetero-octamer assembly focused on the septin GTPase domain and ignored the CTD, in part because the yeast septin Cdc10 lacks a CTD but displays similar chaperone interactions to Cdc12 and the other CTD-containing septins (Johnson *et al*., 2015; Denney *et al*., 2021). Thus it was unknown if chaperones also engage otherwise unoccupied septin hetero-oligomerization interfaces within the CTDs to promote efficient biogenesis of functional septin complexes.

One abundant cytosolic chaperone in budding yeast, Hsp104, displayed septin interactions that were unique compared to other chaperones. Hsp104 forms a homohexamer that is capable of disassembling protein oligomers, including β aggregates, by threading substrate proteins through a central channel and thereby unfolding them (Shorter and Southworth, 2019). We previously reported that Hsp104 binds all budding yeast septins and shows evidence of increased binding to G-interface mutant septins (Johnson *et al*., 2015; Denney *et al*., 2021). Intriguingly, overexpression of a double-mutant Hsp104 (G217S T499I) causes lethality in wild-type yeast cells in a manner that mimics septin dysfunction: cells become highly elongated and fail in cytokinesis (Hartwell, 1971; Schirmer *et al*., 2004). Either of two spontaneous single missense mutations in the Cdc12 CTD is sufficient to confer resistance to the lethality of Hsp104(G217S T499I) (Schirmer *et al*., 2004). These observations suggested that wild-type Hsp104 may normally bind the wild-type Cdc12 CTD without adverse effect on Cdc12 function, whereas the G217S T499I mutations unleash Hsp104 unfolding activity upon Cdc12, preventing septin hetero-octamer and filament assembly. The Cdc12 mutations might confer resistance by inhibiting Hsp104(G217S T499I) binding. However, these speculations had not been tested.

Here we investigated homo-oligomerization of the Cdc12 CTD and the contexts in which it might be physiologically relevant. Substitutions in a residue in the Cdc12 CTD, E368, altered both homo-oligomerization and the effects of Hsp104(G217S T499I), leading us to further explore the relationship between Cdc12 oligomerization and Hsp104.

## RESULTS

### Coiled-coil-mediated homo-oligomerization of the Cdc12 C-terminal domain

Previous studies provided evidence of CTD-mediated Cdc12 homo-oligomerization. In a yeast two-hybrid assay of septin-septin interactions (which may be complicated by interactions with endogenous septins), Cdc12– Cdc12 interaction generated the strongest signal and required the CTD (Farkasovsky *et al*., 2005). Purified full-length Cdc12 was almost exclusively a homodimer when analyzed by size exclusion chromatography (SEC) (Hassell *et al*., 2022). The Cdc12 CTD (residues 339-407) fused to GST binds full-length Cdc12 in vitro, demonstrating that the CTD sequence alone is capable of mediating Cdc12 homo-oligomerization (Versele *et al*., 2004), and analysis of a small part of the Cdc12 CTD (residues 368-407) revealed evidence of homo-oligomeric coiled coils that could also weakly hetero-dimerize with the Cdc3 CTD (Barth *et al*., 2008). We analyzed the purified Cdc12 CTD by SEC-multi-angle light scattering (SEC-MALS) and found evidence of coiled-coil-mediated Cdc12 homodimerization and, unexpectedly, homotrimerization (Fig. 1 D). The Cdc12 CTD was either fused to a short residual N-terminal sequence (GSHM, in single-letter amino acid code) following cleavage with thrombin from a larger hexahistidine (H_6_) fusion, or fused at its N terminus to a H_6_ tag and *E. coli* maltose-binding protein (MBP). GSHM fused to Cdc12 residues 316-407 or 341-407 generated a slightly asymmetrical peak with predicted molecular mass 2.6 times that of the monomer (Fig. 1 D). We interpret these signals as a mix of homodimers and homotrimers undergoing rapid exchange relative to the light-scattering timescale. The indistinguishable behavior of the two constructs shows that the unstructured tail in the CTD (residues 316-340) does not contribute to the oligomeric equilibrium. We also directly detected a homotrimeric species when we crosslinked the shorter construct with bis(sulfosuccinimidyl)suberate (BS3) and resolved the products by tricine-SDS-PAGE (Fig. 1 E).

We used AlphaFold3 (Abramson *et al*., 2024) to predict the structure of the Cdc12 CTD homotrimer at higher resolution. The result was a parallel trimer with high-quality metrics (high average predicted local distance difference test (pLDDT) for the coiled coil region (77.3) and low interchain predicted aligned errors (pAEs), Fig. S1 A). Including the residual GSHM residues and/or removing residues 316-340 did not affect the outcome (Fig. 1 F, Fig .S1 B). To determine the coiled-coil orientation in solution, we employed SEC coupled with small angle X-ray scattering (SAXS) using N-terminal MBP-fused constructs to add mass to one end of each helix. All relevant information regarding the SEC-SAXS data analysis is presented in Table S1. We analyzed two distinct constructs that differed in the number of residues between the coiled-coil forming sequences and MBP: “MBP-Cdc12CC” has a three-residue linker (SGS) before Cdc12 residue 340, whereas in “MBP-Cdc12CC+8” the Cdc12 residues following the SGS linker begin at Cdc12 residue 332. Initially, frames corresponding to the top of the SEC elution peaks were analysed. Dimensionless Kratky plots indicate folded complexes (Figs. S2, S3). All values for MBP-Cdc12CC+8 were consistent with a mix of dimers and trimers, as with the isolated Cdc12 sequences. MBP-Cdc12CC, on the other hand, behaved as a relatively pure homodimer. We suspect that the shorter linker between the MBP and Cdc12 sequences in this construct led to interactions between the MBP portions that prevented trimerization.

Atomistic modelling of MBP-Cdc12CC data with MultiFoXs (Schneidman-Duhovny *et al*., 2016) using flexible linkers agreed with a parallel dimeric assembly (χ^2^= 1.41, CorMap *P*-value = 0.051). The data for MBP-Cdc12CC+8 could not be fitted well with either a single parallel dimer (di-p, χ^2^ = 2.17, CorMap *P*-value = 0.000) or a single parallel trimer (tri-p, χ^2^ = 7.91, CorMap *P*-value = 0.000). We then employed EOM-NNLSJOE (Bernadó *et al*., 2007; Tria *et al*., 2015) to analyse the MBP-Cdc12CC+8 dataset considering a mixture of parallel dimers and trimers. Antiparallel assemblies were not considered for the fit given the incompatibility between their calculated *R*_g_’s and the experimental *P*(*r*)-derived *R*_g_ (Fig. S4). RANCH was used to build 10,000 linker-flexible models, equally divided between parallel dimers and trimers (5,000 models each). EOM-NNLSJOE identified an ensemble in agreement with the experimental data (χ^2^ = 1.07, CorMap *P*-value = 0.203), in which 66.0% of the models are parallel trimers and 34.0% are parallel dimers.

An additional analysis was conducted on two parts of the MBP-Cdc12CC+8 elution peak, selecting groups of contiguous frames in which trimers (peak front) and dimers (peak tail) were likely enriched (Fig. S5). Indeed, the calculation of mass, *D*_max_ and Porod volume using these two datasets was consistent with this assumption. The *P*(*r*) function of the peak tail dataset resembled that obtained for MBP-Cdc12CC. Real space *R*_g_’s from the peak front and the peak tail datasets matched well with the mean/median of tri-p and di-p models generated by RANCH, respectively (Fig. S4). Atomistic model fitting with a single-state parallel dimer using MultiFoXs yields appropriate χ^2^ for both reduced regions (χ^2^= 1.28 and 1.20 for peak front and peak tail datasets, respectively) although with CorMap *P*-values (0.0057 and 0.0053, respectively) insufficient to assure similitude between experimental and calculated data. MultiFoXs analysis using a single-state parallel trimer generates inadequate fit for both regions. This might indicate that MBP-Cdc12CC+8 is flexible enough to prevent a good fit of its SAXS data using a single oligomer and conformer. Fit of the MBP-Cdc12CC+8 peak front dataset with EOM-NNLSJOE using parallel dimers and trimers results in a good fit with an ensemble composed of a majority of trimers (χ^2^ = 1.01, CorMap *P*-value = 0.940; 53.4% tri-p, 46.6% di-p). For MBP-Cdc12CC+8 peak tail dataset, parallel dimers and trimers also fit the data well, with only a small fraction of trimers (χ^2^ = 0.97, CorMap *P*-value = 0.310; 93.0% di-p, 7.0% tri-p). This confirmed our initial conjecture that MBP-Cdc12CC+8 exists as a mixture of dimers and trimers (Fig. 1 G). Together, these observations are consistent with an intrinsic ability of the Cdc12 CTD to form two kinds of homomeric parallel coiled coils in solution: homodimers and homotrimers.

### Mutations in a conserved motif alter Cdc12 homo-oligomerization in vitro

Previous studies identified a sequence motif enriched in homotrimeric coiled coils, RhxxhE, wherein “h” denotes a hydrophobic residue and “x” is any residue (Kammerer *et al*., 2005). A perfect match to this motif is found in the middle of the Cdc12 CTD (RFRQWE), representing the most highly conserved stretch of CTD residues (Fig. 1 B). W367 is conserved in most Cdc12 homologs and its presence in the core of the predicted homotrimer stands out, interacting hydrophobically with the two other Trp from other chains and also with F364. In the prediction, W367 and E368 side chains from different helices are in close proximity (Fig. 2 A), and residues within the core pack against each other with acute angles, as expected for coiled-coil trimers. The coiled-coil radius reaches its maximum value of ∼7.7 Å near the RhxxhE motif, likely to accommodate bulky side-chain residues (F364 and W367) in the core, and gradually decreases to a typical value (∼6.0 Å) towards the C-terminus. The enlarged Cdc12 coiled-coil radius, which creates cavities in the region between F353 and W367, likely underlies its metastability.

**Figure 2.**
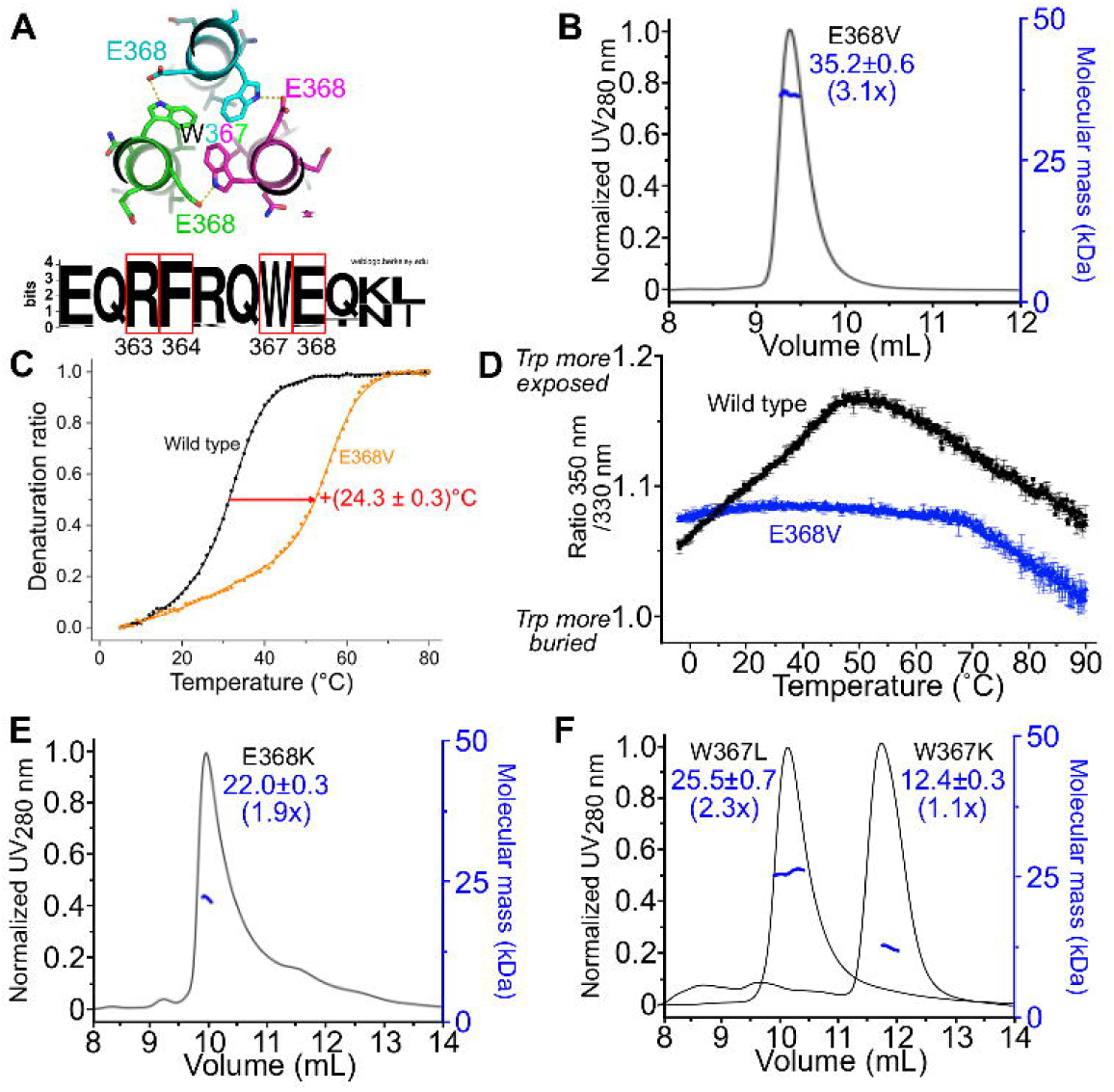
Mutations in a conserved homotrimerization motif within the Cdc12 C-terminal domain alter coiled-coil oligomerization. (A) The W367 environment within the predicted Cdc12 CTD homotrimeric coiled coil, and sequence logo of the neighbouring residues, highlighting R363, F364, W367 and E368, the residues that match the RhxxhE homotrimerization motif. (B) As in Fig. 1 D, SEC-MALS analysis of purified Cdc12 CTD with the E368V mutation. (C) Circular dichroism denaturation transitions monitored at 222 nm of wild type (black) or E368V mutant (orange) Cdc12 CTD at 10 µM. The red arrow indicates the difference in temperature for which 50% of the protein was denatured. (D) NanoDSF analysis of the intrinsic fluorescence intensity ratio (350/330 nm) of wild type (black) or E368V mutant (blue) at 90 µM over the indicated temperature range during unfolding. Error bars represent standard deviation from duplicate measurements. As W367 is the sole tryptophan in Cdc12 CTD and there are only two tyrosines, the signals acquired predominantly report on W367. (E-F) As in (B), SEC-MALS analysis of purified Cdc12 CTD with the indicated mutations.

We identified a single substitution in the homotrimerization motif, E368V, that drives the normal CTD homodimer:homotrimer equilibrium to homotrimers exclusively (Fig. 2 B). The E368V homotrimers were exceedingly thermostable (Fig. 2 C, Table S2). In the AlphaFold model, the side chain of E368 projects slightly away from the hydrophobic core of the trimer that includes W367 and F364 (Fig. 2 A); burial of the hydrophobic valine side chain in this hydrophobic core likely confers the high stability of the E368V homotrimer. Consistent with the prediction, nano differential scanning fluorimetry experiments indicated that W367 is more buried when E368 is replaced by valine (Fig. 2 D). Replacing E368 with lysine (E368K), a residue found in this position in the Cdc12 homolog from the yeast *Ogataea parapolymorpha*, should alter the contact with W367 without disturbing the hydrophobic core, and indeed this mutation maintained helical character while shifting the equilibrium to homodimers (Fig. 2 E, Table S4). Disturbing the hydrophobic core by replacing W367 with hydrophilic lysine (W367K) ablated oligomerization and reduced helical character (Tables S2-4), whereas substitution to another hydrophobic residue, leucine, did not (Fig. 2 F). These results support a model in which the CTD of Cdc12 mediates metastable coiled-coil homo-oligomerization via a previously unrecognized, highly conserved motif, mutations in which can drive homotrimerization or block homo-oligomerization altogether.

### Evidence for homo-oligomerization by Cdc12 overexpressed in yeast cells

In the top-ranked AlphaFold3 prediction for a full-length Cdc12 trimer, homotrimerization was mediated by the same three-helix coiled coil predicted by our in silico and in vitro experiments with the isolated CTDs (Fig. 3 A). [We note that in this trimeric context the globular Cdc12 GTPase domains did not interact with each other. By contrast, in all five predictions for a full-length Cdc12 homodimer, the globular GTPase domains interacted via the canonical septin G or NC interface (Fig. S6), even though such Cdc12–Cdc12 interfaces are not found in the native septin hetero-octamer (Fig. 1 A).] To ask if such Cdc12 homo-oligomers can occur in living cells, we used the *GAL1/10* promoter and culture medium containing 2% galactose to drive overexpression of C-terminally GFP-tagged Cdc12, lysed cells, separated clarified lysates by SEC, and used an anti-GFP antibody to detect relative Cdc12-GFP levels in fractions corresponding to expected sizes of hetero-octamers, homotrimers, and homodimers. Whereas in past studies we used high-salt SEC buffers to disassemble septin filaments in yeast lysates (Johnson *et al*., 2015), here we used physiological salt (150 mM) to avoid potential effects on coiled-coil stability. GFP signal was found in three apparent peaks: mostly in the hetero-octamer fraction (expected ∼400 kDa, observed 334-391 kDa), less in a homotrimer-sized fraction (expected ∼210 kDa, observed 208-244 kDa) and even less in a homodimer-sized fraction (expected ∼144 kDa, observed 130-152 kDa) (Fig. 3 B,C). While SDS-PAGE and immunoblot analysis confirmed the presence of full-length Cdc12-GFP in each of these fractions, the homodimer-sized fraction also contained a prominent ∼30-kDa band of unknown origin recognized by the GFP antibody (Fig. S7). However, as we observed in vitro with purified Cdc12 CTD, the relative abundance of apparent homotrimers increased, and the putative homodimers decreased, upon introduction of the E368V mutation (Fig. 3 B). We thus suspect that the ∼30-kDa band reflects post-fractionation proteolysis of full-length Cdc12-GFP homodimers. Due to its severe effects on cell proliferation (see below), we did not assess the W367K mutant in the same way. While they do not address the presence or absence of other proteins in complex with the overexpressed Cdc12-GFP, these data are consistent with the accumulation of excess molecules of Cdc12 in CTD-mediated homodimers and -trimers in vivo.

**Figure 3.**
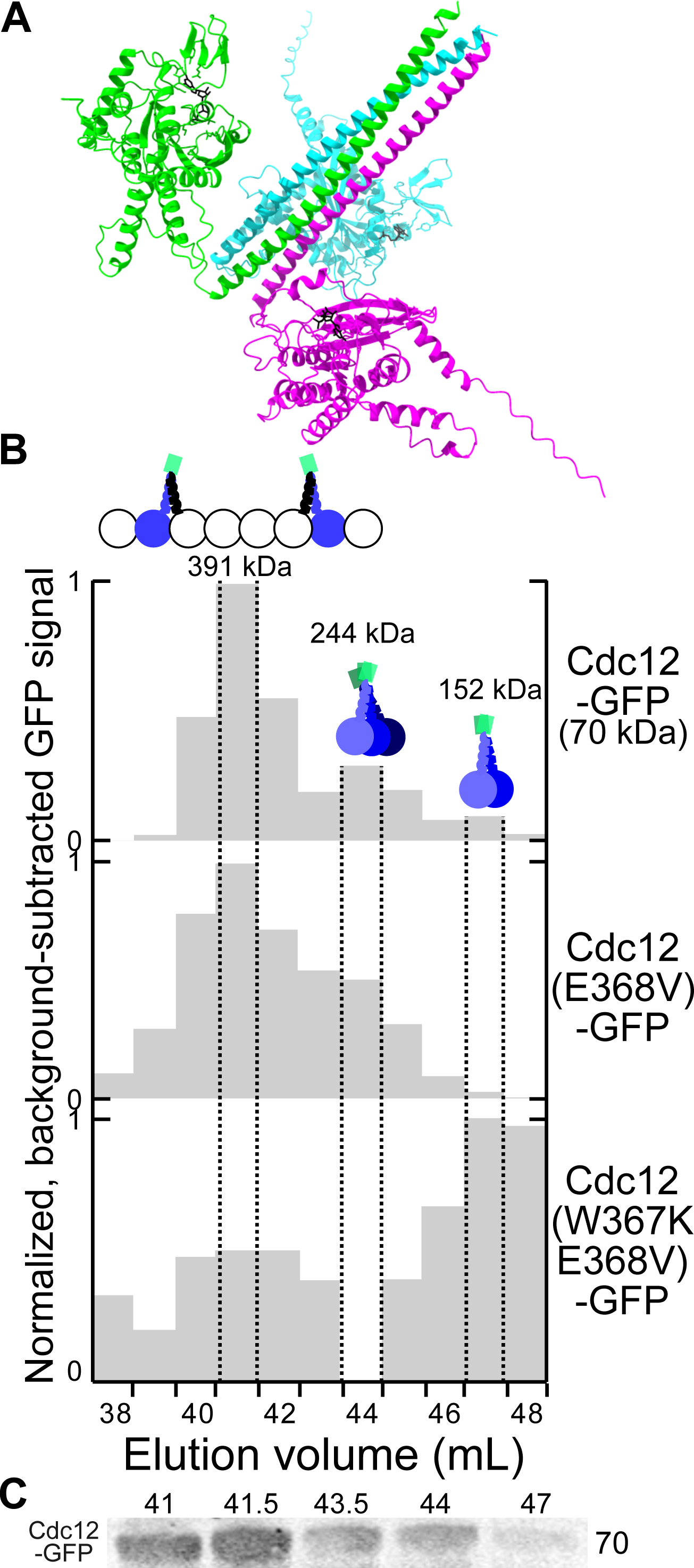
Homo-oligomerization by overexpressed Cdc12 molecules in yeast lysates. (A) Top-ranked AlphaFold3 prediction for homotrimer of full-length Cdc12 bound to GDP. (B) SEC analysis of lysates of wild-type yeast cells (strain BY4741) overexpressing the indicated version of C-terminally GFP-tagged Cdc12. Plotted are the relative amount of Cdc12-GFP in each 1-mL elution fraction as determined by dot blotting and detection with anti-GFP antibody and fluorescently labeled anti-GFP antibodies. For each dot, fluorescence was normalized to the dot with the highest fluorescence intensity. Background fluorescence was then corrected by subtracting the normalized signal of the dot with the lowest fluorescence intensity from all normalized signals. The dashed lines outline fractions corresponding to the three apparent peaks; values given above each peak indicate the largest size in kDa of molecules expected in that fraction, based on calibration of the column with molecular weight standards (Fig. S7 A). Cartoons indicate the Cdc12-containing septin complex that each peak likely represents, based on a monomeric Cdc12-GFP size of 70 kDa, with the largest complex being a septin hetero-octamer. Green squares represent GFP. Plasmids were pMVB2 (“Cdc12-GFP”), BD7C264E (“Cdc12(E368V)-GFP”), and 9F2B6E98 (“Cdc12(W367K E368V)-GFP”). (C) The experiment with wild-type Cdc12-GFP was repeated but 0.5-mL fractions were collected and the protein was precipitated and resolved by SDS-PAGE prior to immunoblotting. Fractions corresponding to the indicated elution volumes are shown, with position of 70-kDa molecular weight standard indicated at right; additional fractions and uncropped blots are shown in Fig. S7.

We wondered if the increased homotrimer stability induced by the E368V mutation might “lock” the CTDs of excess Cdc12 subunits into permanent homotrimers that would be unable to incorporate into septin hetero-oligomers. However, excess molecules of Cdc12(E368V)-GFP showed no defect in post-translational filament assembly (Fig. S8), suggesting that even the E368V-mutant Cdc12 homo-oligomers are sufficiently metastable to eventually hetero-oligomerize.

### Cdc12 C-terminal domain homo-oligomerization mutants exert potent dominant-negative effects on septin filament function

To further study potential roles for Cdc12 CTD-mediated homo-oligomerization in septin hetero-oligomerization and function in vivo, we overexpressed GFP-tagged Cdc12 or W367K or E368V mutants thereof in haploid cells co-expressing wild-type, untagged Cdc12 and mCherry-tagged Cdc3 from the respective genomic loci. We compared colony growth, cell morphology, and septin localization of cells cultured in galactose-containing medium to control conditions with 2% glucose, which represses *GAL1/10* promoter activity. Surprisingly, wild-type Cdc12-GFP overexpression slightly reduced colony growth (Fig. S9 A), and cells were frequently elongated and displayed mislocalization of mCherry-Cdc3, a phenotype that was not previously observed using this same Cdc12-GFP plasmid in cells of the same strain background but lacking any tag on Cdc3 (Versele *et al*., 2004; Benson and McMurray, 2023). The mCherry-Cdc3 fusion we used has also never before been reported to itself cause septin dysfunction (Gao *et al*., 2007; McMurray, 2016). We suspected that, since Cdc3 and Cdc12 directly interact via an interface involving residues near their N and C termini (Fig. 1 A), the defects we observed represent a “synthetic” effect of the C-terminal GFP tag on Cdc12 combined with the mCherry tag inserted near the N terminus of Cdc3.

To address this concern, we repeated these experiments in yeast cells with no tag on any other septin except the GFP tag on the overexpressed Cdc12 (the same set-up used in the SEC experiments above). Here, only the mutant versions of Cdc12-GFP had any detectable effect on colony growth and cell morphology, and these effects were equivalent to those seen in the strain with tagged Cdc3 (Fig. 4 A and Fig. S9 A). Thus, the effects of CTD mutations in Cdc12 were not a consequence of Cdc3 dysfunction caused by the mCherry tag. Whereas the E368V mutant had no obvious effect, the W367K mutant was nearly lethal upon overexpression, causing drastic morphology and cytokinesis defects (and mCherry-Cdc3 mislocalization; Fig. 4 and S9 A,B). We also tested a W367K E368V double mutant construct, which caused defects but not as severe as W367K alone (Fig. S9 A,B). SEC results suggested that Cdc12(W367K E368V) molecules undergo some degree of homo-oligomerization in vivo and in vitro (Fig. 3 B, Fig. S9 C); the destabilizing effect of the hydrophilic lysine at position 367 is likely counteracted by the hydrophobic valine at position 368.

**Figure 4.**
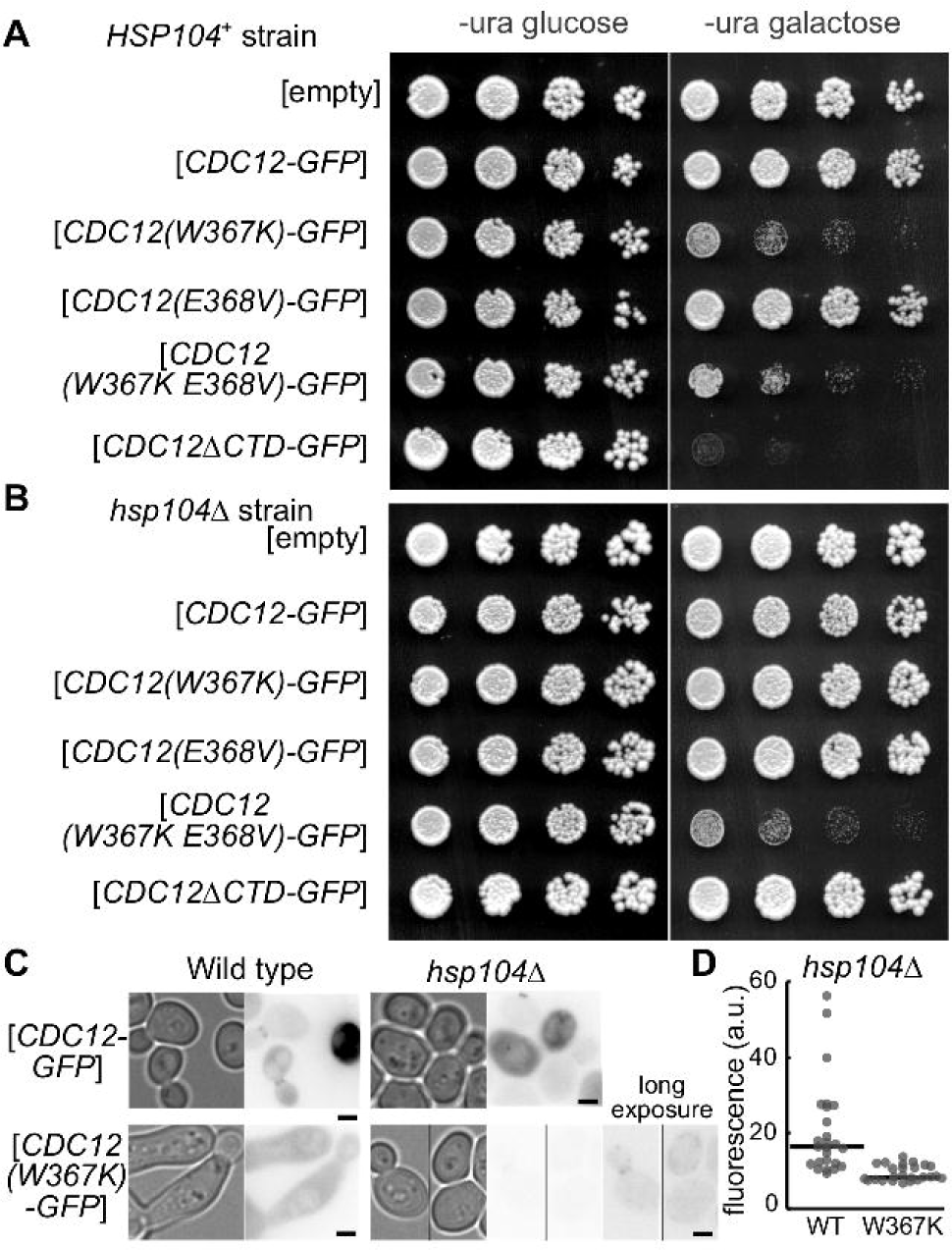
Dominant inhibition of septin function by C-terminal domain homo-oligomerization mutants of the septin Cdc12. (A) Five-fold serial dilutions of yeast cells (strain BY4741) were spotted on medium selective for the indicated *URA3*-marked plasmid and either repressing (“-ura glucose”) or inducing (“-ura galactose”) expression of the indicated version of GFP-tagged Cdc12 or the empty vector pRS316. *CDC12* plasmids were pMVB2 (“*CDC12-GFP*”), C6F1B99E (“*CDC12(W367K)-GFP*”), (BD7C264E (“*CDC12(E368V)-GFP*”), 9F2B6E98 (“*CDC12(W367K E368V)-GFP*”), and pMVB160 (“*CDC12*Δ*CTD-GFP*”). Plates were imaged after 2 days incubation at 30°C. (B) As in (A) but with the *hsp104*Δ yeast strain JTY4013. (C) Micrographs of representative cells scraped from the galactose plates in (A) and (B) and imaged by transmitted light or the GFP cube (60-msec exposure). “Long exposure” indicates a 250-msec exposure. Scale bars, 2 µm. (D) Fluorescence in a 10-pixel-diameter circular area was quantified for 25 cells each from the cultures as shown in (C) of *hsp104*Δ yeast strain JTY4013 carrying either the plasmid pMVB2 (“WT”) or C6F1B99E (“W367K”). “a.u.”, arbitrary units; lines indicate medians.

We also included a truncated version of Cdc12-GFP lacking the CTD (residues 339-407 deleted), which was already known to be dominant lethal when overexpressed (Johnson *et al*., 2015). The W367K mutant was nearly as potently lethal as the ΔCTD mutant (Fig. 4 A). We interpret the ability of the Cdc12 mutants to interfere with septin function despite the presence of wild-type Cdc12 as evidence that the mutant proteins co-assemble with wild-type septins into dysfunctional complexes/filaments.

When we determined the sequence of the mutant Cdc12-GFP plasmids to confirm the presence of the mutations, we discovered another mutation already present in the “wild-type” Cdc12-GFP parent plasmid, encoding the substitution N379D. N379 is highly conserved and makes a predicted contact in the Cdc12 CTD homotrimer but not the Cdc3–Cdc12 heterodimer (Fig. 1 B). It is possible that N379D contributes to the septin dysfunction we saw in cells with mCherry-tagged Cdc3 overexpressing Cdc12-GFP. However, restoring N379 in the W367K plasmid did not alter its ability to dominantly perturb septin function (Fig. S10 A). Furthermore, introducing the W367K mutation into the endogenous *CDC12* gene in haploid cells with mCherry-tagged Cdc3 confirmed that this substitution alone is sufficient to confer severe proliferation, morphology, and septin localization defects (Fig. S10 B). Thus we consider the N379D mutation to be irrelevant with regard to the effects of the CTD mutations we deliberately made.

### Mutating the Cdc12 CTD homotrimerization motif mimics the effects of CTD truncations on cellular septin functions

According to our model, Cdc12(W367K) dominantly perturbs septin function not because excess Cdc12 monomers are unable to homo-oligomerize, but because Cdc12(W367K) monomers are able to co-assemble with other septins into hetero-octamers, and the filaments made from these octamers are ultimately dysfunctional. Since W367 is located in the CTD, it might perturb septin function in a way similar to other CTD mutations, many of which cause phenotypes that are modified by temperature. For example, truncation of residues 392-407 (associated with the *cdc12-6* allele) is tolerated at 22°C but is lethal at ≥30°C (Adams and Pringle, 1984; Johnson *et al*., 2015). [Like the W367K mutant, the Δ392-407 truncation blocked homo-oligomerization by purified Cdc12 CTD (Table S4)]. We asked if the septin dysfunction caused by Cdc12(W367K) overexpression is also modified by temperature. Indeed, cultivation at 37°C exacerbated, and cultivation at 22°C ameliorated, the adverse effects of Cdc12(W367K), as well as the effects of the other CTD mutants, including Cdc12ΔCTD (Fig. 5 A, Fig. S10 B). Thus Cdc12(W367K) phenocopies the effects of other CTD mutations on septin function.

**Figure 5.**
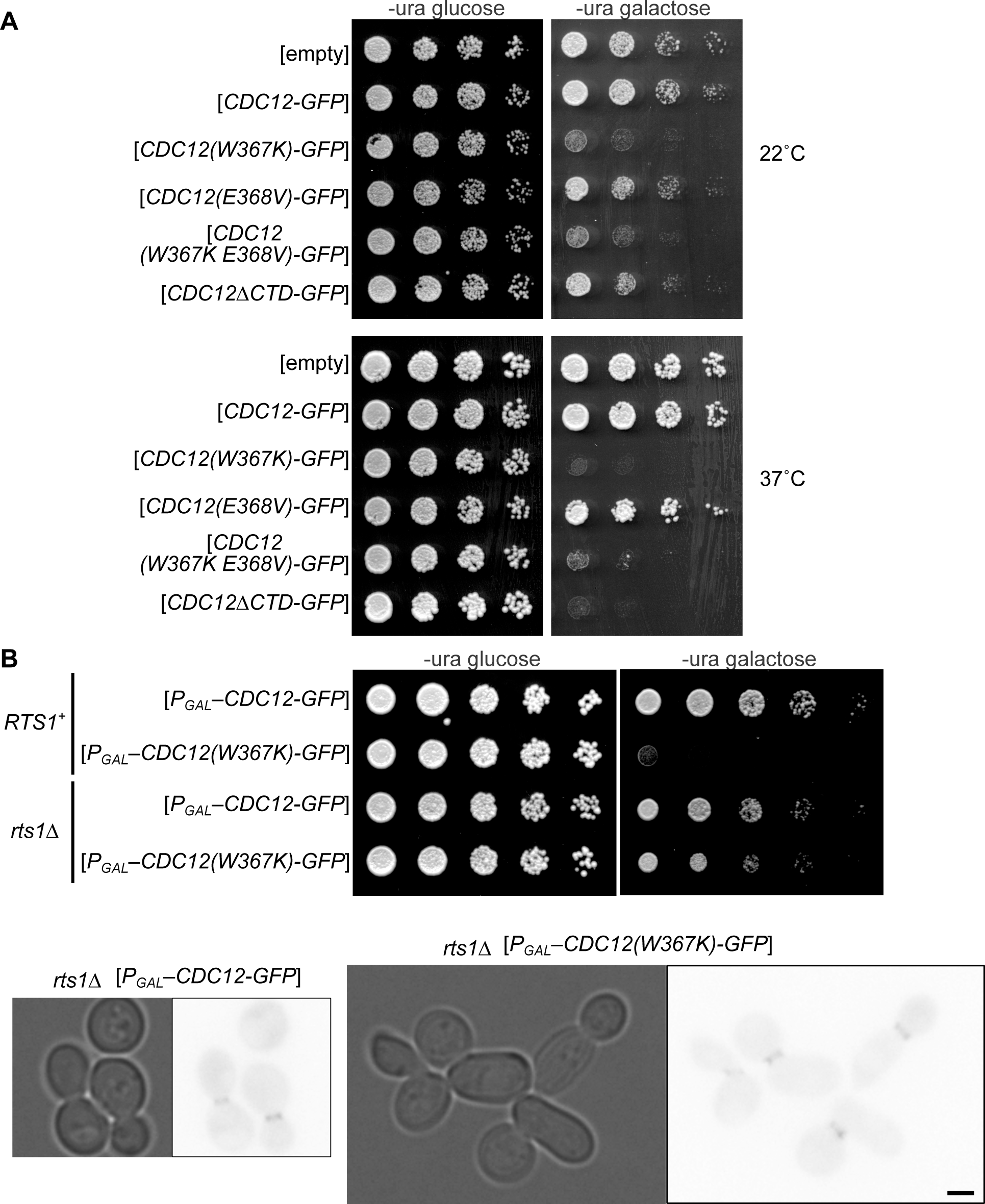
The PP2A phosphatase subunit Rts1 is required for dominant inhibition of septin function by C-terminal domain homo-oligomerization mutants of the septin Cdc12. (A) As in Fig. 4 A but at the indicated temperatures. (B) Top, as in (A), dilution series of wild-type (strain BY4741) or *rts1*Δ (strain H06957) cells carrying plasmids encoding the indicated Cdc12-GFP fusion under control of the galactose-inducible *GAL1/10* promoter. Plasmids were pMVB2 (“*CDC12-GFP*”) or C6F1B99E (“*CDC12(W367K)-GFP*”). Bottom, cells were scraped from colonies on the galactose plates in (A), suspended in water, and imaged by transmitted light or the GFP cube (60-msec exposure). Scale bar is 2 µm.

Deletion of the *RTS1* gene allows *cdc12-6* cells to proliferate at 30°C (Dobbelaere *et al*., 2003). *RTS1* encodes a phosphatase that drives destabilization of septin filaments and higher-order structural transitions during the yeast cell division cycle (Dobbelaere *et al*., 2003). If Cdc12(W367K) overexpression perturbs septin function by populating hetero-octamers with CTD-defective subunits, then *rts1*Δ might also confer the ability to proliferate at 30°C despite Cdc12(W367K) overexpression. Indeed, *rts1*Δ cells overexpressing Cdc12(W367K)-GFP were able to form colonies almost as well as cells overexpressing wild-type Cdc12-GFP, and showed only subtle morphological defects (Fig. 5 B). Furthermore, in *rts1*Δ cells Cdc12(W367K)-GFP localized strongly to septin rings at bud necks (Fig. 5 B). These findings support the conclusion that the W367K mutation does not perturb septin hetero-octamer assembly per se but the resulting octamers are defective in Cdc12 CTD function in a manner that resembles dysfunction associated with loss/truncation of the CTD. Moreover, septin filaments populated with Cdc12(W367K) subunits are severely dysfunctional only in the presence of Rts1; otherwise, they function almost normally.

### Substitutions of Cdc12 CTD residue E368 confer resistance to the lethal effect of Hsp104(G217S T499I)

Either of two single missense mutations in Cdc12 confers resistance to the lethality caused by overexpression of Hsp104(G217S T499I) (Schirmer *et al*., 2004). One of these mutations, E368Q, alters a residue we implicated in homotrimerization. We used AlphaFold3 to predict where the substrate-binding domain of Hsp104 (residues 1-152) binds Cdc12. First, as a control, we generated predictions of Hsp104 binding to residues 96-151 of a known Hsp104 substrate, the prion-forming protein Sup35 (Sweeny *et al*., 2015). Satisfyingly, all five predictions showed Hsp104 binding to precisely the region of Sup35 found recently by others using purified proteins (Shen *et al*., 2024) (Fig. 6 A, Fig. S11 A).

**Figure 6.**
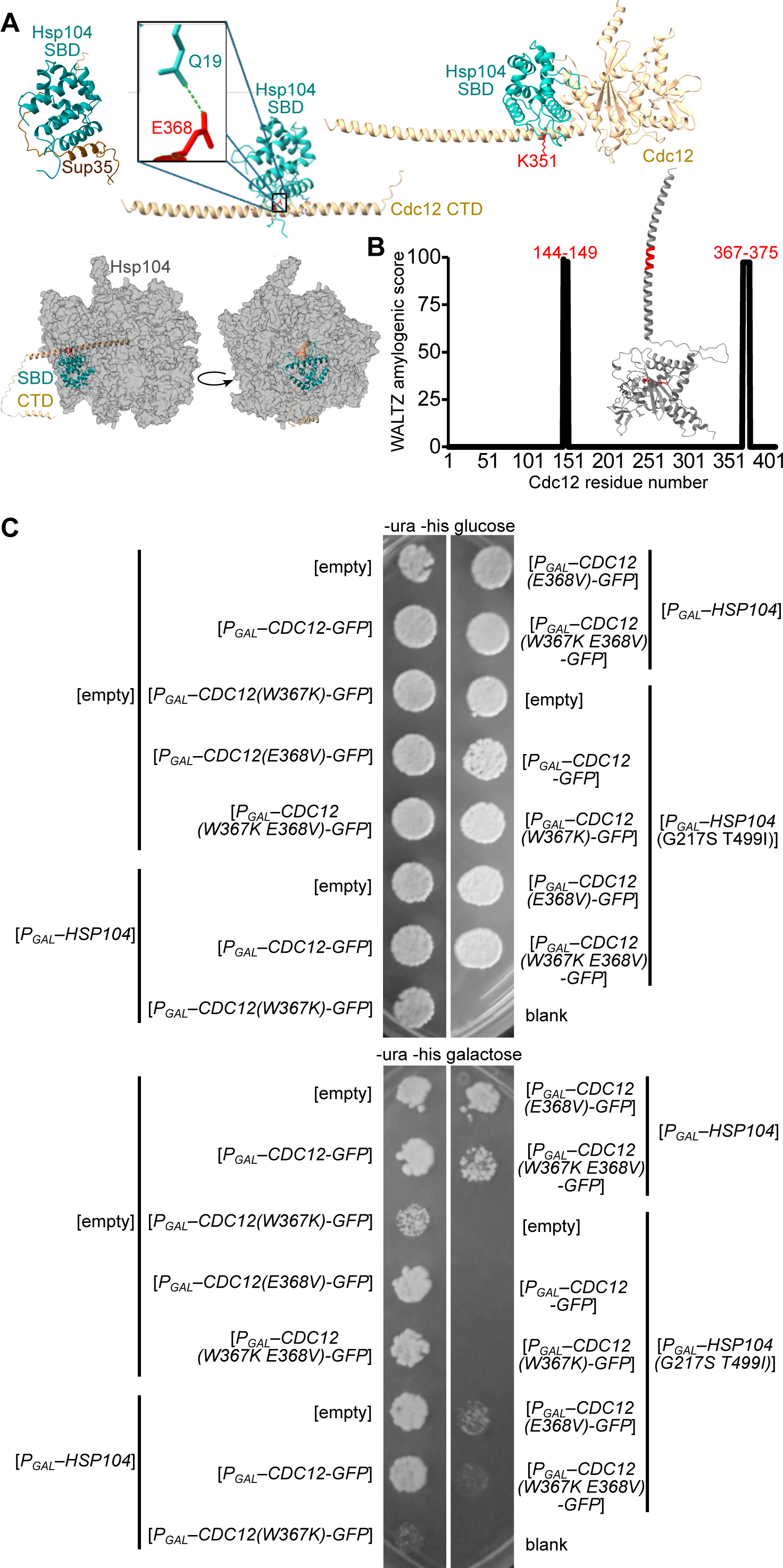
Evidence for direct Hsp104 binding to E368 in the Cdc12 C-terminal domain. (A) Top-ranked AlphaFold3 predictions for binding of the substrate-binding domain (“SBD”) of Hsp104 (residues 1-152) to the known substrate Sup35 (residues 96-151), the Cdc12 CTD (residues 297-407), or full-length Cdc12. Below, the complex of Hsp104 homohexamer bound to Cdc12 CTD was predicted by structurally aligning the Hsp104 SBD–Cdc12 CTD prediction to chain B of a closed-conformation Hsp104 cryo-EM structure (PDB 6N8T). (B) Left, WALTZ amyloidogenic scores plotted against the sequence of Cdc12. Right, the two regions with high WALTZ scores are colored in red in the top-ranked AlphaFold3 prediction for GDP-bound full-length Cdc12. (C) Serial dilutions were performed as in Fig. 4 A of wild-type cells (strain BY4741) on glucose or galactose medium selective for *HIS3*- and *URA3*-marked plasmids, either empty vectors (“[empty]”) or encoding the indicated alleles of *HSP104* and *CDC12*, incubated at 30°C for 2 days before imaging. For the sake of space, only the most concentrated spot in each dilution series is shown. Empty vectors were pRS313 and pRS316, other plasmids were 5679 (“*P_GAL_–HSP104*”), 5707 (“*P_GAL_–HSP104(G217S T499I)*”), pMVB2 (“*P_GAL_–CDC12-GFP*”), C6F1B99E (“*P_GAL_–CDC12(W367K)-GFP*”), (BD7C264E (“*P_GAL_–CDC12(E368V)-GFP*”), and 9F2B6E98 (“*P_GAL_–CDC12(W367K E368V)-GFP*”). No cells were spotted in the “blank” row.

Next, we generated predictions for Hsp104 binding to Cdc12, either the CTD alone (residues 297-407) or full-length Cdc12. In both cases, in the highest-confidence predictions Hsp104 contacts Cdc12 via residues that lie within the hydrophobic groove between helix 1 and helix 4 and overlap with residues that contact Sup35 and other known Hsp104 substrates (Harari *et al*., 2022) (Fig. 6 A). Hsp104 binding to the Cdc12 CTD centered around Cdc12 E368, and the side chain of Cdc12 E368 made predicted contact with Hsp104 Q19 (Fig. 6 A). When we used full-length Cdc12, Hsp104 binding centered around K351, with additional residues outside the Hsp104 hydrophobic groove contacting the globular GTPase domain of Cdc12 (Fig. 6 A). K351N is the other mutation that confers Hsp104(G217S T499I) resistance (Schirmer *et al*., 2004).

When we superimposed the Hsp104 homohexamer structure (PDB 6N8T) on the prediction for the Hsp104 substrate-binding domain bound to the Cdc12 CTD, the Cdc12 CTD projected directly into the central pore (Fig. 6 A). These predictions support a model in which Hsp104(G217S T499I) binds the Cdc12 CTD in two locations that are primed for threading and Cdc12 unfolding. Hsp104 threading can disassemble otherwise irreversible amyloid-type aggregates formed from β-sheet-like intermolecular interactions (Shorter and Lindquist, 2004). We previously noted that a sequence-based algorithm predicts high β-aggregation potential for the β4 strand in the septin GTPase domain that is ultimately buried in the septin-septin G interface, and we directly crosslinked chaperones to the Cdc10 β4 strand in vivo (Hassell *et al*., 2022). Indeed, in all AlphaFold3 predictions, the Hsp104 substrate-binding domain bound the aggregation-prone β4 peptides from Cdc10 and Cdc12 via the same interface as it binds Sup35 (Fig. S11 B). Whereas the Cdc12 CTD is predicted to be natively helical, the WALTZ algorithm (Maurer-Stroh *et al*., 2010) predicted residues 367-375 as the only non-β4 region with β-aggregation potential (Fig. 6 B). These findings are consistent with a model in which Hsp104 binds aggregation-prone regions in septin proteins, which includes a part of the Cdc12 CTD also involved in homo-oligomerization.

Notably, when wild-type Hsp104 takes apart β aggregates of Sup35, it does not unfold all of Sup35. Instead, Hsp104 threads only the Sup35 N-terminal tail, and leaves the C-terminal GTPase domain intact (Sweeny *et al*., 2015). This partial substrate processing behavior of Hsp104 has also been observed with other substrates (Haslberger *et al*., 2008). Thus our unbiased in silico predictions provide strong support for a model in which –– due to two misfolding/aggregation-prone regions, one in the GTPase domain and one in the CTD –– the Cdc12 CTD is a native substrate for folding assistance and/or aggregation reversal by wild-type Hsp104, and these events need not unfold the Cdc12 GTPase domain. The G217S T499I mutations in Hsp104 hyperactivate unfoldase activity, driving complete unfolding of the Cdc12 and resulting in defective septin filament assembly and cytokinesis failure. In this model, the E368Q mutation in Cdc12 prevents Hsp104(G217S T499I) binding to the CTD, which is frequently exposed due to coiled-coil metastability; the aggregation-prone region in the GTPase domain would be buried in the Cdc12–Cdc11 G interface and consequently hidden from Hsp104(G217S T499I).

One prediction of this model is that the E368V substitution should, like E368Q, confer protection from Hsp104(G217S T499I) overexpression. We co-overexpressed in the same cells Hsp104(G217S T499I) and Cdc12-GFP or CTD mutants thereof and assessed colony growth via dilution series. Consistent with the prediction, overexpression of the Cdc12(E368V) allowed colony growth by cells also overexpressing Hsp104(G217S T499I) (Fig. 6 C). Morphology of cells from these colonies and bud neck localization of Cdc12-GFP therein correlated with colony growth (Fig. S12 A). Overexpression of wild-type Hsp104 had no obvious effect on colony growth, cell morphology, or Cdc12-GFP localization (Fig. 6 C and Fig. S12 A).

These results support a model in which Hsp104 binds the Cdc12 CTD unless a CTD mutation (E368V) that promotes homotrimerization is present. We imagined two ways this could work: the mutation could directly eliminate a critical contact between Hsp104 and Cdc12, or the mutation could drive homotrimerization, which buries the residue(s) otherwise involved in Hsp104–Cdc12 contact. To distinguish between these two models, we asked if the E368Q mutation also promotes CTD homotrimerization in vitro. Unlike E368V, and instead like the wild type, purified Cdc12(E368Q) CTD formed a mix of homodimers and trimers (Table S4). Thus we favor the interpretation that mutations at E368 confer resistance to overexpression of Hsp104(G217S T499I) due to loss of direct contact between Hsp104 and E368, not as a result of burial of Hsp104–Cdc12 contacts specifically in a coiled-coil homotrimer.

### Hsp104 potentiates the dominant-negative effects of homo-oligomerization-defective Cdc12(W367K) on septin assembly by protecting CTD-mutant Cdc12 against proteasomal degradation

The findings above suggest that Hsp104 binds the Cdc12 CTD and that mutations affecting CTD-mediated Cdc12 homo-oligomerization also affect interaction with Hsp104. We therefore asked if deleting *HSP104* alters the phenotypic consequences of overexpressing CTD-mutant versions of Cdc12. Strikingly, the absence of Hsp104 completely eliminated the functional defects associated with overexpression of Cdc12(W367K)-GFP but had no effect on the phenotypes associated with overexpression of Cdc12(W367K E368V)-GFP (Fig. 4 B, C). The amelioration of Cdc12(W367K)-induced septin dysfunction in *hsp104*Δ cells was accompanied by a drastic decrease in the cellular levels of Cdc12(W367K)-GFP (Fig. 4 C). By contrast, wild-type Cdc12-GFP was unaffected by deletion of *HSP104* (Fig. 4 C). Quantification of cytoplasmic GFP signal in *hsp104*Δ cells confirmed that Cdc12(W367K)-GFP levels were significantly lower than wild-type Cdc12-GFP (Mann Whitney U test *P* = 7.9×10^−7^, Fig. 4 D). The simplest explanation for the amelioration of septin dysfunction is that the absence of Hsp104 specifically prevented high-level accumulation of Cdc12(W367K)-GFP, and therefore that Hsp104 is required for such accumulation.

Hsp104 could promote high-level accumulation of Cdc12(W367K) by preventing degradation of monomeric Cdc12. According to this model, the effects of Cdc12(W367K E368V) overexpression are unchanged by *hsp104*Δ because the E368V mutation restores Cdc12 homo-oligomerization, not because the E368V mutation prevents Hsp104 interaction with the Cdc12 CTD. To learn more about how changes to the Cdc12 CTD influence the dependency on Hsp104, we tested the effect of *hsp104*Δ on overexpression of Cdc12ΔCTD-GFP. As with Cdc12(W367K)-GFP, cells lacking Hsp104 were entirely unaffected by the presence of the Cdc12ΔCTD-GFP-encoding plasmid (Fig. 4 B). Notably, faint Cdc12(W367K)-GFP signals could be detected in septin rings at the bud necks of *hsp104*Δ cells (Fig. 4 C). Thus we found Cdc12(W367K) localization to the bud neck in two independent scenarios: in *rts1*Δ cells (Fig. 5 B), where its adverse effects on septin filament function are seemingly neutralized, and in *hsp104*Δ cells, where the presence of a greater proportion of wild-type (untagged) Cdc12 in rods/filaments apparently suffices for function.

Human Septin-5 is degraded by the proteasome (Zhang *et al*., 2000). To ask if the proteasome is responsible for the lack of high-level accumulation of Cdc12(W367K) in *hsp104*Δ cells, we combined *hsp104*Δ with a hypomorphic allele of a proteasome subunit, *pre1(S142F)*, which reduces the proteolytic activity of the proteasome (Heinemeyer *et al*., 1991; Seufert and Jentsch, 1992)). In these cells, overexpressed Cdc12(W367K)-GFP accumulated to sufficiently high levels to perturb septin function (Fig. 7 A, B), representing a restoration of the behavior observed in *HSP104^+^* cells. Quantification of cytoplasmic GFP signal confirmed that in these cells Cdc12(W367K)-GFP levels were not significantly different than wild-type Cdc12-GFP (Mann Whitney U test *P* = 0.09, Fig. 7 C). We conclude from these data that Hsp104 protects excess Cdc12 monomers from proteasomal degradation. In the absence of Hsp104, the excess mutant Cdc12 molecules stabilized by the proteasome mutation must not all be irreversibly misfolded or aggregated –– at least in the conditions tested here –– because at least some are apparently competent to interact with other septins and dominantly perturb septin function.

**Figure 7.**
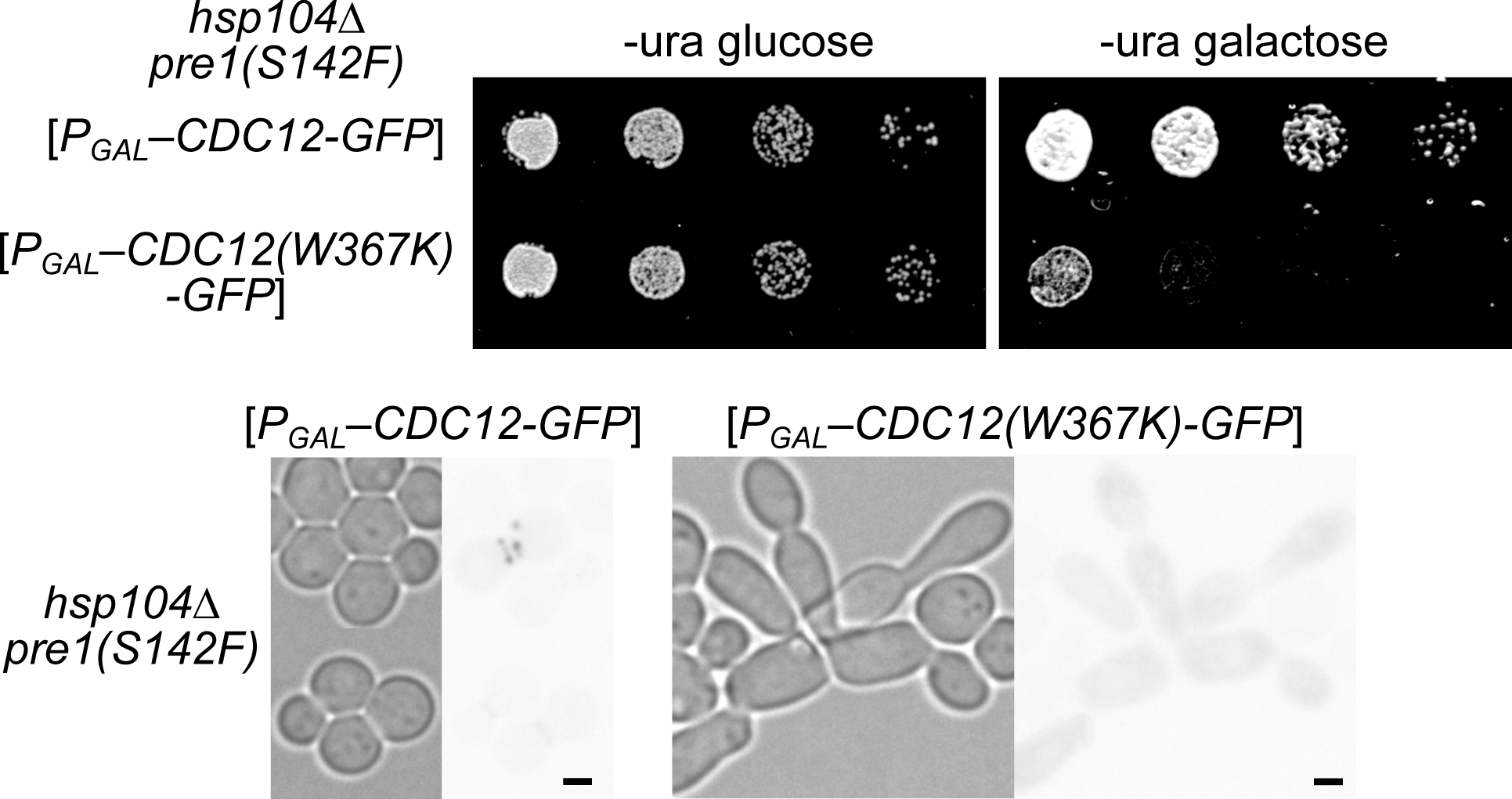
Hsp104 protects monomeric Cdc12 against proteasomal degradation. (A) Dilution series as in Fig. 4 A with the *hsp104*Δ *pre1(S142F)* yeast strain F93544A5 carrying plasmids pMVB2 (“*CDC12-GFP*”) or C6F1B99E (“*CDC12(W367K)-GFP*”). (B) Micrographs of representative cells scraped from the galactose plate and imaged with transmitted light or the GFP cube (250-msec exposures). Scale bars, 2 µm. (C) Fluorescence in a 10-pixel-diameter circular area of the cytoplasm was quantified for 25 cells each from the cells imaged as in (B). “a.u.”, arbitrary units; lines indicate medians.

### Hsp104 co-localization and co-elution with Cdc12 CTD mutants

For Hsp104 substrates that form visible foci in cells, Hsp104 co-localizes with those foci (Lum *et al*., 2004). We previously showed that the same construct we used here for overexpressing Cdc12-GFP (but not other constructs with different Cdc12 fusions) leads to the formation of foci in a subset of cells, and these foci rarely co-localize with mCherry-tagged Hsp104 (Hassell *et al*., 2022). We exploited the focus formation by GFP-tagged Cdc12 to test our model that Hsp104 stabilizes monomeric Cdc12 via direct binding. Whereas most cells overexpressing Cdc12(W367K)-GFP become grossly misshapen, a subset of cells have relatively normal morphology, allowing us to compare Hsp104-mCherry–Cdc12-GFP co-localization between cells overexpressing wild-type Cdc12-GFP and cells overexpressing Cdc12(W367K)-GFP. The W367K mutation had no effect on the frequency of Cdc12-GFP foci but caused a >20-fold increase in the co-localization of Cdc12-GFP and Hsp104-mCherry in foci: among 340 cells expressing wild-type Cdc12-GFP, there were 183 GFP foci and 139 mCherry foci, of which only 2 overlapped (0.6%), whereas among 197 cells expressing Cdc12(W367K)-GFP, there were 113 GFP foci and 107 mCherry foci, of which 27 overlapped (13.7%; see Fig. 8 A for representative images). Increased co-localization of Cdc12(W367K)-GFP with Hsp104 is consistent with Hsp104 binding to Cdc12 molecules that are not engaged in CTD-mediated homo-oligomerization. Note that the foci formed by GFP-tagged Cdc12 in these experiments are an artifact of the GFP tag and should not be taken as evidence of Cdc12 aggregation per se; instead, they conveniently concentrated Cdc12-GFP signal in a way that allowed us to detect co-localization with Hsp104.

**Figure 8.**
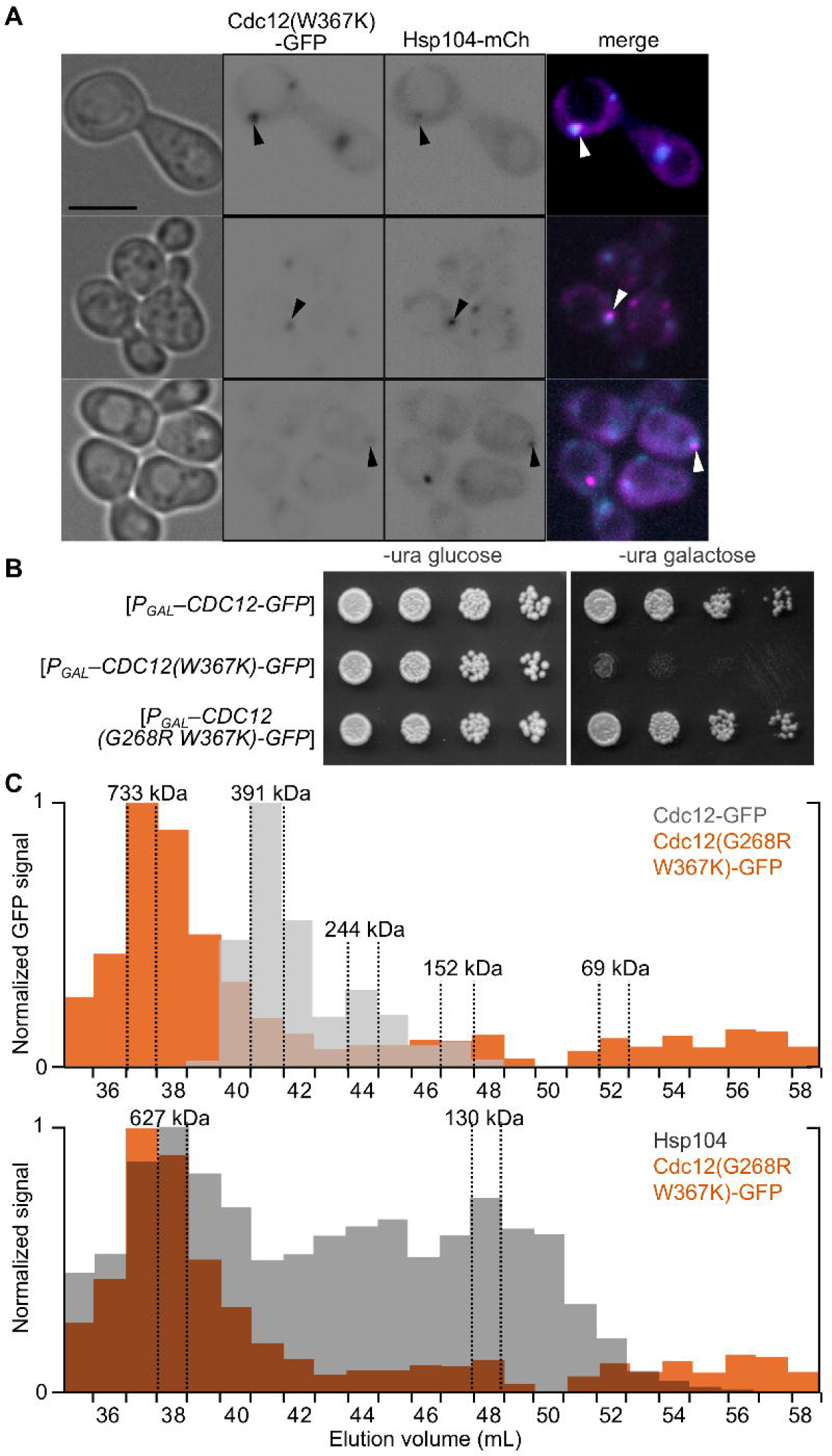
Hsp104 co-localizes and co-elutes with monomeric mutant Cdc12. (A) Representative images of cells of *HSP104-mCherry* strain H06799 carrying Cdc12(W367K)-GFP-expressing plasmid C6F1B99E cultured in selective medium with 2% galactose and imaged with transmitted light (first column of images), the GFP cube (second column, 60-msec exposure), or the Texas Red cube (third column, 250-msec exposure). The fourth column shows false-colored overlay/merge images. Arrowheads point to overlapping foci. Scale bar is 5 µm. (B) Serial dilutions of cells of strain BY4741 carrying plasmid 1EB21EA6 were spotted on selective medium with 2% glucose or 2% galactose and incubated at 30°C for 3 days. (C) As in Fig. 3 B, SEC analysis of lysates of cells of strain BY4741 carrying plasmid pMVB2 (“Cdc12-GFP”) or YEp-Cdc12(G268R W367K)-GFP (“Cdc12(G268R W367K)-GFP”). Top plot shows results from blotting with anti-GFP antibodies. The data for Cdc12-GFP (gray bars) are the same as in Fig. 3 B, in which only fractions 38-48 were analyzed. For the Cdc12(G268R W367K)-GFP lysate, additional fractions were analyzed. kDa values above peaks indicate the calculated molecular weight for the largest molecules in that fraction, based on molecular weight standards analyzed in a separate SEC run. The 69-kDa fraction is where monomeric Cdc12(W367K)-GFP would be expected to elute. The bottom plot has the same values for GFP detection in the Cdc12(G268R W367K)-GFP lysate (orange) and also includes values for detection of Hsp104 on the same blot using anti-Hsp104 antibodies (dark gray).

We found independent evidence of Hsp104 association with monomeric Cdc12 in SEC experiments with yeast lysates in which we exploited the exclusionary effect of the G268R mutation in the Cdc12 G interface (Johnson *et al*., 2015) to overcome the lethality of Cdc12(W367K) overexpression (Fig. 8 B). In contrast to the dominant-negative behavior of CTD mutants, mutant septins harboring substitutions in the G interface are rendered innocuous via prolonged G interface binding to cytosolic chaperones and exclusion from septin hetero-octamers (Johnson *et al*., 2015; Hassell *et al*., 2022). Rather than eluting as monomers, overexpressed Cdc12(G268R W367K)-GFP eluted in fractions with a peak corresponding to ∼700-kDa (Fig. 8 C). This peak matches the expected size of an Hsp104 hexamer bound to Cdc12(G268R W367K)-GFP (694 kDa) and, when we exposed the blot to an anti-Hsp104 antibody, overlapped with a “shoulder” on the peak of Hsp104 elution (Fig. 8 C). The ∼627-kDa fraction with the peak of Hsp104 matches the size of Hsp104 hexamers without bound substrate (624 kDa). Together with the co-localization of Hsp104-mCherry and excess Cdc12(W367K)-GFP, these findings strongly support a model in which hexameric Hsp104 binds and sequesters excess Cdc12 monomers.

### Hsp104 is required to maintain super-stoichiometric expression of septins that cannot undergo CTD-mediated homo-oligomerization

Despite lacking a CTD (Fig. 1 A), Cdc10 can homo-oligomerize: purified Cdc10-6xHis binds purified GST-Cdc10 in vitro (Versele *et al*., 2004), Cdc10 expressed individually in *E. coli* purifies as apparent homo-oligomers (Baur *et al*., 2019), and available in vivo evidence suggests Cdc10 homodimerizes prior to Cdc10– Cdc3 interaction during octamer assembly (Weems and McMurray, 2017). If a septin CTD is required for Hsp104 independence, then the absence of Hsp104 should prevent high-level accumulation of Cdc10. Alternatively, if homodimerization per se is sufficient to stabilize an excess septin against degradation despite the absence of Hsp104, then excess Cdc10 should not require Hsp104 for stability. In support of the first hypothesis, Cdc10-GFP overexpressed using the *MET25* promoter accumulated to high levels in wild-type cells but not in *hsp104*Δ cells, where only faint bud neck signals were detected (Fig. 9 A). Quantification of cytoplasmic GFP signal confirmed that Cdc10-GFP levels were significantly lower in *hsp104*Δ cells than in wild-type cells (Mann Whitney U test *P* = 3.7×10^−9^, Fig. 9 B). Furthermore, analogous to the effects of Cdc12ΔCTD overexpression, overexpression of Cdc10Δ 0, a version of Cdc10 lacking a critical component of the homodimerization interface that allows hetero-octamers to assemble (McMurray *et al*., 2011), was lethal to wild-type cells but *hsp104*Δ cells were completely unaffected (Fig. 9 A). Thus homodimerization per se is not sufficient to stabilize an excess septin in the absence of Hsp104.

**Figure 9.**
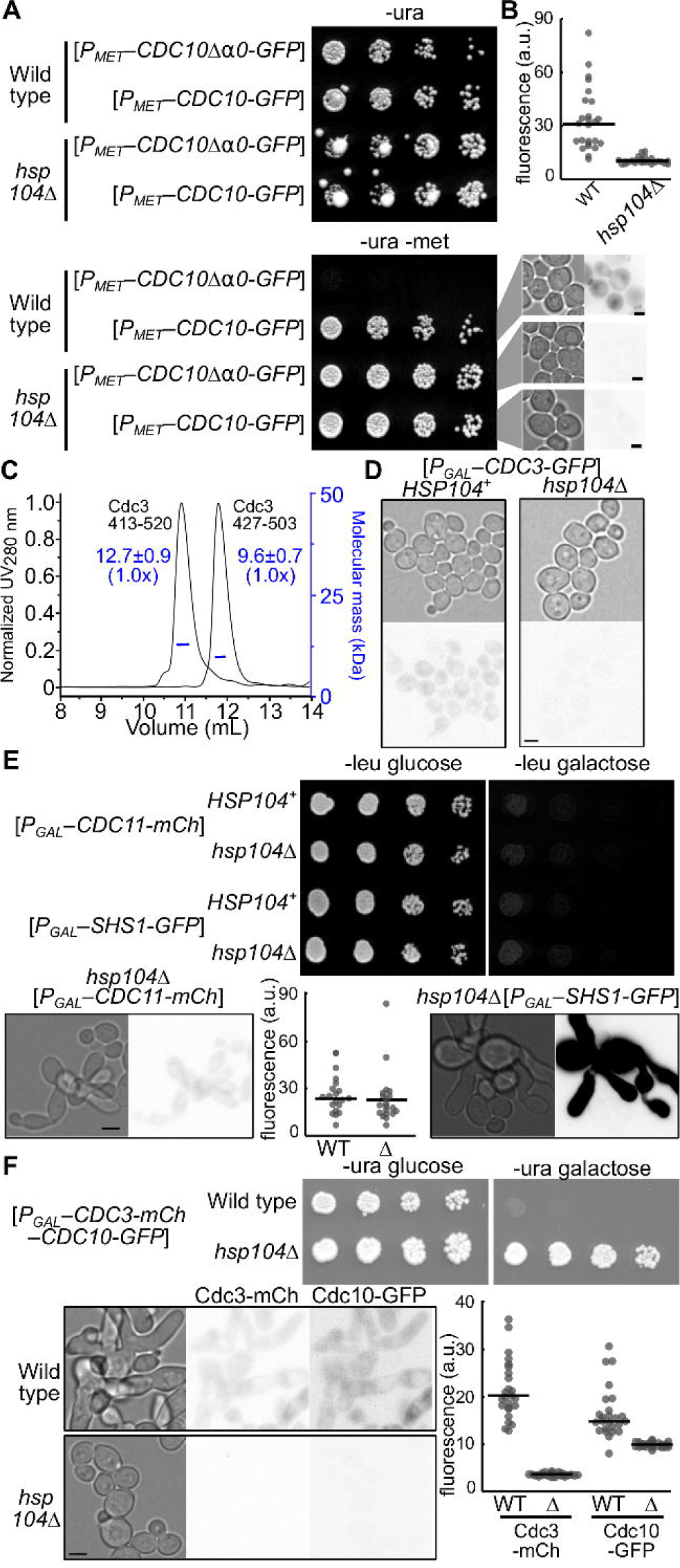
Specific septins require Hsp104 for super-stoichiometric stability. (A) Five-fold serial dilutions of yeast cells were spotted on medium selective for the indicated *URA3*-marked plasmid and either repressing (containing methionine, “-ura”) or inducing (lacking methionine, “-ura -met”) expression of the indicated tagged septin. Plates were incubated at 30°C and imaged after 2 days. Micrographs at right show cells scraped from that plate and imaged by transmitted light (left) or with the GFP cube (right, 60-msec exposure). Scale bar is 2 µm. Strains were BY4742 (“Wild type”) and JTY4013 (“*hsp104*Δ”) and plasmids were pMETp-CDC10-GFP (“*P_MET_–CDC10-GFP*”) and pJT3456 (“*P_MET_–CDC10*Δ *0-GFP*”). (B) Fluorescence in a 10-pixel-diameter circular area was quantified for 25 cells each from the colonies as shown in (A) of BY4742 (“WT”) or *hsp104*Δ yeast strain JTY4013 (“*hsp104*Δ”) carrying the plasmid pMETp-CDC10-GFP. “a.u.”, arbitrary units; lines indicate medians. (C) As in Fig. 2 B, SEC-MALS analysis of purified Cdc3 CTD. The CTD residues present in the two constructs are indicated. (D) The same strains as in (A) but carrying plasmid pMVB1 (“*P_GAL_–CDC3-GFP*”) were cultured in medium selective for the plasmid and containing 2% galactose to induce Cdc3-GFP expression. Shown are representative images of cells visualized by transmitted light (left) or with the GFP cube (right, 60-msec exposure). Scale bar is 2 µm. (E) Top, serial dilution as in (A) but with glucose as the repressor and galactose as the inducer, and *LEU2*-marked plasmids pGF-IVL-287 (“*P_GAL_–CDC11-mCh*”) and pGF-IVL-286 (“*P_GAL_–SHS1-GFP*”). Below, cells from the galactose plate, imaging Cdc11-mCherry (250-msec exposure) or Shs1-GFP (250-msec exposure). Scale bar is 5 µm. The plot quantifies Cdc11-mCherry fluorescence in as in (B) but with 20 cells per genotype; “Δ” indicates *hsp104*Δ. (F) As in (E) but with *URA3-*marked plasmid G01273 and imaging Cdc3-mCherry (250-msec exposure) and Cdc10-GFP (250-msec exposure). Scale bar is 5 µm.

Conversely, while Cdc3 has a CTD (Fig. 1 A), the purified Cdc3 CTD does not homo-oligomerize at all (Fig. 9 C), and only a fraction of purified full-length Cdc3 homo-oligomerizes (Hassell *et al*., 2022). By contrast, purified full-length Cdc11 is nearly 100% homodimers and this homodimerization requires the CTD (Brausemann *et al*., 2016). If CTD-mediated homo-oligomerization, not just the presence of a CTD or homodimerization alone, is required to stabilize an excess septin in the absence of Hsp104, then excess Cdc3 molecules, but not excess Cdc11 molecules, should require Hsp104 for their stability. Indeed, compared to the fluorescence observed in wild-type cells overexpressing Cdc3-GFP from the *GAL1/10* promoter, signal in *hsp104*Δ cells was greatly reduced (Fig. 9 D). By contrast, Hsp104 was not required for the accumulation of high levels of Cdc11 (or its paralog Shs1, which also has a CTD) or for the septin dysfunction known to result from overexpression of either of these septins (Sopko *et al*., 2006) (Fig. 9 E). Quantification confirmed that Cdc11-mCherry levels were not significantly different in *hsp104*Δ cells than in wild-type cells (Mann Whitney U test *P* = 0.19, Fig. 9 E).Thus having a CTD with coiled-coil-forming potential is not enough to stabilize a septin: that CTD must also confer efficient homo-oligomerization.

The low levels of Cdc10-GFP and Cdc12(W367K)-GFP found at the bud neck upon attempted overexpression of those septins in *hsp104*Δ cells (Fig. 4 C, Fig. 9 A) suggested that in the absence of Hsp104 the mutant septins are protected from degradation if they successfully incorporated into septin-hetero-oligomers or filaments. If protection from degradation reflects burial of otherwise exposed proteasome-targeting regions in septin-septin interfaces (e.g. the G interface), then we wondered if co-overexpressing two septins that interact via the G interface would bury these regions and provide protection. We recently showed that simultaneous co-overexpression of tagged versions of Cdc3 and Cdc10 –– which interact via a G interface (Fig. 1 A) –– severely perturbs septin function, as evidenced by highly elongated cells (Benson and McMurray, 2023). Here we found that this functional perturbation also manifests as defects in colony formation at 30°C (Fig. 9 F). This effect was entirely lost in *hsp104*Δ cells and levels of both tagged Cdc3 and Cdc10 were greatly reduced (Fig. 9 F, Mann Whitney U test *P* = 1.4×10^−9^ for Cdc3-mCherry, 2.6×10^−8^ for Cdc10-GFP), suggesting that the G interface partners could not protect each other.

When Cdc3 is specifically tagged with a fragment of the YFP derivative Venus (“V_C_”) and co-overexpressed with Cdc10 tagged with a BiFC-compatible Venus fragment (“V_N_”), in addition to inducing cell elongation, the “excess” molecules of Cdc3-V_C_ and Cdc10-V_N_ co-assemble into stable, plasma-membrane-associated filaments composed only of tagged Cdc3 and Cdc10 (Benson and McMurray, 2023). Even in this scenario, the absence of Hsp104 prevented both the high-level accumulation of BiFC fluorescence and the associated cell elongation and plasma membrane localization (Fig. S12 B). Instead, only low levels of Cdc3–Cdc10 BiFC signal were detected at bud necks (Fig. S12 B), similar to the situation with attempted overexpression of Cdc10-GFP alone. Unique among the five mitotic yeast septins, Cdc3 has a ∼100-residue N-terminal domain (Fig. 1 A) that we previously proposed may occlude the Cdc3 G interface and prevent Cdc3–Cdc10 interaction until Cdc3 interacts with Cdc12 (Weems and McMurray, 2017). We tried removing the Cdc3 NTD to improve Cdc3–Cdc10 G dimerization, but this mutation –– which does not block the assembly of membrane-associated filaments upon Cdc3-V_C_–Cdc10-V_N_ co-overexpression in wild-type cells (Benson and McMurray, 2023) –– failed to restore the dominant perturbation of septin function or high-level accumulation of Cdc3–Cdc10 BiFC signal in *hsp104*Δ cells (Fig. S12 B). Thus even the ability to polymerize into membrane-associated filaments does not protect “excess” Cdc10 from degradation when Hsp104 is absent.

Immunoblotting confirmed these findings: whereas steady-state overexpression levels of tagged Cdc11 (∼15-fold higher than endogenous/native levels) or Shs1 were unaffected by *HSP104* deletion, Cdc3 and Cdc10 levels were reduced in *hsp104*Δ cells to an amount equivalent to endogenous/native levels (Fig. S13). Thus all available evidence supports the idea that CTD-mediated homo-oligomerization lifts the Hsp104 requirement for a septin to accumulate at superstoichiometric levels, and in the absence of Hsp104 only incorporation into native-like hetero-octamers is able to protect a CTD-less septin from degradation.

### Septin-encoding mRNAs differ in copy number per cell and subcellular localization

Our findings reveal unappreciated roles for transient septin homo-oligomerization and the cellular proteostasis machinery in the ability of cells to maintain excess septin subunits. We discovered these roles via experimental, high-level overexpression of single septin subunits. What is the natural context in which these maintenance mechanisms may have evolved? We reasoned that if the endogenous septin genes are infrequently transcribed, this could create a scenario in which the number of septin-encoding mRNAs per cell fluctuates widely within a population of cells. Efficient translation of septin mRNAs (estimates range from five to 11 ribosomes per septin mRNA (Arava *et al*., 2003)) would then generate transient imbalances in subunit stoichiometry within a given cell. Indeed, existing evidence from transcriptome-wide studies pointed to a low number of septin-encoding mRNAs per cell. For Cdc12, estimates ranged from 0.3 to 3.2 copies/cell, depending on the technique (Velculescu *et al*., 1997; Holstege *et al*., 1998; Miura *et al*., 2008); other septin mRNAs were in a similar range (Cdc3 1.5-2.9 copies/cell; Cdc10 1.7-3.0; Cdc11 1-3.1 (Velculescu *et al*., 1997; Holstege *et al*., 1998; Miura *et al*., 2008)).

These published values come from relatively indirect assays. To directly determine cell-to-cell variation in septin mRNA copy number, we used single-molecule inexpensive fluorescence in situ hybridization (smiFISH) (Tsanov *et al*., 2016). We first validated our experimental approach by localizing in the same cells mRNAs encoding Ash1, which are specifically localized to the bud tip (Long *et al*., 1997), and Abp140, which are specifically localized in the mother cell distal from the bud (Kilchert and Spang, 2011) (Fig. S14). Next, using mutant cells missing or depleted for the target mRNA to control for non-specific background signal, we localized mRNAs encoding Cdc3 and/or Cdc10 and found that the number of septin mRNAs per cell was low and highly variable (Fig. 10 A, Fig. S14), in agreement with the indirect estimates. Importantly, Cdc3 and Cdc10 mRNA copy numbers per cell varied independently (Fig. 10 A), providing many potential opportunities for stoichiometric imbalances.

**Figure 10.**
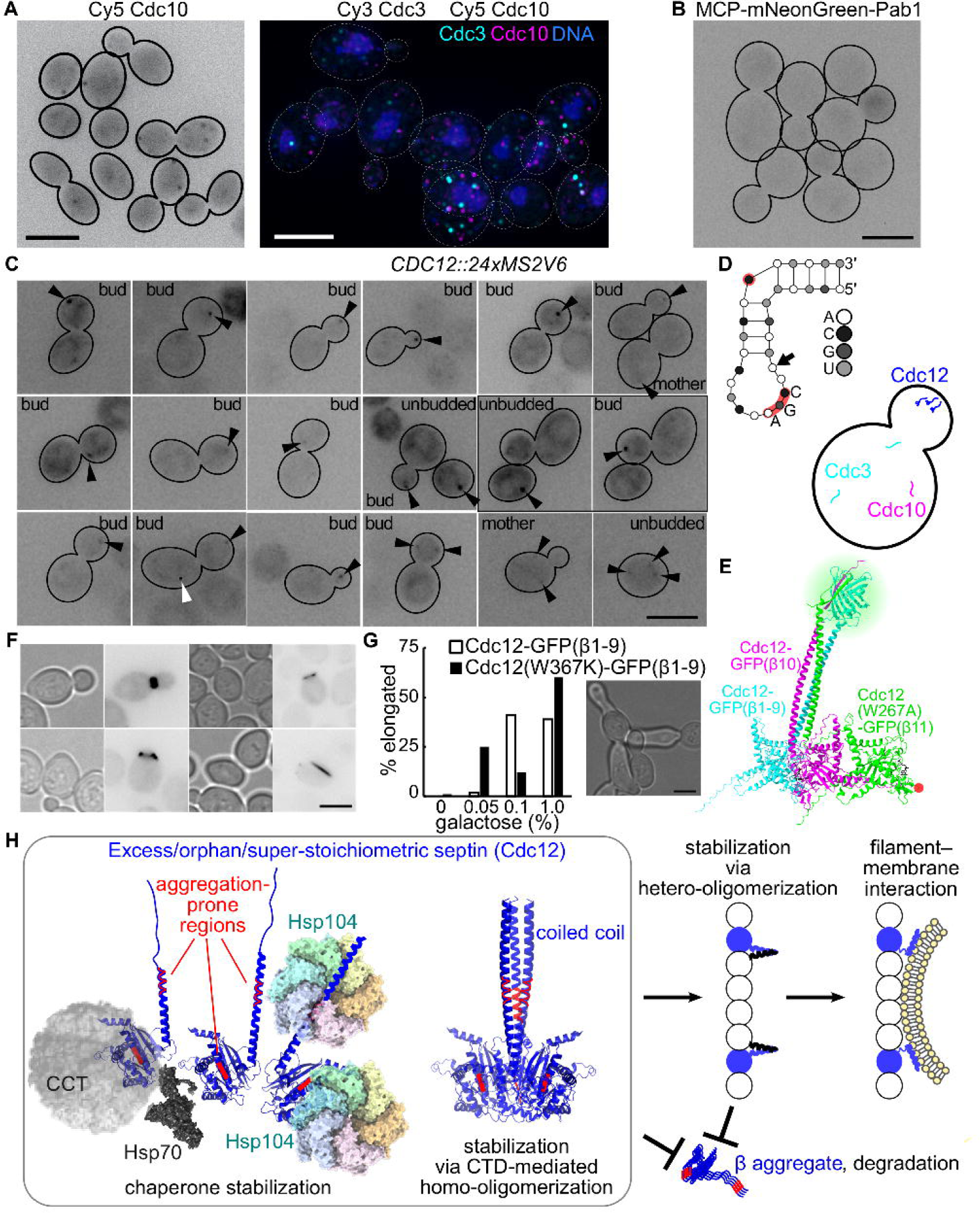
Low, variable numbers of septin-encoding mRNAs with distinct subcellular localization patterns create opportunities for transient septin homo-oligomerization. (A) Septin mRNAs detected by smiFISH in wild-type (strain BY4743) cells. Left, cells exposed only to a Cy5-labeled Cdc10 probe set. Right, cells exposed to a Cy5-labeled Cdc10 probe set and a Cy3-labeled Cdc3 probe set plus DAPI staining of the DNA. Solid (left) or dashed (right) lines outline the cell walls. Scale bar, 5 µm. (B) Haploid cells of strain 7M3Q9V2W, which lacks MS2 stem loops but expresses mNeonGreen-tagged MS2 coat protein fused to a portion of the Pab1 protein, were grown in liquid culture containing 1 mM methionine to mid-log phase and imaged for green fluorescence. (C) As in (B) but with strain 5B8N4K9X in which 24 copies of the MS2V6 stem loop are inserted in the 3’UTR of Cdc12. Labels in each panel indicate whether the cell(s) with foci are buds, mother cells, or unbudded. Black arrowheads indicate foci of tagged Cdc12 mRNA. White arrowhead is a “hot pixel” in the imaging camera and does not reflect mNeonGreen signal. Boxes surround pairs of images of the same cells but at different focal planes. Scale bar, 5 µm. (D) Schematic illustration, in the style of a recent study identifying RNA motifs involved in bud tip localization (Seiler *et al*., 2024), of key features of the predicted stem-loop formed by a 29-nt region of the Cdc12 mRNA (-39 to -11 relative to the start codon). The CGA trinucleotide motif in the loop is shown and red highlights cytosine-containing motifs key for She2–She3 binding to other mRNAs; the arrow points to an adenosine at the 5’ end of the loop. Cartoon below illustrates an example of distinct copy number and localization patterns of different septin mRNAs. (E) Schematic of trimolecular fluorescence complementation (TriFC) assay to detect Cdc12 homotrimerization in vivo. Ribbon diagram shows top-ranked AlphaFold3 prediction for the complex formed by three C-terminal fusions of GFP fragments to Cdc12. Note that the Cdc12-GFP(β11) fusion harbors the W267A mutation (highlighted in red) to inhibit interactions between the Cdc12 GTPase domains. (F) Representative micrographs of TriFC signal detected in diploid yeast cells (strain BY4743) grown in medium with 2% raffinose (0% galactose) and selective for the plasmids encoding the three fusions, which were D2AC2EB4 (“Cdc12-GFP(β1-9)”), 76068B19 (“Cdc12-GFP(β10)”), and E2438448 (“Cdc12(W267A)-GFP(β11)”). Scale bar, 5 µm. (G) The percentage among 100 cells analyzed per sample that were elongated (length to width ratio >2, a symptom of septin dysfunction) within cultures with the indicated galactose concentration and expressing Cdc12-GFP(β10), Cdc12(W267A)-GFP(β11), and either wild-type Cdc12-GFP(β1-9) or, as indicated, a derivative carrying Cdc12 mutation W367K (from plasmid 15C00ABB). Right, example elongated cells. Scale bar, 5 µm. (H) Schematic illustration, adapted from Hassell et al., 2022, of the proposed role for CTD-mediated coiled coil homo-oligomerization in the pathway by which excess/orphan molecules of a budding yeast septin (using Cdc12 as an example) assemble post-translationally into hetero-octamers. Two parallel modes of conformational stabilization are shown, one mediated by chaperones and the other by coiled-coil homo-oligomerization. Hsp104 binds aggregation-prone sequences (red). Both modes prevent aggregation and proteasome-mediated degradation. Co-assembly into a septin hetero-octamer also stabilizes. In the case of Cdc12, hetero-oligomerization requires dissociation of homo-oligomeric coiled coils and assembly of hetero-dimeric coiled coils, which also dissociate when septin filaments interact with the plasma membrane.

We monitored Cdc12 mRNA localization using an in vivo method in which an array of binding sites for the MS2 coat protein (MCP) is inserted in the 3’ UTR of the mRNA of interest and co-expression of fluorescently tagged MCP generates cytoplasmic foci marking the tagged mRNA (Haim *et al*., 2007; Haim-Vilmovsky and Gerst, 2009). A recent study (Li *et al*., 2022) corrected MS2-induced destabilization of mRNA by fusing full-length poly(A)-binding protein, Pab1, to MCP, where Pab1 promotes termination of translation of the tagged mRNA and prevents nonsense-mediated decay (Amrani *et al*., 2004). To prevent binding of the MCP-Pab1 fusion to other RNAs, that study mutated Phe residues in two of the four Pab1 RNA Recognition Motifs (RRMs) (Li *et al*., 2022). Pab1 lacking RRM4 altogether retains the ability to stabilize a tethered mRNA (Deardorff and Sachs, 1997), so we used a shorter Pab1 fusion with RRM4 deleted and the F366V mutation in RRM2. Twenty four copies of an improved MS2 binding site (MS2V6) (Tutucci *et al*., 2018) were inserted immediately 3’ of the stop codon in the genomic *CDC12* locus. Expression of mNeonGreen-tagged MCP-Pab1 fusion was under control of the *MET25* promoter, allowing repression via methionine in the medium, which was crucial to maintain low levels that did not form foci in the absence of MS2 binding sites (no focus among 444 cells examined; representative images shown in Fig. 10 B).

Unexpectedly, discrete MCP-mNeonGreen-Pab1 signals representing Cdc12 mRNA were most frequently found as single bright foci in the buds of budded cells (14 of 20 cells with foci; Fig. 10 C). Signal was much less frequently detected in unbudded (3 of 20) or budded mother (2 of 20) cells (Fig. 10 C). Cells with more than one focus were also rare (3 of 20) and these foci were comparatively dim (Fig. 10 C). We saw no more than three Cdc12 mRNA foci per cell, in agreement with the published estimates of Cdc12 mRNAs copy number. Since we intentionally kept MCP levels low to prevent false-positive signals, we cannot interpret cells with no detectable signal (which were the majority) as having no Cdc12 mRNA; they may just not express enough MCP. Most (12 of 14) of the bright foci in buds were in the half of the bud distal from the mother along the mother-bud axis (Fig. 10 C). We conclude that Cdc12 is among the ∼35 other known mRNAs –– most encoding proteins involved in cell polarity –– that cluster at the bud tip (Shepard *et al*., 2003; Andoh *et al*., 2006; Aronov *et al*., 2007; Oeffinger *et al*., 2007; Takeuchi-Andoh *et al*., 2017). While additional studies are needed to determine what directs Cdc12 mRNA localization, a 29-nt sequence near the 5’ end of the Cdc12 mRNA possesses all the key features of the She2–She3 binding sites found in bud-tip-localized mRNAs, including a predicted stem-loop structure with an adenosine at the 5’ end of the loop, the loop sequence ACGAA, and, seven nucleotides away from the cytosine in that sequence, a single cytosine in a bulge of the stem (Jambhekar *et al*., 2005; Olivier *et al*., 2005; Müller *et al*., 2011; Seiler *et al*., 2024) (Fig. 10 D). Thus Cdc12 mRNA localizes differently than Cdc3 and Cdc10 (Fig. 10 D), a pattern that likely creates further opportunities for transient subunit imbalances and transient homo-oligomerization (see Discussion).

### Detecting Cdc12 homotrimers in vivo using split-GFP reveals high levels in rare cells

Our model predicts that orphan septin homo-oligomers form in only some cells and, when they do form, they are transient, making them challenging to detect in living cells without high-level overexpression, which might artificially drive interactions. However, a few mRNAs must be sufficient to generate enough protein to replenish the septin supply that is reduced by about half with each cell division (McMurray and Thorner, 2008), meaning ∼5,500 Cdc12 polypeptides (Ho *et al*., 2018), a value consistent with previous estimates of protein yield per mRNA (Futcher *et al*., 1999; Lahtvee *et al*., 2017). If so, then in the rare cells that –– due to stochastic variation of septin mRNA copy number –– have too many Cdc12 mRNAs, tens of thousands of excess Cdc12 proteins should be made. If a significant fraction form homotrimers, we reasoned that we might be able to detect them in those rare cells, provided we use an assay that “traps” otherwise transient protein-protein interactions.

We previously used a bipartite split-GFP approach (bimolecular fluorescence complementation, BiFC) to detect transient septin-chaperone binding in vivo (Denney *et al*., 2021), wherein the irreversible reconstitution of the split GFP creates a long-lived fluorescent signal from a short-lived binding event. Since septin BiFC could reflect cross-filament interactions (Fig. 1 A), we used a tripartite (TriFC) approach with Cdc12 to specifically detect homotrimers. In the TriFC system, the 11-stranded GFP β barrel is split into three fragments corresponding to β strands 1-9, 10, and 11 (Cabantous *et al*., 2013; Finnigan *et al*., 2016) (Fig. 10 E). We looked for fluorescence in cells co-expressing three Cdc12 fusion proteins with the GFP fragments plus untagged endogenous Cdc12. To favor the coiled-coil interaction as a driver of TriFC, we included in the Cdc12-GFP(β11) fusion the G-interface mutation W267A, known to weaken non-native Cdc12–Cdc12 G interface interactions (McMurray *et al*., 2011). Indeed, in all five AlphaFold3 predictions for the complex formed by the three fusions, the GFP was reconstituted at the tip of the same three-helix coiled coil supported by our other experiments, and while all models included at least one interaction between GTPase domains, the Cdc12(W267A)-GFP(β11) G interface was never engaged (Fig. 10 E, Fig. S15 A).

To have some control over the balance of expression of the three fusions, we expressed Cdc12-GFP(β1-9) from the *GAL1/10* promoter, while the other two fusions were expressed from the *CDC12* promoter. In 2% raffinose (no galactose), *GAL1/10* transcription is neither repressed nor induced (Johnston *et al*., 1994), which we reasoned would best match the low level of *CDC12* promoter-driven transcription. However, given four distinct versions of Cdc12 expressed in these cells (untagged, β1-9, β10, and β11), even if expression levels were perfectly equal, at most 9.375% of Cdc12 trimers would be comprised of the three distinct TriFC fusions and thus capable of generating fluorescence. All three fusion proteins were expressed from distinct plasmids, which likely vary in copy number per cell, and expression of each individual gene is also likely to be “bursty”, further decreasing the expected frequency of TriFC-positive cells. Nonetheless, we were able to detect a small population of cells with bright TriFC signals (see Fig. 10 F for representative images). As expected, we only saw such cells in 2% raffinose, and introducing the W367K mutation that blocks homo-oligomerization in vitro into the Cdc12-GFP(β1-9) fusion completely eliminated TriFC signal (no TriFC-positive cell among 17,379 analyzed).

The lack of fluorescence in cultures with galactose was expected, since an excess of Cdc12-GFP(β1-9) should form non-fluorescent homotrimers. In these cultures cell morphology was abnormal and worsened with increasing galactose concentrations (Fig. 10 G), indicative of dominant perturbation of septin function by Cdc12-GFP(β1-9) as it displaced Cdc12 molecules with no or smaller tags in hetero-octamers and filaments. The same or worse galactose-dependent morphological defects were observed with Cdc12(W367K)-GFP(β1-9) (Fig. 10 G), providing a convenient confirmation that the mutant protein was expressed at equivalent levels to wild type Cdc12-GFP(β1-9).

Whereas our model predicts that Cdc12 homotrimers spend the entirety of their fleeting existence in the cytoplasm, TriFC signal was found either at cortical current/former sites of budding (where rings of septin filaments are normally found (Haarer and Pringle, 1987; Ong *et al*., 2014)), or in long “fibers” (Fig. 10 F) that have also been previously reported for C-terminal Cdc12 fusions (McMurray, 2016). The irreversibility of split-GFP reconstitution explains the cortical/bud neck signals: coiled coil metastability should allow one of the Cdc12 fusions to disengage from the homotrimeric coiled coil and engage with the CTD of Cdc3 in the context of a septin hetero-octamer, without dissociating the reconstituted GFP. Indeed, this very scenario was represented in the highest-ranked AlphaFold3 prediction when we included Cdc3, Cdc10, and Cdc11 with the three Cdc12 fusions (Fig. S15 B). Incorporation of one Cdc12 fusion into septin filaments recruits to the bud neck the other two Cdc12 fusions still linked via the TriFC event, analogous to the bud neck signals we previously observed by BiFC for septin interactions with cytosolic chaperones (Denney *et al*., 2021). These data are thus fully consistent with a model in which CTD-mediated homodimers and homotrimers are transient species formed by superstoichiometric molecules of Cdc12 en route toward hetero-octamer and filament assembly once partner septins become available.

## DISCUSSION

Various factors dictate the fate of excess subunits of heteromeric protein complexes. A previous study used yeast strains with an extra copy of a single chromosome to generate a two-fold excess of individual subunits and found that 73% of proteins that function in complexes either aggregated or were degraded (Brennan *et al*., 2019). Those authors proposed that “excess proteins that do not aggregate or get degraded are simply less aggregation-prone and lack the signals that target them for degradation” (Brennan *et al*., 2019). While in this study Cdc3 and Cdc12 were among the 437 proteins found in “aggregates” (as defined by pelleting during centrifugation of lysates) (Brennan *et al*., 2019), we have clearly shown here and elsewhere (Hassell *et al*., 2022) that individual yeast septins expressed at high levels are not irreversibly aggregated. Thus despite being intrinsically aggregation-prone and possessing signals able to target them for degradation, excess yeast septins avoid these fates via a combination of assistance from chaperones and coiled-coil-mediated homo-oligomerization (Fig. 10 E).

Homo-oligomerization by septins that do not normally occupy two adjacent positions within a septin filament has previously been viewed as a vestige from the evolutionary past, potentially generating “off-pathway” homo-oligomers (Hassell *et al*., 2022; Brown and McMurray, 2025) or, in some cases, alternative hetero-oligomers with distinct functions (hetero-hexamers vs hetero-octamers) (Johnson *et al*., 2020). Roles in hetero-oligomer assembly for the conserved predicted coiled-coil-forming sequences in many septin CTDs have remained mostly enigmatic, particularly with regard to their metastable character. In the septin literature, CTDs mostly contribute to higher-order septin hetero-oligomer structure and function: how filaments interact with each other and with membranes and/or other proteins (Finnigan *et al*., 2015a; Sala *et al*., 2016; Cannon *et al*., 2019; Leonardo *et al*., 2021; Woods *et al*., 2021; Castro *et al*., 2023). There is no place for septin homotrimers in current models, which is why we were so intrigued by our observation of Cdc12 CTD homotrimerization in vitro and particularly the presence of a highly conserved homotrimerization motif. Indeed, to our knowledge the Cdc12 CTD is the first septin coiled coil to exhibit trimerization. Human SEPT1, SEPT2, SEPT4, SEPT5, SEPT6, SEPT7, SEPT8, SEPT14 and SmSEPT10 CTDs are all homodimers (Leonardo *et al*., 2021; Cavini *et al*., 2023, 2024).

While we found evidence of Cdc12 homotrimers in vivo, Cdc12 homodimers were also present and appear to function in the same way: together with Hsp104, they help protect excess Cdc12 molecules against degradation (Fig. 10 E). The Cdc12(W367K E368V) mutant was particularly informative: if E368 is required for Hsp104–Cdc12 binding, then Hsp104 should be incapable of stabilizing excess Cdc12(W367K E368V) molecules, yet Cdc12(W367K E368V)-GFP accumulated to high levels and dominantly interfered with septin function in the absence of Hsp104 (Fig. 4 B). Thus the key property deciding the distinct fates of excess Cdc12(W367K) versus Cdc12(W367K E368V) is most likely the ability of Cdc12(W367K E368V) to homo-oligomerize to some degree. We conclude that an important, previously unappreciated function of CTD-mediated homo-oligomerization is to promote the assembly of a transient, metastable reservoir of super-stoichiometric septins that are competent to incorporate into septin hetero-octamers and filaments once partner septins become available (Fig. 10 H). Homotrimers would simply have a greater “storage capacity” than homodimers. Indeed, studies of the RhxxhE motif in other coiled coils found that while it is enriched in homotrimers, many proteins with the motif form homodimers (Testa *et al*., 2009). In Cdc12 and homologous septins, the motif contains bulky side chains (Trp and Phe) that are not typically found in the core of dimeric coiled coils. These residues likely favor homotrimerization to accommodate these side chains in a slightly larger radius of a trimeric coiled coil (Woolfson, 2023). Cdc12 homotrimerization also fits with our unexpected finding that Cdc12 mRNAs tend to cluster near the bud tip, whereas Cdc3 and Cdc10 mRNAs show no clear spatial bias. Our co-translational assembly model (Hassell *et al*., 2022) predicts that full-length Cdc3 and Cdc10 polypeptides diffuse from their sites of synthesis and bind nascent Cdc12 polypeptides emerging slowly from polysomes. Confining translation of multiple Cdc12 mRNAs to a small region of the cytoplasm likely creates opportunities for “local” Cdc12 homo-trimerization while other septin proteins are in transit from remote locations (Fig. 10 D).

We could find no precedent in the literature for the loss of high-level accumulation we observed for specific septins in *hsp104*Δ cells. Deletion of *HSP104* did not prevent the *GAL1/10* promoter-driven accumulation of any of many aggregation-prone human proteins or fragments thereof, including Huntingtin (Giorgini *et al*., 2005), TDP-43, FUS, -synuclein, or EWSR1 (Jackrel and Shorter, 2014; Jackrel *et al*., 2014). On the contrary, Hsp104 is *required for* the degradation of yeast-expressed aggregation-prone versions of human ataxin-1 (Lee and Goldberg, 2010), apolipoprotein B, and cystic fibrosis transmembrane conductance regulator (CFTR) (Doonan *et al*., 2019). We also find it remarkable that of >1,000 essential proteins in *S. cerevisiae*, Hsp104(G214S T499I) specifically perturbs the septin Cdc12 (Schirmer *et al*., 2004); the additional Hsp104 binding sites in the Cdc12 CTD and their persistent exposure for Hsp104 binding presumably explain why other septins are not targets of Hsp104(G214S T499I).

How are excess septins targeted for proteolysis? Like Parkin, yeast Dma1 and Dma2 are E3 ubiquitin-protein ligases that mediate ubiquitinylation of septins, and the mammalian Dma homolog, RNF8, promotes ubiquitinylation of human Septin-7 (Chahwan *et al*., 2013). Interestingly, Cdc12 K351 was found to be ubiquitinylated in a proteomic study (Swaney *et al*., 2013), but this cannot be the only lysine mediating Cdc12 degradation, since excess Cdc12ΔCTD was degraded in *hsp104*Δ cells (Fig. 4 C). Future work will be required to dissect the molecular properties that make excess septins targets for Hsp104 intervention and, when Hsp104 is absent, intervention by the proteasome. Such studies could reveal valuable insights regarding the molecular etiology of human diseases linked to septin aggregation and degradation.

Earlier in vitro studies supported a model in which a G-partnerless septin is prone to loss of bound nucleotide, which destabilizes the G interface and predisposes to β aggregation (Garcia *et al*., 2007; Pissuti Damalio *et al*., 2012; Kumagai *et al*., 2019). It is easy to imagine how this could work, given the proximity of the nucleotide-binding pocket and key features of the G dimer interface to the aggregation-prone β4 strand. Consistent with this hypothesis, a Cdc10 mutant that likely cannot bind GDP shows prolonged interaction with Hsp104 (Johnson *et al*., 2015; Denney *et al*., 2021). However, in the non-native septin filaments composed only of “excess” Cdc3 and Cdc10 that assemble upon simultaneous co-overexpression of specific tagged forms of these two septins (Benson and McMurray, 2023), the nucleotide-binding pockets of Cdc3 and Cdc10 should be fully occupied, yet deletion of *HSP104* blocked accumulation of Cdc3–Cdc10 (Fig. 9 D). This finding thus points to the earliest stages of individual septin folding –– prior to septin-septin hetero-oligomerization –– as the crucial period for stabilization by Hsp104. In support of this model, we previously detected Hsp104 co-eluting specifically with newly-made molecules of a G-interface-mutant Cdc10 (Johnson *et al*., 2015).

If Hsp104 acts on individual septins before they hetero-oligomerize, why are superstoichiometric septins specifically susceptible, when hetero-oligomerization is the step where stoichiometry is defined? One possibility is that GTP/GDP availability is limited (Bianchi-Smiraglia *et al*., 2021; Scott *et al*., 2024), and excess septins are poor nucleotide scavengers and tend to be nucleotide-free and consequently misfolded. Alternatively, kinetics may be the key. Septins that are delayed in G dimerization are excluded from hetero-octamers by faster-associating competitor septins and their G interfaces are instead bound by chaperones (Johnson *et al*., 2015; Schaefer *et al*., 2016; Denney *et al*., 2021). If co-overexpressed Cdc3 and Cdc10 are slower to assemble into homodimers or non-native Cdc3–Cdc10 polymers than to incorporate into hetero-octamers, then the window of misfolding/aggregation opportunity is wider for these excess septins than for septins that quickly homo-oligomerize, and the requirement for Hsp104 is correspondingly greater. An untested prediction of this model is that, like its paralog Cdc11, purified Shs1 quickly forms CTD-mediated homodimers that would serve in cells to stabilize superstoichiometric Shs1 against aggregation/degradation. The existence of such homodimers in vivo likely explains why Cdc11 and Shs1 perturb septin hetero-octamer function when overexpressed: they “cap” octamer ends and interfere with polymerization into filaments.

Whereas previous studies emphasized the importance of a G-dimer partner in stabilizing the septin GTPase domain against aggregation and/or chaperone sequestration, here we propose that homo-oligomerization via the CTD –– considered an extension of the NC interface –– also performs this function (Fig. 10 H). A portion of the Cdc12 CTD has homology to a C-terminal domain of Ran GTPases that allosterically communicates with parts of the GTPase domain that change conformation upon GTP hydrolysis (Vetter *et al*., 1999; Nilsson *et al*., 2002; Weems *et al*., 2014). Removing the Cdc12 CTD accelerates GTPase activity (Versele and Thorner, 2004), providing more evidence of such communication. Several other mechanisms have also been proposed that link conformational changes at the G interface with changes in parts of the NC interface (Sirajuddin *et al*., 2009; Zeraik *et al*., 2014; Mendonça *et al*., 2024). It is also possible that CTD-mediated homo-oligomerization simply brings the GTPase domains in close proximity and promotes otherwise low-affinity associations between them (see Figs. S6, S15), which somehow protects against degradation (Fig. 10 H). Additional studies will be needed to better understand what stabilizes septins.

We are careful not to assume that aggregation necessarily precedes or drives septin degradation. What constitutes a septin “aggregate” is also imprecisely defined. Upon mild heat shock, some misfolded mutant yeast proteins transiently coalesce with Hsp104 and other chaperones, which protects them against proteasomal degradation (Boronat *et al*., 2023). Due to limited resolution, the mostly diffuse cellular fluorescence we observe for overexpressed, tagged single septins cannot be interpreted as “soluble” protein, and our SEC analysis would have missed large complexes that co-eluted with hetero-octamers or pelleted when we clarified our lysates. While the simplest explanation is that Hsp104 binding counteracts aggregation by excess septins, further studies will be required to rigorously test this model.

As it remains mechanistically unclear how PP2A–Rts1 (PP2A^Rts1^) phosphatase activity affects septins, we do not fully understand why *RTS1* deletion ameliorated the septin dysfunction observed upon overexpression of Cdc12(W367K). PP2A^Rts1^ dephosphorylates Shs1 (Dobbelaere *et al*., 2003), the other mitotic yeast septin with a C-terminal curvature-sensing amphipathic helix; deletion of both amphipathic helices severely cripples septin function (Woods *et al*., 2021). PP2A^Rts1^ acts on Shs1 during or just prior to cytokinesis (Dobbelaere *et al*., 2003) and may modify curvature sensing by the Shs1 amphipathic helix to accommodate changes in plasma membrane curvature. Additionally, a phosphoproteomic study identified Ser509 in the CTD of Cdc3 as undergoing Rts1-dependent dephosphorylation (Touati *et al*., 2019). Cdc3 Ser509 lies just C-terminal of the residues predicted to engage the CTD of Cdc12 in a heterodimeric coiled coil (Fig. 1 A) and adjacent to the Cdc3 “polybasic region 3” (**S**PVPT<u>KKK</u>GFL<u>R</u>, basic residues underlined), which may also influence membrane interactions (Cavini *et al*., 2024). The dephosphorylation events appear to drive “mobility” of septins within the septin ring (Dobbelaere *et al*., 2003) at the same time when polarized fluorescence microscopy reveals large-scale reorientation of the Cdc12 CTD (Vrabioiu and Mitchison, 2006; DeMay *et al*., 2011; McQuilken *et al*., 2017). These changes in higher-order septin structure likely reflect orchestrated rearrangement of the meshwork of septin filaments with respect to each other and the plasma membrane. CTD mutations in Cdc12 disrupt these dynamics; persistent phosphorylation of Shs1 and the Cdc3 CTD in *rts1*Δ cells may allow cytokinesis by restoring filament-filament interactions and/or interactions between filaments and the membrane. We speculate that the temperature-sensitive nature of the dysfunction caused by Cdc12 CTD mutations (and by *RTS1* deletion in cells with wild-type Cdc12 (Dobbelaere *et al*., 2003)) reflect temperature effects on plasma membrane properties that indirectly impinge on higher-order septin structure and function. Such effects likely explain why mutating R363 (also within the Cdc12 trimerization motif) to Lys has little effect on CTD oligomerization in vitro (Table S4) but makes cells sensitive to cold temperatures (Cvrcková and Nasmyth, 1993; Weems *et al*., 2014), which strongly alter membrane fluidity and lipid composition (Gunde-Cimerman *et al*., 2013).

Human cells lack Hsp104 and the Cdc10 homolog septin-9 lacks a CTD but at least in some circumstances septin-9 appears to persist stably outside of septin hetero-oligomers. For example, whereas in a human chronic myelogenous leukemia cell line ectopic expression of tagged septin-9 or depletion of endogenous septin-7 (the G dimer partner) results in disappearance of endogenous septin-9 molecules not incorporated into hetero-oligomers (Sellin *et al*., 2012), in an osteosarcoma cell line mutating the G interface of septin-9 results in the accumulation of what appear to be septin-9 monomers (Kuzmić *et al*., 2022). Mutating the Septin-7 G interface in mouse embryonic fibroblasts also generates apparent Septin-9 monomers (Abbey *et al*., 2016). Most septin-9 isoforms have extended N-terminal domains that may confer monomer stability. Alternatively, other mammalian chaperones might perform Hsp104’s septin-stabilizing function.

Cancer cells appear to exploit the homo-oligomerization properties of septin CTD coiled coils for other purposes: spontaneous fusions of the epidermal growth factor receptor (EGFR) to the coiled coil sequence in the CTD of human Septin-14 are the most frequently found functional gene fusion in glioblastoma (Frattini *et al*., 2013) and have also been found in colorectal cancer (Li *et al*., 2020) and lung adenocarcinoma (Zhu *et al*., 2019). Artificial EGFR dimerization via fusion to a leucine zipper is sufficient to activate its signaling activity in a ligand-independent manner (Kourouniotis *et al*., 2016). Hence, fusion to the Septin-14 sequence presumably drives EGFR homodimerization. Notably, Septin-14 is not predicted to homodimerize within the septin hetero-octamers that comprise septin filaments in non-cancer cells (Wang *et al*., 2025), but purified Septin-14 CTD can indeed form coiled-coil homodimers in vitro (Cavini *et al*., 2023). Whether transient Septin-14 coiled-coil homo-oligomers play a role in septin hetero-octamer assembly is an interesting question for future research.

## METHODS

### Strains and plasmids

Plasmids for *E. coli* expression of Cdc12 (Uniprot:P32468) or Cdc3 (Uniprot:P32457) or fragments thereof were made by PCR amplification from full-length synthetic genes with optimized codons for *E. coli* (Synbio). A tryptophan residue was inserted at the N terminus of Cdc3 CTD constructs. For CTD fragments, the amplified DNA fragment was purified and subcloned using restriction enzymes (*Nde*I and *Xho*I) into pET28a(+) vector (Novagen), yielding a sequence with an N-terminal H_6_ tag and a thrombin cut site. For MBP-fused Cdc12 constructs, the insert with *BamH*I and *Xho*I terminal sites was cloned into pOPINM vector (Novagen) with the In-fusion HD cloning kit (Takara), resulting in a coding sequence with a H_6_ tag and MBP at the N-terminus and a tripeptide linker (SGS) between the MBP and the Cdc12 sequence. Site-directed mutagenesis of *E. coli* expression plasmids was performed using non-overlapping primers (NEBaseChanger, New England BioLabs) and the mutations were confirmed by Sanger sequencing.

The following plasmids were used for yeast expression. pMVB2 (Versele *et al*., 2004) was used to express Cdc12-GFP under control of the *GAL1/10* promoter, and derivatives thereof were previously published (pMVB160, encoding Cdc12ΔCTD-GFP (Versele *et al*., 2004)), made via site-directed mutagenesis by Keyclone Technologies (C6F1B99E, encoding Cdc12(N379D W367K)-GFP; BD7C264E, encoding Cdc12(N379D E368V)-GFP), 9F2B6E98, encoding Cdc12(N379D W367K E368V)-GFP)) or GenScript (10633A79, encoding Cdc12(W367K)-GFP, with the N379D mutation repaired). pMVB169 (*ARS/CEN URA3 P_GAL1/10_–CDC3*) is based on plasmid TS395 (Carminati and Stearns, 1997) with the *CDC3* ORF cloned into the *Bam*HI site but has no GFP, and was constructed by Matthias Versele while in the lab of Jeremy Thorner. D2AC2EB4, encoding Cdc12-GFP(β1-9), was made by recombination in yeast: pMVB2 digested with *Xba*I and *Not*I was co-transformed into strain BY4742 with a PCR product made from template plasmid pGF-IVL794 (*GAL1/10* promoter driving the fragment of GFP(β1-9) (Finnigan *et al*., 2016)) with primers Cdc12-GFP(1-9)fw and Cdc12-GFP(1-9)re. Primer sequences are provided in Table S5. 15C00ABB, encoding Cdc12(W367K)-GFP(β1-9) was made by GenScript via site-directed mutagenesis of template plasmid D2AC2EB4. 76068B19, encoding Cdc12-GFP(β10), was made by recombination in yeast: pLP29 (Lippincott and Li, 2000) was digested with *Bam*HI and *Xba*I and co-transformed into BY4741 with annealed primers Cdc12-GFP(10)fw and Cdc12-GFP(10)re. E2438448, encoding Cdc12(W267A)-GFP(β11), was made by recombination in yeast: YCpL-Cdc12W267A (McMurray *et al*., 2011), a derivative of pMVB45 (Versele *et al*., 2004), was linearized with *Bam*HI and co-transformed into BY4741 with annealed and extended oligos Cdc12-GFP(11)fw and Cdc12-GFP(11)reFIXED. pRS313 and pRS316 have been described previously (Sikorski and Hieter, 1989), as have *HSP104* plasmids 5679 and 5707 (Schirmer *et al*., 2004). Plasmid 1EB21EA6 encodes Cdc12(G268R W367K)-GFP under *GAL1/10* promoter control and was made by GenScript via site-directed mutagenesis from template plasmid 10633A79. Plasmid YEp-Cdc12(G268R W367K)-GFP encodes Cdc12(G268R W367K)-GFP under *CDC12* promoter control was made by recombination in yeast: pMVB62 (2 µm origin, *CDC12-GFP HIS3* (Versele *et al*., 2004)) was digested with *Mlu*I and *Hpa*I to excise a portion of the *CDC12* coding sequence and transformed into BY4741 yeast cells carrying plasmid 1EB21EA6. pMETp-CDC10-GFP and the derivative encoding Cdc10 Δ0 (pJT3456) have been described previously (McMurray *et al*., 2011), as has the *GAL1/10 CDC3-GFP URA3* plasmid pMVB1 (Schaefer *et al*., 2016), pGF-IVL-287 (*LEU2 P_GAL_–CDC11-mCherry*) and pGF-IVL-286 (*LEU2 P_GAL_–SHS1-GFP* (Takagi *et al*., 2021), and plasmids G01273 (*P_GAL_–CDC3-mCherry–CDC10-GFP*), E00432 (*P_GAL_–CDC3*Δ*NTD-V_C_–CDC10-V_N_*), and E00435 (*P_GAL_–CDC3*Δ*NTD-V_C_–CDC10-V_N_*) (Benson and McMurray, 2023). For expression of mNeonGreen-tagged MS2 coat protein (MCP) fused to Pab1, we used the MCP sequence from pWL078 (Li *et al*., 2022), the sequence of an established Pab1ΔRRM4 construct (Deardorff and Sachs, 1997) plus the F366V mutation, and the mNeonGreen sequence from pDRF1-GW ymNeongreen-T2A-mCherry (Botman *et al*., 2019). This sequence was inserted into a plasmid (pBID2-EP-8) with a promoter controlled by the synthetic transcription factor sTF42 (Rantasalo *et al*., 2016), creating plasmid 14D13F62.

The following yeast strains were used. Existing strains were BY4741, BY4742, and BY4743 (Winzeler *et al*., 1999); JTY4013 (BY4742 *hsp104*Δ*::kanMX* (Johnson *et al*., 2020)); H06799 (BY4741 *HSP104-mCherry* (Hassell *et al*., 2022)); H07151 (BY4741 *mCherry-CDC3::LEU2* (Benson and McMurray, 2023)); H06798 (BY4741 *CDC10-GFP::HIS3MX6* (Huh *et al*., 2003)), GFY-160 (BY4741 *CDC11-mCherry::Sphis5 SHS1-eGFP::natMX*) (Finnigan *et al*., 2015b), and CEN.PK2-1D (EUROpean Saccharomyces Cerevisiae ARchive for Functional Analysis (EUROSCARF) RRID:SCR_003093 #30000B). The *cdc10*Δ*::kanMX* strain JTY4010 (McMurray *et al*., 2011) is originally from the haploid deletion collection (Winzeler *et al*., 1999). The *pre1(S142F) hsp104*Δ strain F93544A5 was made by crossing JTY4013 (Johnson *et al*., 2015) with the “*pre1-1*” strain from the BY4741-based temperature-sensitive mutant collection (Li *et al*., 2011), inducing sporulation in the resulting diploid by plating on 1% potassium acetate agar medium, and dissecting the resulting tetrads. The *cdc3*Δ*::kanMX/CDC3* strain from the heterozygous deletion collection (Winzeler *et al*., 1999), which is also heterozygous for a deletion allele at the *MET25* locus, was transformed with pMVB169. Cells from a plasmid-selective liquid culture were plated onto galactose-containing lead nitrate medium (Cost and Boeke, 1996). Spontaneous recombination events in between rDNA sequences frequently lead to loss of heterozygosity of centromere-distal loci, including *MET25* and *CDC3*. A brown colony, presumed to be *met25*Δ*/met25*Δ *cdc3*Δ*/cdc3*Δ, was confirmed by plating to YPD (2% peptone, 1% yeast extract, 2% glucose) to require galactose for survival; the resulting strain is named MMY0296. The *cdc12(W367K) mCherry-CDC3* strain 4H9K2X7P was made by co-transforming the following DNA molecules into cells of strain H07151 that were already carrying plasmid pMVB2: *Bsa*I-cut pBBK03, which encodes *Streptococcus pyogenes* Cas9, carries a promoter for guide RNA expression, and is marked with a hygromycin-resistance marker (Bean *et al*., 2022); annealed and extended oligos CDC12gRNAfw and CDC12gRNAre, which encode a guide RNA targeting *CDC12* sequence AATCAGACTGAACGGTGACTTGG; and annealed and extended oligos cdc12W367KdonorFW and cdc12W367KdonorRE, which encode a repair template harboring the W367K substitution and synonymous codons at adjacent positions to block guide RNA binding. Fortuitously, the N379D mutation in pMVB2 also blocks guide RNA binding, hence the transformation was plated to hygromycin medium (Goldbio # H-270-1, used at 200 µg/mL) with 2% galactose and resulting colonies were screened for growth on hygromycin medium with 2% glucose to repress Cdc12-GFP expression from pMVB2. The 3’ end of the endogenous *CDC12* genes from six slow-growing clones were amplified by PCR using primers cdc12midfw and 3’cdc12re and subjected to Sanger sequencing with the cdc12midfw primer, confirming the presence of the W367K mutation in all six. The strains for Cdc12 mRNA localization were created by first integrating into CEN.PK2-1D *Nsi*I-cut plasmid B5830 (Rantasalo *et al*., 2016), which encodes a synthetic transcription factor (sTF42) under control of the *MET25* promoter, creating strain 18FE4ED7. Next, 18FE4ED7 was transformed with a PCR product from template plasmid pET264-pUC 24xMS2V6 Loxp KANr Loxp (a gift from Robert Singer & Evelina Tutucci (Addgene plasmid # 104393 ; http://n2t.net/addgene:104393 ; RRID:Addgene_104393) (Tutucci *et al*., 2018) using primers Cdc12-24xMS2V6fw and Cdc12-24xMS2V6re, introducing 24 copies of the MS2V6 sequence into the Cdc12 3’UTR. The kanamycin resistance gene was excised from this insertion by transient Cre recombinase expression using plasmid pBF3060 (a gift from Nancy DaSilva & Suzanne Sandmeyer (Addgene plasmid # 26853 ; http://n2t.net/addgene:26853 ; RRID:Addgene_26853) (Fang *et al*., 2011), creating strain 3308F197. Finally, *EcoR*V-cut plasmid 14D13F62 was integrated into strain 18FE4ED7 and strain 3308F197, creating strain 7M3Q9V2W (with no tagged mRNA) and 5B8N4K9X (with tagged Cdc12 mRNA). Synthetic medium for yeast growth was based on YC (0.1 g/L Arg, Leu, Lys, Thr, Trp, and uracil; 0.05 g/L Asp, His, Ile, Met, Phe, Pro, Ser, Tyr, and Val; 0.01 g/L adenine; 1.7 g/L Yeast Nitrogen Base without amino acids or ammonium sulfate; 5 g/L ammonium sulfate; 2% dextrose or galactose) with individual components eliminated as appropriate for plasmid selection, or methionine concentration increased to 1 mM for Cdc12 mRNA localization experiments. For solid media, agar was added to 2%.

### Peptide expression and purification

*E. coli* Rosetta(DE3) (Novagen) or T7Express (NEB) strains transformed with the respective plasmid were grown in LB medium containing the appropriate antibiotic(s) for selection. Culture was grown at 37 °C and when OD_600nm_ reached 0.6-0.8, isopropyl-β-D-thiogalactopyranoside (IPTG) was added to 0.2 mM to induce protein expression, which was carried out overnight at 18 °C. Cells were harvested by centrifugation, solubilized in buffer ‘2x’ (100 mM Tris pH 8.0, 1 M NaCl) containing 10 mM imidazole, and lysed by sonication. For Cdc12 constructs, the cell extract was cleared by centrifugation, applied to an immobilized metal affinity chromatograph (IMAC) column (Ni-NTA agarose, Qiagen or HisTrap HP, Cytiva) and washed exhaustively with buffer ‘2x’ plus 10 mM imidazole. An additional washing step (4 column volumes, buffer ‘2x’ plus 35 mM imidazole) was carried out prior to elution (6 column volumes, buffer ‘2x’ plus 200 mM imidazole). Bovine thrombin (Sigma Aldrich #T7326) was added to the eluted peptide in a 2 U/mg ratio and the solution was dialyzed overnight against buffer ‘1x’ (50 mM Tris pH 8.0, 500 mM NaCl) using a 3 kDa dialysis membrane (Spectrum labs), except for the MBP-fused constructs. Solution was loaded to a Benzamidine Sepharose 6B resin (GE Healthcare) to remove thrombin and later to a second metal affinity chromatography to remove Histag-containing fragments. Size-exclusion chromatography (SEC) was performed in buffer containing 50 mM Tris pH 8.0, 300 mM NaCl using Superdex 75 10/300 (for CTD and CC constructs) or Superdex 200 10/300 columns (for MBP-fused constructs) assisted by an AKTA FPLC system (GE Healthcare). Protein purity was analyzed either by 12% SDS-PAGE or 16% tricine-SDS-PAGE. Cdc3C (UniProt: P32457, residues 413– 520) and Cdc3CC (residues 427–503) were purified using the same general procedure, except that both proteins were obtained through a refolding protocol. The insoluble fraction of the cell lysate was resuspended in ‘2x’ buffer containing 6 M urea, and the clarified material was loaded onto a Ni–NTA agarose (Qiagen) column. The resin was subsequently washed with ‘2x’ buffer (urea-free) using a 60-minutes linear gradient on an FPLC system, corresponding to a decrease of 0.1 M urea per minute. Cdc3C and Cdc3CC were then eluted as usual with buffer ‘2x’ plus 200 mM imidazole.

### Circular dichroism (CD) spectroscopy

Far-UV circular dichroism (CD) experiments were conducted using a Jasco J-815 spectropolarimeter equipped with a Peltier temperature control module. Purified peptides were prepared at a concentration of 10 µM in a buffer consisting of 20 mM sodium phosphate, 50 mM NaCl, 10% glycerol, pH 7.5. Buffer exchange was performed with a sample dilution of at least 100-fold. Sample was placed in a 1 mm Suprasil quartz cuvette and CD spectra were collected at 4°C in a wavelength range from 195 to 260 nm, in 1 nm steps, a speed of 100 nm/min and a response time of 0.5 s. A series of six scans were collected, which were averaged and subsequently subtracted from the buffer contribution. Secondary structure content was calculated with BeStSel (Micsonai *et al*., 2025). Thermal denaturation experiments were monitored at 222 nm over a temperature range of 5 to 80°C, with data collected every 1°C and a temperature ramp rate of 1°C/min. Ellipticity values were normalized from zero (native state, low temperature) to one (denatured state, high temperature) using the measured ellipticities on each plateaus. Sigmoid curves were fitted with either a single or a double Boltzmann equation to obtain the CD melting temperature(s) (*T*_m_).

### Size-exclusion chromatography coupled to multi-angle light scattering (SEC-MALS)

Purified peptides at a concentration of 300 µM (∼2.6 and 3.4 mg/ml for Cdc12CC and Cdc12C constructs, respectively) were loaded into a Superdex75 10/300 GL column coupled to a Waters 600 Controller HPLC chromatograph. A 0.5 ml/min flow rate and a 50 mM Tris, 300 mM NaCl, pH 8.0 running buffer were used. Multi-angle light scattering (MALS) and the differential refractive index (dRI) of the eluted fractions were analyzed by miniDAWN-TREOS and Optilab T-rEX detectors (Waters Corp.), respectively. Light scattering coefficients obtained from BSA monomer peak were used in MALS detectors normalization. Data were analyzed with ASTRA 7 software.

### Size-exclusion chromatography coupled to small-angle X-ray scattering (SEC-SAXS)

SEC-SAXS experiments were performed at the B21 beamline (Diamond Light Source, UK) (Cowieson *et al*., 2020). The setup employed an Agilent 1200 HPLC system with a Superdex 200 increase 3.2/300 column (GE Healthcare). Samples were prepared at a concentration of 5 mg/ml and a volume of 45 μl was injected and analyzed at a flow rate of 0.075 ml/min in buffer containing 50 mM Tris, 150 mM NaCl, pH 7.5. Data collection was acquired at 15 °C with a wavelength of 0.9464 Å by an EigerX 4 M (Dectris) detector positioned at a fixed camera length of 3.7209 m. The angular *q* range recorded was from 0.0045 to 0.34 Å^−1^ with a photon flux of 4.1012 s^−1^.

For data analysis, ScatterIV (Tully *et al*., 2021), ATSAS (Manalastas-Cantos *et al*., 2021) and BioXTAS RAW (Hopkins *et al*., 2017) were used. Mass estimation was carried out through Bayesian inference (Hajizadeh *et al*., 2018), Size&Shape (Franke *et al*., 2018) and the volume of correlation methods (Rambo and Tainer, 2013). Atomistic modeling was carried out with MultiFoXS (Schneidman-Duhovny *et al*., 2016) and/or EOM-NNLSJOE (Bernadó *et al*., 2007; Tria *et al*., 2015) considering flexible linkers between the MBPs and the CC helices and using either dimers, trimers or an ensemble of both. The radius of gyration (*R*_g_) in an ensemble was calculated as a weighted root mean square of the individual *R*_g_ values. Ab initio bead density models were generated initially by DAMMIF (Franke and Svergun, 2009) averaged over 23 independent runs using DAMAVER (Volkov and Svergun, 2003), and later by a single run of DAMMIN (Svergun, 1999) in ATSAS online 3.2.1 using dummy atoms with 5 Å radius. Calculated and experimental curves were compared using the correlation map method (CorMap) with a *P*-value of 0.01 as threshold (Franke *et al*., 2015).

### Size-exclusion chromatography of yeast lysates

50-mL cultures in synthetic plasmid-selective medium were pelleted and the cells were washed with PBS, resuspended in 1 mL PBS and transferred to a 2-mL screw-cap microcentrifuge tube, and lysed by four rounds of vortexing for 1 min each following the addition of 0.5-mm zirconia/silica beads (Biospec Products #11079105z). The tubes were incubated on ice for 1 min between each round of vortexing. Lysates were clarified by centrifugation at 16,300 rcf for 10 min at 4°C. 0.5 mL of clarified lysate was separated in PBS on a HiLoad 16/60 Sephacryl S200 column (GE Healthcare) column run at 0.5 mL/min. The column had been previously calibrated with ∼1 mg thyroglobulin, ∼1.5 mg alcohol dehydrogenase from *S. cerevisiae*, and ∼2 mg albumin (from kit #MWGF1000, Sigma-Aldrich) dissolved together in 500 µL PBS (Fig. S7 A).

### BS3 chemical crosslinking

Bis(sulfosuccinimidyl)suberate (BS3) crosslinking agent (molecular weight, 572.43 Da; spacer arm, 11.4 Å, Thermo-Fisher Scientific #21580) was used to analyze the oligomeric state of samples. Buffer used was 50 mM sodium phosphate, 300 mM NaCl, pH 8.5. BS3 concentration used was 2.5 mM (25-fold excess, 100 µM peptide) and samples were incubated on ice for 2 hours and 30 minutes. Reactions were stopped by the addition of 50 mM Tris-HCl pH 8.0 and analysed by 16% tricine-SDS-PAGE.

### Structural model predictions and coiled-coil structural analysis

The sequence logo was generated with WebLogo (Crooks *et al*., 2004) from 249 UniRef50 amino acid sequences identified using Cdc12 as the query. Deep-learning structural predictions were performed using the AlphaFold3 server (Abramson *et al*., 2024). Model confidence was assessed by pLDDT (predicted local distance difference test) and pAE (predicted aligned error). Coiled-coil radii were obtained using SamCC-Turbo (Szczepaniak *et al*., 2021). Predictions were deposited in ModelArchive. The “Contacts” function in ChimeraX v1.8 with van der Waals overlap ≥ -0.40 Å was used to predict intermolecular contacts.

### Nanodifferential scanning fluorimetry (NanoDSF)

Intrinsic protein fluorescence was analysed by Prometheus Panta (Nanotemper Technologies). The excitation wavelength of 280 nm was used, and the emission was measured at 330 nm and 350 nm. The buffer used was 25 mM sodium phosphate, 150 mM NaCl, pH 7.0. Samples at 90 µM concentration (1 mg/ml) were loaded into nanoDSF capillaries (Nanotemper Technologies, #PR-C006) in duplicates and transferred to the Prometheus NT.48 apparatus. Samples were heated from 18 to 90 °C (unfolding phase) and 578 measurements were taken (every 0.125 °C). Data were analyzed with Prometheus PR Control software (NanoTemper Technologies).

### Immunoblotting of SEC fractions of yeast lysates

For analysis by SDS-PAGE, 0.5-mL fractions were collected in a deep-well 96-well plate. To each fraction, 100 µL of 100% trichloroacetic acid (TCA) was added and the plate was stored on ice in a 4°C room overnight. Fractions were transferred to microcentrifuge tubes and centrifuged at 13,000 rcf for 10 minutes at 4°C. Pellets were washed twice with cold acetone (0.5 mL each time) and air dried. After dissolving the pellets in 50 µL 1X Laemmli buffer (62.5 mM Tris-HCl (pH 6.8), 2% SDS, 10% glycerol, 0.005% bromophenol blue), the samples were heated for 5 min at 85°C and frozen at -70°C. Samples were thawed at 37°C and 20 µL or 30 µL was resolved by Tris-glycine SDS-PAGE with 4-15% gradient gels (Bio-Rad #4561083). Proteins were transferred to PVDF membranes (Bio-Rad #1620264), blocked with Intercept (TBS) Blocking Buffer (Li-Cor #927-60001), and detected with anti-GFP antibody (Sigma-Aldrich #11814460001 RRID:AB_390913) and an infrared-fluorophore-labeled secondary antibody (Life Technologies Corporation #W10815 RRID:AB_2556797). After fluorescence scanning, the membranes were stained with Amido Black 10B (Serva #12310).

For dot blotting, 1-mL fractions were collected and the entirety of each was applied to a pre-wetted nitrocellulose membrane (GE Water & Process Technologies #WP4HYB0010) via vacuum using a Hybri-dot manifold (Bethesda Research Laboratories #1050MM). One dot was loaded with PBS alone, as a background control. The membrane was air dried, incubated in TBS for 5 min, and then blocked by incubation in Intercept (TBS) Blocking Buffer. The membrane was processed as described above using the same anti-GFP antibody or an anti-Hsp104 antibody (Enzo Life Sciences #ADI-SPA-1040-D RRID:AB_2039208) and secondary antibody Invitrogen #35569 RRID:AB_1965957 to detect the Hsp104 antibody. Signal intensities were quantified using FIJI (Schindelin *et al*., 2012) using circular regions of interest slightly larger than the dot size. The signal intensity of the PBS background control dot was subtracted from each, and the background-corrected signals were normalized to that of the dot with the highest signal.

### Immunoblotting of yeast total protein extracts

For analysis of total protein (unfractionated), protein was extracted from yeast cultures grown to mid-or late-log phase. Cell pellets were washed with 10% TCA and frozen in a dry ice/ethanol bath then thawed in 150 μL of 1.85 M NaOH and 7.4% β-mercaptoethanol. 150 μL of 100% TCA was added before incubation on ice for 10 min. Precipitated proteins were pelleted at 4°C at maximum speed for 10 min, washed twice with cold acetone (1 mL each time), and air dried. Pellets were resuspended in 75 μL of 0.1 M Tris base, 5% SDS. After addition of 25 µL 4X Laemmli buffer, the samples were boiled for 5 min and frozen at -20°C. Samples were thawed at 55°C and either resolved by SDS-PAGE as described above, or 3 uL aliquots were spotted on nitrocellulose membranes. Dot-blot membranes were blocked using Intercept (TBS) Blocking Buffer and then submerged for 1 hr in antibodies (rabbit anti-Glucose-6-Phosphate Dehydrogenase (Zwf1), Sigma-Aldrich #A9521, RRID AB_258454, or rabbit anti-mCherry, EnCor Cat# RPCA-mCherry, RRID AB_2571870) diluted 1:1,000 in blocking buffer. Membranes were washed thrice with TBST and submerged for 1 hr in a 1:1000 dilution of Peroxidase Conjugated Goat Anti-Rabbit IgG (ThermoScientific #32460 RRID:AB_1185567) in blocking buffer. After three TBST washes, the membranes were submerged in SuperSignal™ West Pico Chemiluminescent Substrate (Pierce #34080) for 10 minutes prior to imaging with a Bio-Rad Chemidoc MP imaging system. Quantification was performed in FIJI by calculating the integrated density for each spot using a 118×120-pixel oval area. Values for two empty areas of the same size on the same membrane were averaged and subtracted from the values for each sample on the same membrane. Each background-corrected value for mCherry was then normalized to that of Zwf1. Finally, the Zwf1-normalized values were normalized to that of the *CDC3-mCherry* sample (strain H07151).

### smiFISH

smiFISH was performed using Stellaris RNA FISH buffers (SMF-HB1, SMF-WA1, SMF-WB1, Biosearch Technologies) essentially as described previously (Tsanov *et al*., 2016) but modified for budding yeast cells according to Stellaris instructions. Briefly, fixed cells were prepared from 50-mL YPD cultures grown to an OD_600_ of 0.2-0.5 and pelleted at 1600 x *g* for 4 minutes in a centrifuge. Cells were resuspended in 1 mL ice-cold methanol, transferred to a 1.5-mL microcentrifuge tube, and incubated at -20°C for 10 minutes. Following another pelleting step and two washes with 1 mL ice-cold Fixation buffer (1.2 M sorbitol, 0.1 M potassium phosphate), cells were resuspended in 1 mL of Fixation buffer plus 2.5 µL Zymolyase 20T (#Z1000, US Biologicals) 2.5 mg/mL dissolved in water). Cell wall digestion was performed at 30°C for ∼2 hr then cells were pelleted at 1,000 x *g* for 3-5 minutes and washed twice with ice-cold Fixation buffer. Cells were permeabilized by resuspending in 1 mL 70% ethanol and stored at 4°C prior to hybridization. For hybridization, 300 µL of fixed yeast cells in 70% ethanol were pelleted at 400 x *g* and resuspended in 100 μL Hybridization Buffer containing probes, then incubated in the dark at 30°C overnight. 100 µL of Wash Buffer A was added and cells were pelleted at 400 x *g* for 5 minutes. Cells were resuspended in 1 mL of Wash Buffer A and incubated in the dark at 30°C for 30 minutes. After pelleting again at 400 x *g* for 5 minutes, cells were resuspended in 1 mL of DAPI stain (Wash Buffer A consisting of 5 ng/mL DAPI) and incubated in the dark at 30°C for 30 minutes. Cells were pelleted and resuspended in 1 mL of Wash Buffer B for 5 minutes, then pelleted and resuspended in a small drop of Vectashield Mounting Medium (Vector Laboratories, #H-1000). 5-10 µL of Vectashield-suspended yeast cells were spotted on a glass microscope slide and then covered with an 18 × 18 mm square #1 coverglass. The coverglass was pressed onto the slide and excess mounting medium was removed with a laboratory wipe. The coverglass perimeter was sealed with clear nail polish. All probes were purchased from Integrated DNA Technologies. Primary probes (13-19 per mRNA) were designed manually; sequences used are provided in Table S5. Primary probes were combined in equimolar amounts and diluted to 20 µM and hybridized with the fluorescent FLAP probe. 3.125 µL of hybridized probe was added to 100 µL of Hybridization buffer. For the negative control for the Cdc3 probe set, strain MMY0296 was cultured in plasmid-selective medium containing 2% raffinose overnight and then glucose was added to 2% final concentration to block new *CDC3* transcription. The cells were then fixed after 6 hr at 30°C. The *cdc10*Δ*::kanMX* strain is known to be as large as wild-type diploid cells (Jorgensen *et al*., 2002).

### Microscopy

All microscopy was performed at room temperature (22°C). Most micrographs used cells spotted from liquid suspension onto 1% agarose pads and were captured with an EVOSfl all-in-one epifluorescence microscope (Thermo-Fisher Scientific) equipped with a 60×/1.25 NA oil objective objective, GFP (#AMEP4651) and Texas Red (#AMEP4655) LED/filter cubes, and Sony ICX285AL Interline CCD 1360×1024-pixel camera. SmiFISH slides were imaged using a widefield DeltaVision Microscope equipped with 60×/1.42 NA oil objective, TRITC and Cy5 filter sets, and a CoolSnap ES2 1392×1040-pixel camera, using consistent laser intensity and exposure times across samples. DeltaVision images were deconvoluted with SoftWorx (Applied Precision, Inc).

Maximum-intesity projections (of 36 z slices 0.2 μm apart) and other image adjustments were performed using FIJI software (Schindelin *et al*., 2012). Cellular length-to-width ratios were calculated with FIJI line tool as described previously (Benson and McMurray, 2023). Cdc12 mRNA foci were determined to be in the mother-proximal or -distal half of the bud by measuring in FIJI the location of the center of each focus on a line drawn from the bud neck to bud tip along the mother-bud axis.

## ONLINE SUPPLEMENTAL MATERIAL

Table S1 shows SEC-SAXS data and analysis. Table S2 shows melting temperatures of Cdc12C wild type and mutants monitored by circular dichroism. Table S3 shows circular dichroism spectral analysis of wild-type and mutant purified Cdc12 CTD. Table S4 shows calculated mass of peptides in the SEC-MALS experiments. Table S5 shows sequences of select oligonucleotides used in this study. Fig. S1 shows AlphaFold Multimer modeling metrics and output. Fig. S2 shows SEC-SAXS of MBP-Cdc12CC. Fig. S3 shows SEC-SAXS of MBP-Cdc12CC+8. Fig. S4 shows the radius of gyration (Rg) from MBP-Cdc12CC+8 assemblies generated by RANCH. Fig. S5 shows SAXS analyses of MBP-Cdc12CC+8 SEC fractions. Fig. S6 shows AlphaFold3 predictions of Cdc12•GDP homodimers. Fig. S7 shows data from size exclusion chromatography of yeast lysates overexpressing Cdc12-GFP or mutants thereof. Fig. S8 shows post-translational assembly into septin filaments by Cdc12-GFP and Cdc12(E368V)-GFP. Fig. S9 shows effects of mutations altering Cdc12 CTD oligomerization state on septin function caused by overexpression of GFP-tagged Cdc12 in cells with mCherry-tagged Cdc3. Fig. S10 shows effects of the N379D mutation discovered in the Cdc12-GFP-overexpressing plasmid. Fig. S11 shows AlphaFold3 predictions of Hsp104–substrate binding. Fig. S12 shows effects on colony growth, cellular morphology, and/or septin localization of overexpression of septins and/or Hsp104. Fig. S13 shows levels of tagged septins quantified by immunoblotting. Fig. S14 shows experimental controls for smiFISH detection of septin mRNAs. Fig. S15 shows AlphaFold3 predictions of Cdc12 TriFC complexes.

## DATA AVAILABILITY

All data supporting the findings of this study are available within the paper and its supplemental materials, including source data files. SEC-SAXS datasets and analysis are available in SASBDB (sasbdb.org) with the respective accession codes: MBP-Cdc12 CTD 332-407aa “MBP-Cdc12CC+8” (SASDYR4), MBP-Cdc12CC+8 peak front (SASDYS4), MBP-Cdc12CC+8 peak tail (SASDYU4) and MBP-Cdc12 CTD 340-407aa “MBP-Cdc12CC” (SASDYQ4).

The following AlphaFold structural predictions are available in ModelArchive (modelarchive.org) with the respective accession codes: Hsp104 N-terminal domain/Cdc12 WALTZ amyloidogenic sequence (ma-k1rre), Hsp104 N-terminal domain/Cdc12 C-terminal domain (ma-gmeo1), Hsp104 N-terminal domain/full-length Cdc12 (ma-huwgf), Hsp104 N-terminal domain/Cdc10 WALTZ amyloidogenic sequence (ma-5n6qq), Hsp104 N-terminal domain/Sup35 binding site (ma-vj5d5), Cdc12-Cdc3 heterodimeric coiled coil (ma-t43av), Cdc12 homotrimeric coiled coil (ma-nmv5d), shorter Cdc12 homotrimeric coiled coil (ma-evurw), full-length Cdc12 dimer with GDP (ma-5hhc4), full-length Cdc12 trimer with GDP (ma-99kcr), full-length Cdc11-3xCdc12-Cdc3-Cdc10 TriFC complex (ma-37ka2) and full-length 3xCdc12 TriFC complex (ma-s4slk).

## ACKNOWLEDGEMENTS

We thank Prof. Matthew P. Crump (University of Bristol, UK) for providing SEC-SAXS beamtime, B21 beamline specialists (Diamond Light Source, UK) and Daniel Frank (BIOSAXS GmbH, Germany) for the assistance with SEC-SAXS data acquisition/analysis, the Integrated Structural Biology Platform of Carlos Chagas Institute and Dr. Beatriz Gomes Guimarães (Fiocruz Paraná, Brazil) for the use of and help with the nanoDSF/Prometheus Panta, Matt Taliaferro (University of Colorado School of Medicine) for use of the Deltavision microscope, Lenny Teytelman (Protocols.io) for advice with smiFISH, and Allison McClure (University of Colorado School of Medicine) for FPLC use. We gratefully acknowledge the National Institutes of Health (grants T32 GM136444 and R35 GM148198), Fundação de Amparo à Pesquisa do Estado de São Paulo (grants 2015/16812-0, 2018/19992-7, 2020/02897-1, 2022/00262-4 and 2024/23432-8), and a pilot grant from the department of Cell and Developmental Biology at the University of Colorado School of Medicine for financial support. The funders had no role in study design, data collection and analysis, decision to publish or preparation of the manuscript. The authors declare no competing financial interests.

## AUTHOR CONTRIBUTIONS

Italo A. Cavini: conceptualization, methodology, validation, formal analysis, investigation, visualization, project administration, writing – original draft, review, and editing. Randi M. Yeager: methodology, investigation, visualization. Alejandra Velasquez: investigation, visualization. Andressa P.A. Pinto: methodology, investigation, formal analysis. Ana P.U. Araujo: conceptualization, supervision, funding acquisition, writing – review and editing. Richard C. Garratt: conceptualization, project administration, funding acquisition, writing – review and editing. Michael McMurray: conceptualization, methodology, investigation, formal analysis, validation, visualization, supervision, project administration, funding acquisition, writing – original draft, review, and editing.

## Notes

### Competing Interest Statement

The authors have declared no competing interest.

### Summary of Updates

Section on Cdc12(W367K) bud neck localization in hsp104∆ cells updated to clarify that this is another scenario where the mutant protein is able to incorporate into septin filaments. Section on Cdc12 mRNA localization (and corresponding section in Methods) updated to note that most bright foci in buds were in the portion of the bud distal from the mother cell. Figure 3 was updated to include missing molecular weight marker size. Supplemental files were also updated to provide missing scale bars and molecular weight marker sizes.

