## Supplemental files for "Coiled-coil homo-oligomerization and disaggregase Hsp104 act in parallel to stabilize orphan septins"

Table S1 SEC-SAXS data and analysis.

| **SASBDB entry** | **SASDYR4** | **SASDYS4** | **SASDYU4** | **SASDYQ4** |
| --- | --- | --- | --- | --- |
| **Sample** | **MBP-**  **Cdc12CC+8** | **MBP- Cdc12CC+8 (peak front)** | **MBP- Cdc12CC+8 (peak tail)** | **MBP-**  **Cdc12CC** |
| **Instrumental** |  |  |  |  |
| Instrument and SEC parameters: | beamline, B21 Diamond Light Source; frames, 600 total (one every 3 s);  column, Superdex 200 Increase 3.2/300; flow rate, 0.075 ml/min;  concentration at injection, 5.0 mg/ml; injection volume, 45 μl;  buffer, 50 mM Tris-HCl, 150 mM NaCl, pH 7.5 | | | |
| Selected frames (total number of frames) | 337-350 (14) | 345-353 (9) | 392-407 (16) | 364-373 (10) |
| **Guinier analysis** |  |  |  |  |
| *I*(0) ± *σ*, cm^-1^ | 0.0943 ± 0.0001 | 0.0951 ± 0.0002 | 0.0252 ± 0.0001 | 0.0823 ± 0.0002 |
| *R*_g_ ± *σ*, Å | 47.79 ± 0.09 | 48.6 ± 0.2 | 44.2 ± 0.2 | 44.3 ± 0.2 |
| *q*_min_ (calculated *q*_min_ max, π/*D*_max_), Å^-1^ | 0.0151 (0.0180) | 0.0149 (0.0180) | 0.0164 (0.0187) | 0.0139 (0.0188) |
| *q*.*R*_g_ max | 1.2936 | 1.2530 | 1.2945 | 1.1294 |
| *r*^2^ | 0.9988 | 0.9957 | 0.9904 | 0.9953 |
| **PDDF/*P*(*r*) analysis** |  |  |  |  |
| *I*(0), cm^-1^ | 0.0957 ± 0.0001 | 0.0949 ± 0.0004 | 0.0256 ± 0.0001 | 0.0825 ± 0.0002 |
| Real space *R*_g_, Å | 49.3 ± 0.2 | 49.4 ± 0.2 | 46.4 ± 0.3 | 45.2 ± 0.1 |
| *D*_max_, Å | 175 | 175 | 168 | 167 |
| *q* range^*^, Å^-1^ | 0.0151-0.2589 | 0.0149-0.2000 | 0.0164-0.2150 | 0.0139-0.2394 |
| *χ*^2^ (total estimate from GNOM) | 1.037 (0.82) | 1.059 (0.84) | 0.977 (0.71) | 1.336 (0.72) |
| Porod volume *V*_p_, 10^3^ Å^-3^ | 335 | 345 | 256 | 262 |
| Resolution from SASRES, Å | 40.39 | 47.54 | 42.46 | 45.74 |
| DATCLASS shape | flat | flat | flat | flat |
| AMBIMETER score | 2.91 | 2.91 | 2.77 | 2.70 |
| **Molecular mass determination (ratio to monomeric mass)** | | | | |
| From chemical composition, kDa | 51.01289 | 51.01289 | 51.01289 | 50.10885 |
| From Bayesian inference, kDa | 138.2 (2.71) | 146.8 (2.88) | 109.1 (2.14) | 109.1 (2.18) |
| From Size&Shape, kDa | 166.0 (3.25) | 159.1 (3.12) | 111.6 (2.19) | 110.4 (2.20) |
| From volume of correlation *V*_c_, kDa | 137.1 (2.69) | 146.1 (2.86) | 104.6 (2.05) | 99.1 (1.98) |
| **Shape model fitting** |  |  |  |  |
| *DAMMIF (fast, default parameters, P1 symmetry, no anisotropy assumptions, 23 reconstructions):* | | | | |
| NSD (standard deviation) | 0.89 (0.17) | 1.00 (0.17) | 1.47 (0.31) | 1.10 (0.21) |
| *DAMMIN (expert mode, r_dummy atoms_ = 5 Å):* | | | | |
| *χ*^2^, CorMap *P*-value | 1.036, 0.583 | 1.061, 0.278 | 0.980, 0.045 | 1.333, 0.013 |
| **Atomistic modelling** |  |  |  |  |
| *MultiFoXS (parallel dimer, single state, flexible linkers*^†^*, 10,000 iterations):* | | | | |
| *R*_g_, Å | 45.53 | 45.75 | 43.78 | 42.93 |
| *c*_1_, *c*_2_ | 1.04, 4.00 | 1.04, 3.24 | 0.99, 0.04 | 1.00, 2.02 |
| *χ*^2^, CorMap *P*-value | 2.17, 0.000 | 1.20, 0.005 | 1.28, 0.006 | 1.41, 0.051 |
| *MultiFoXS (parallel trimer, single state, flexible linkers*^†^*, 10,000 iterations):* | | | | |
| *R*_g_, Å | 47.65 | 48.36 | 48.16 |  |
| *c*_1_, *c*_2_ | 0.99, -0.50 | 0.99, -0.50 | 0.99, -0.50 |  |
| *χ*^2^, CorMap *P*-value | 7.91, 0.000 | 4.91, 0.000 | 16.4, 0.000 |  |
| *EOM-NNLSJOE (MBP free, CC fixed*^ψ^*, flexible linkers*^⁋^*, constant subtraction allowed):* | | | | |
| No. of representative structures | 9 | 8 | 6 |  |
| Oligomer percentage | 34.0% dimers,  66.0% trimers | 46.6% dimers,  53.4% trimers | 93.0% dimers,  7.0% trimers |  |
| Ensemble-averaged *R*_g_, Å | 47.70 | 47.92 | 44.74 |  |
| *χ*^2^, CorMap *P*-value | 1.07, 0.203 | 1.01, 0.940 | 0.97, 0.310 |  |

^*^Also used for mass determination, model fitting and atomistic modelling.

^†^Five-residues linker, 378-382 aa for MBP-Cdc12CC, 382-386 aa for MBP-Cdc12CC+8.

^⁋^Eleven-residues linker, 379-389 aa for MBP-Cdc12CC+8.

^ψ^Parallel trimeric CC from AlphaFold (5000 models) and parallel dimeric CC generated from trimer (5000 models).

Table S2 Melting temperatures of Cdc12C wild type and mutants monitored by circular dichroism at 222 nm. Peptide concentration, 10 µM. Values were taken from duplicates.

| **Peptide** | ***T*_m_, °C** | **Δ*T*_m_, °C** | **Number of states considered during fit** | **Transition width, °C** |
| --- | --- | --- | --- | --- |
| Wild type | 31.2 ± 0.1 | --- | 2 | 5.1 ± 0.1 |
| K391N-L392stop (mutant -6) | ND | --- | --- | 14 ± 1 |
| W367K | ND | --- | --- | 8.1 ± 0.3 |
| W367K-E368V | 23.6 ± 0.1 | -7.4 ± 0.2 | 2 | 5.6 ± 0.1 |
| R363K (mutant -122) | 30.3 ± 0.1 | -0.9 ± 0.2 | 2 | 5.9 ± 0.1 |
| W367L | 33.8 ± 0.1 | 2.6 ± 0.2 | 2 | 5.2 ± 0.1 |
| E368K | 31.7 ± 0.1 | 0.5 ± 0.2 | 2 | 6.1 ± 0.1 |
| E368Q | 34.4 ± 0.1 | 3.2 ± 0.2 | 2 | 6.1 ± 0.1 |
| E368V | 55.4 ± 0.2 | 24.2 ± 0.3 | 3^*^ | 4.8 ± 0.2 |

^*^Second transition at *T*_m_ = (27 ± 3) °C.

Table S3 Circular dichroism spectral analysis of wild-type and mutant purified Cdc12 CTD. The 222/208 nm ratio, indicative of coiled-coil assemblies, and the secondary structure determination by BeStSel (Micsonai *et al.* 2025) are presented. Peptide concentration, 10 µM. Temperature, 4 °C.

| **Peptide** | **222/208 nm ratio ± 0.04** | **Secondary structure determination, %** | | | |
| --- | --- | --- | --- | --- | --- |
|  |  | **Regular helix** | **Distorted helix** | **Helices**  **(regular+**  **distorted)** | **Other** |
| Wild type | 0.99 | 70.6 | 22.9 | 93.5 | 6.5 |
| K391N-L392stop (mutant -6) | 0.72 | 20.2 | 11.1 | 31.2 | 68.8 |
| W367K | 0.83 | 29.7 | 14.2 | 44.0 | 56.0 |
| W367K-E368V | 0.88 | 37.6 | 17.7 | 55.3 | 44.7 |
| R363K (mutant -122) | 1.02 | 54.9 | 18.9 | 73.9 | 26.1 |
| W367L | 0.96 | 58.2 | 22.5 | 80.7 | 19.3 |
| E368K | 0.99 | 62.6 | 21.9 | 84.5 | 15.5 |
| E368Q | 0.96 | 67.9 | 21.1 | 89.0 | 11.0 |
| E368V | 0.93 | 46.0 | 17.6 | 63.5 | 36.5 |

Table S4 Calculated mass of peptides in the SEC-MALS experiments.

| **Peptide** | **Calculated mass, kDa**  **(ratio to monomer)** |
| --- | --- |
| Cdc3 |  |
| 413-520 (Cdc3C) |  |
| wild type | 12.7 ± 0.9 (1.0) |
| 427-520 (Cdc3CC) |  |
| wild type | 9.6 ± 0.7 (1.0) |
| Cdc12 |  |
| 316-407 (Cdc12C) |  |
| wild type | 29.0 ± 1.3 (2.6) |
| K391N-L392stop (mutant -6) | 9.9 ± 0.1 (1.1) |
| W367K | 12.4 ± 0.3 (1.1) |
| W367K-E368V | 16.2 ± 1.6 (1.4) |
| R363K (mutant -122) | 24.6 ± 0.6 (2.2) |
| W367L | 25.5 ± 0.7 (2.3) |
| E368K | 22.0 ± 0.3 (1.9) |
| E368Q | 28.2 ± 0.7 (2.5) |
| E368V | 35.2 ± 0.6 (3.1) |
| 341-407 (Cdc12CC) |  |
| wild type | 22.2 ± 1.0 (2.6) |

Table S5 Sequences of select oligonucleotides used in this study.

| **Primer name** | **Sequence (5’ to 3’)** |
| --- | --- |
| Cdc12-GFP(1-9)fw | AACTAGAAGAGCAGGTCAAAAGCTTGCAAGTAAAAAAATCCCATTTAAAAATGAGAAAGGGTGAAGAATTGTTTACTG |
| Cdc12-GFP(1-9)re | AGAGAGAGGTACATACATAAACATGCGCGCACAAAAGCAGAGATTATCTATCATGAACCGTTGTCAGGTAATAAGAC |
| Cdc12-GFP(10)fw | AACTAGAAGAGCAGGTCAAAAGCTTGCAAGTAAAAAAATCCCATTTAAAAGATTTGCCAGATGATCATTATTTGTCTACTCAAACTATTTTGTCTAAAGA |
| Cdc12-GFP(10)re | GTTGTAAAACGACGGCCAGTGAATTCGAGCTCGGTACCCGGGGATCCTCAATTCAAATCTTTAGACAAAATAGTTTGAGTAGACAAATAATGATCATCTG |
| Cdc12-GFP(11)fw | AAGAGCAGGTCAAAAGCTTGCAAGTAAAAAAATCCCATTTAAAAGGATCCGAAAAAAGAGATCATATGGTTTTGTTGGAATATGTTACTGCTGCTGGTAT |
| Cdc12-GFP(11)reFIXED | AGTCACGACGTTGTAAAACGACGGCCAGTGAATTCGAGCTCGGTACCCGGCTAAGAAGCATCAGTAATACCAGCAGCAGTAACATATTCCAACAAAACCA |
| CDC12gRNAfw | TCCTCGCTGGCGCCGGCTGGGCAACACCTTCGGGTGGCGAATGGGACTTTAATCAGACTGAACGGTGACT |
| CDC12gRNAre | GTTGATAACGGACTAGCCTTATTTTAACTTGCTATTTCTAGCTCTAAAACAGTCACCGTTCAGTCTGATT |
| cdc12W367KdonorFW | TATTTCACTGACCAAGTCAAAGCTGAGGAGCAAAGATTTAGACAAAAGGAGCAGAATATTGTCAATGAAAGAATTAGATTGAATGGTGAT |
| cdc12W367KdonorRE | TTTTTTTACTTGCAAGCTTTTGACCTGCTCTTCTAGTTTTTTTACCTTTCCTTGAATTTCTTCTAAATCACCATTCAATCTAATTCTTTC |
| cdc12midfw | TTTCACGCCTCCATTAGACG |
| 3’cdc12re | ACGAGACAAAGAGGAAGACA |
| Cdc12-24xMS2V6fw | TCAAGGAAAGGTAAAAAAACTAGAAGAGCAGGTCAAAAGCTTGCAAGTAAAAAAATCCCATTTAAAATGACGCTCTAGAACTAGTGGATC |
| Cdc12-24xMS2V6re | GACAAAAGAAGAGACAAAGAGATAGGCGTTGAAATTGACGAGACAAAGAGGAAGACATTAATTAATCACACTAGTGGATCTGATATCACC |
| FLAP-Y Cy3 | Cy3-AATGCATGTCGACGAGGTCCGAGTGTAA-Cy3 |
| FLAP-Y Cy5 | Cy5-AATGCATGTCGACGAGGTCCGAGTGTAA-Cy5 |
| Cdc10_FISHa | CAGGCTGTACTGAGCTGAGAGGATCCttacactcggacctcgtcgacatgcatt |
| Cdc10_FISHc | CTTCAACAGACGATGTTCGATCTGttacactcggacctcgtcgacatgcatt |
| Cdc10_FISHe | GAGTACTTTTACCCAATCCGGttacactcggacctcgtcgacatgcatt |
| Cdc10_FISHg | CAGGCAGGGCAGAAATATCATCACCAGttacactcggacctcgtcgacatgcatt |
| Cdc10_FISHh | CAAGCGAACGCGGTCCTCCACAAGAGttacactcggacctcgtcgacatgcatt |
| Cdc10_FISHj | CACAATAGGCTCCCAAGCTTTAGttacactcggacctcgtcgacatgcatt |
| Cdc10_FISHl | CACGTTGGGCTGTCAATTCTTTACGttacactcggacctcgtcgacatgcatt |
| Cdc10_FISHo | CAAGGCTTCAACGTCAAGGCGGCTCttacactcggacctcgtcgacatgcatt |
| Cdc10_FISHq | GCTCCCTAAACTCCGTTCTTTCATCttacactcggacctcgtcgacatgcatt |
| Cdc10_FISHs | CCAACCACTGCAAACGGAATGATAGATCttacactcggacctcgtcgacatgcatt |
| Cdc10_FISHu | GCGCTCCAACGAGTTTTTCTTCCCCTGttacactcggacctcgtcgacatgcatt |
| Cdc10_FISHv | CACACTGGTTGATATCCTCAACttacactcggacctcgtcgacatgcatt |
| Cdc10_FISHw | GGAGATGAGTTCGAATCAAAAATTCCCttacactcggacctcgtcgacatgcatt |
| Cdc10_FISHx | GGCAATTAATTGTCTTGCTCTGAACCCttacactcggacctcgtcgacatgcatt |
| Cdc10_FISHy | GACATATGAGCTGAGGAACGACTATTCGCttacactcggacctcgtcgacatgcatt |
| Cdc10_FISHz | GAGATTCAACGTTGAATGGCGTTGCttacactcggacctcgtcgacatgcatt |
| Cdc3_FISHa | CGTTCAGTTGTCGTGTGATCCTGGttacactcggacctcgtcgacatgcatt |
| Cdc3_FISHc | CCTGCTTAATGGACACTTGTTCCTCCttacactcggacctcgtcgacatgcatt |
| Cdc3_FISHd | GATCATGCTGACGCTCTTCTTGTTCCGttacactcggacctcgtcgacatgcatt |
| Cdc3_FISHf | CGTGTACTGCGAGTCCACTCCATCATGttacactcggacctcgtcgacatgcatt |
| Cdc3_FISHg | CTCTCGCTGTCGTCATTTTGCGTACCGttacactcggacctcgtcgacatgcatt |
| Cdc3_FISHh | CCACCTTTACATCGGATTCAGCTGCCTCttacactcggacctcgtcgacatgcatt |
| Cdc3_FISHi | GGTGATGCCCATACCGAGGCCGGGttacactcggacctcgtcgacatgcatt |
| Cdc3_FISHj | GGTCAGGCAGAACTTGACCCTTTTCGCTttacactcggacctcgtcgacatgcatt |
| Cdc3_FISHk | GACGACGAATGAACTTAATCTCCGGttacactcggacctcgtcgacatgcatt |
| Cdc3_FISHm | CCGTTCTTTATGGACCTTCTGTGCCttacactcggacctcgtcgacatgcatt |
| Cdc3_FISHn | CCATCAGGGCCGACACATAGGAGttacactcggacctcgtcgacatgcatt |
| Cdc3_FISHp | GACCCTCGCCCTGACCCTCCttacactcggacctcgtcgacatgcatt |
| Cdc3_FISHs | CAATCTCCTTAATGATCGGGTCCCttacactcggacctcgtcgacatgcatt |
| Cdc3_FISHv | CCCCCCATGGATAGGAACGGCCttacactcggacctcgtcgacatgcatt |
| Cdc3_FISHy | CCCTTCTTCTTTGTAGGGACGGGGGttacactcggacctcgtcgacatgcatt |
| Ash1_FISHd | GATGTGGCGATGCTGGTGCAGttacactcggacctcgtcgacatgcatt |
| Ash1_FISHi | GAGACGGACGATAGCCTGTCttacactcggacctcgtcgacatgcatt |
| Ash1_FISHj | GGCTTTGTTGTGGGCGCTCCGGttacactcggacctcgtcgacatgcatt |
| Ash1_FISHl | GATGTATCAGGGAAGCGTGCTGCGttacactcggacctcgtcgacatgcatt |
| Ash1_FISHq | CGGCGGTGTGACGGGAGGAGttacactcggacctcgtcgacatgcatt |
| Ash1_FISHs | GAGGCCTCTACTCCTCTAAGCGttacactcggacctcgtcgacatgcatt |
| Ash1_FISHx | GCCTTGGGACGCACAGGTGAGGttacactcggacctcgtcgacatgcatt |
| Ash1_FISHy | GGGTGACCTTGGGCTTGGttacactcggacctcgtcgacatgcatt |
| Ash1_FISHz | GAGGGGGAGAGTCGAGAGCttacactcggacctcgtcgacatgcatt |
| Ash1_FISHaa | GAGCCACTGGACCTTCTACTTCCCttacactcggacctcgtcgacatgcatt |
| Ash1_FISHbb | CACTCTTGTGGTGTGACGAGTGGGttacactcggacctcgtcgacatgcatt |
| Ash1_FISHdd | GCTTTCTTGGCGACCACGATGGCCttacactcggacctcgtcgacatgcatt |
| Ash1_FISHgg | CGCAGTTGCGTTCGGGGACGttacactcggacctcgtcgacatgcatt |
| Abp140_FISHa | GGTAGCATCGCCTTCTTCCTttacactcggacctcgtcgacatgcatt |
| Abp140_FISHb | CCTCTTGCGGAACAGCAGTAGACTCttacactcggacctcgtcgacatgcatt |
| Abp140_FISHc | GGAGCCCTCTTCTTCGTCTTCTTCGttacactcggacctcgtcgacatgcatt |
| Abp140_FISHd | CTTGCAAGAACTTCAGCGCCAGttacactcggacctcgtcgacatgcatt |
| Abp140_FISHe | CCTTCAGTTTCCTCAGCCACACCttacactcggacctcgtcgacatgcatt |
| Abp140_FISHf | CTTCGGCTGGTTCAGATGCAttacactcggacctcgtcgacatgcatt |
| Abp140_FISHg | GCTGTCGTCGACATTAGCGTTGGTGttacactcggacctcgtcgacatgcatt |
| Abp140_FISHh | CGTGGTGTCGCCGGTTGTATCATCAttacactcggacctcgtcgacatgcatt |
| Abp140_FISHi | CACTTCTTGAATGGCGGAGGttacactcggacctcgtcgacatgcatt |
| Abp140_FISHj | CCTCTGCCCTCGTGCGCAATATTCTCttacactcggacctcgtcgacatgcatt |
| Abp140_FISHk | CTGCATTCTGATCGCCTGTGttacactcggacctcgtcgacatgcatt |
| Abp140_FISHl | GTCGCGGCCTATTCTTGAACCttacactcggacctcgtcgacatgcatt |
| Abp140_FISHm | GATCCCAAACGTCAGATTCCTCGGttacactcggacctcgtcgacatgcatt |
| Abp140_FISHn | CCTCCGCCTGCTGGACTTGTttacactcggacctcgtcgacatgcatt |
| Abp140_FISHo | GTCCCAATAACGTGCCGGATttacactcggacctcgtcgacatgcatt |
| Abp140_FISHp | CTAATTCGACGGCTCTTGGAGCttacactcggacctcgtcgacatgcatt |
| Abp140_FISHq | CCATACTGTTGCGTGGCCGTttacactcggacctcgtcgacatgcatt |
| Abp140_FISHr | GAGGTTCGACGCCATCAGGTAAttacactcggacctcgtcgacatgcatt |
| Abp140_FISHt | CAGCTTGTACCCAGCAGCGGttacactcggacctcgtcgacatgcatt |

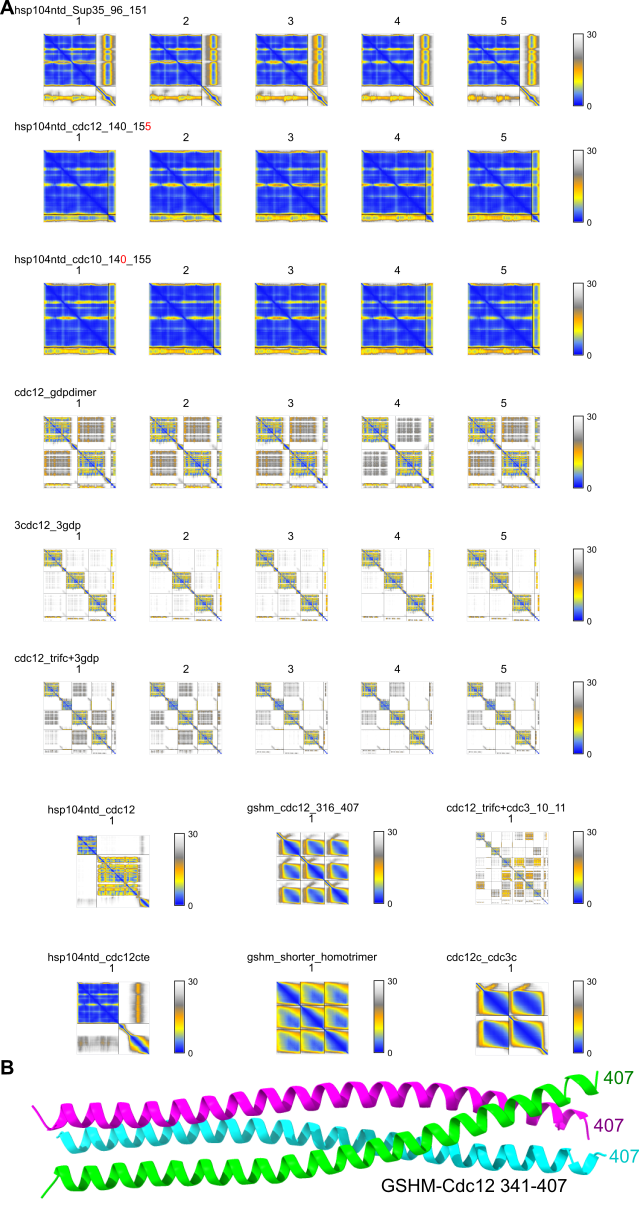

**Figure S1. AlphaFold Multimer modeling metrics and output.** (A) Predicted Aligned Error matrices for the AlphaFold3 predictions in this study. (B) Top-ranked AlphaFold3 prediction for Cdc12 residues 341-407 with residual N-terminal sequence GSHM from protease cleavage.

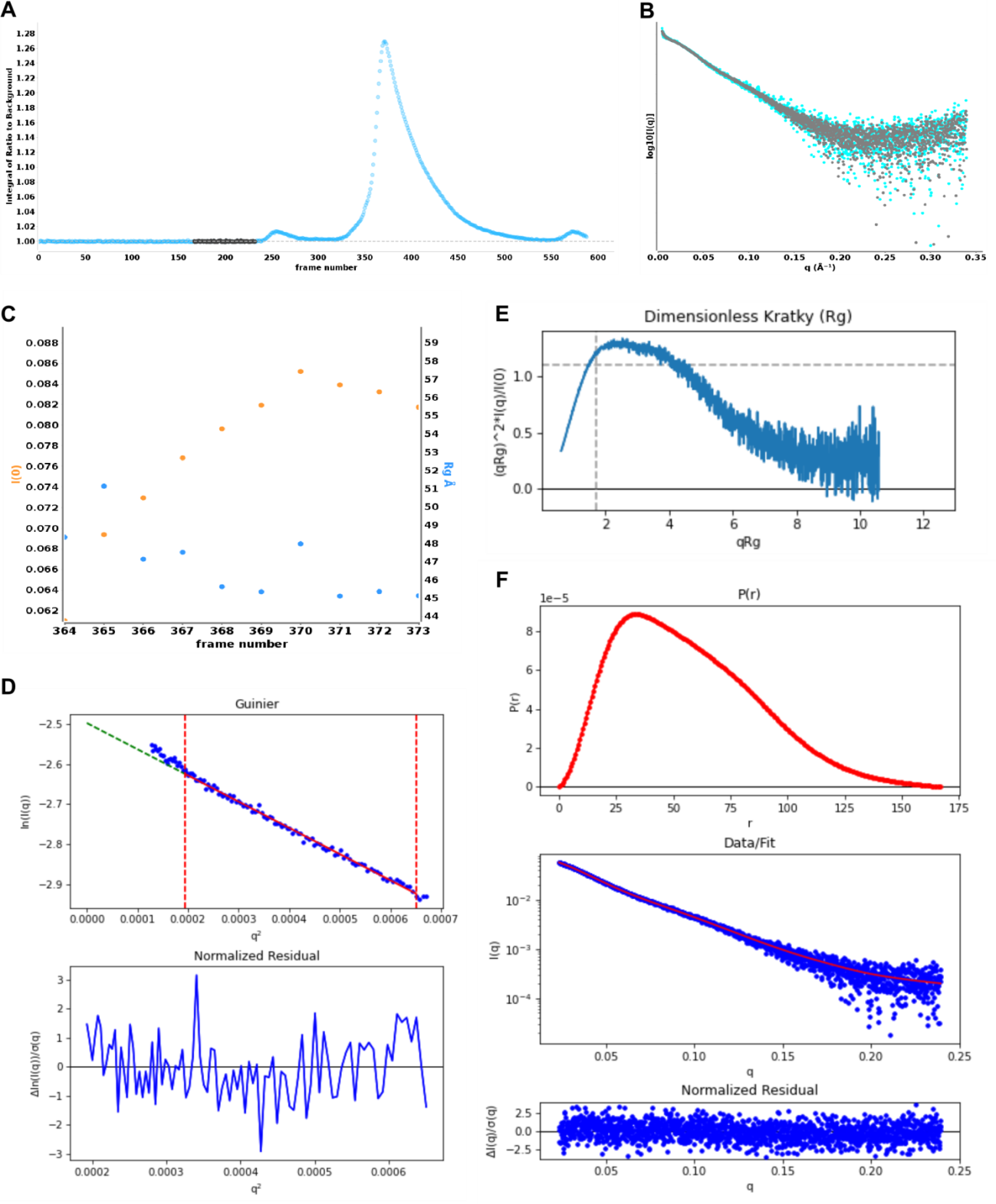

**Figure S2. SEC-SAXS of MBP-Cdc12CC.** (A) SEC profile (blue dots) showing regions used for buffer subtraction (gray dots). (B) Log10 intensity plot of subtracted and merged SAXS frames (black dots, average; cyan dots, median). (C) *I*(0) (orange) and *R*_g_ (blue) across the selected frames. (D) Guinier fit and residuals for data at *q***R*_g_ < 1.3. (E) Dimensionless Kratky plot with crosshair (1.10, 1.73) indicating the expected peak maxima for a globular protein. F) *P*(*r*) distribution (top) with the fit of the *P*(*r*) profile in red to the raw scattering data in blue (middle) and the normalized residuals (bottom).

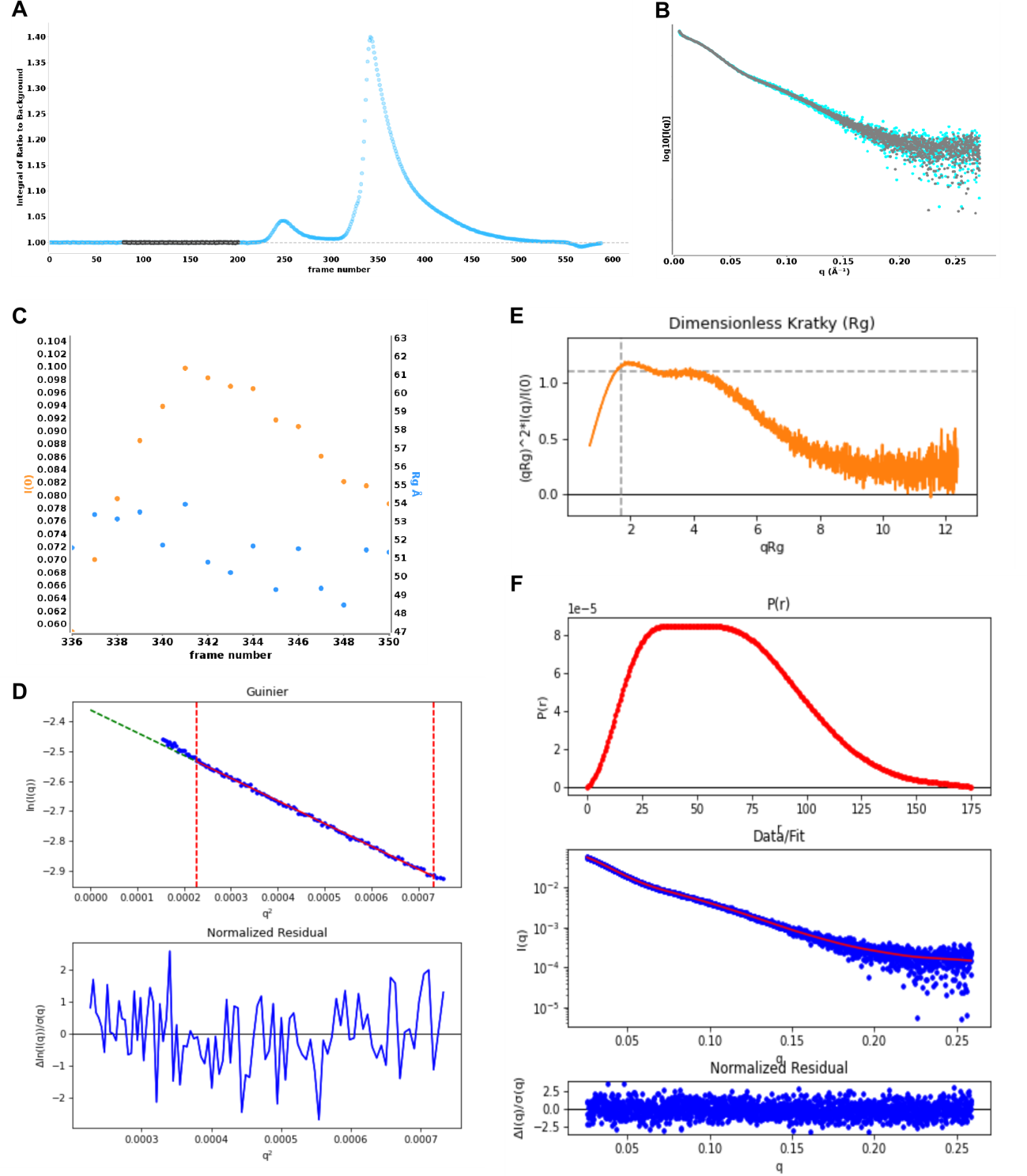

**Figure S3**. **SEC-SAXS of MBP-Cdc12CC+8.** As in Fig. S2 but for MBP-Cdc12CC+8.

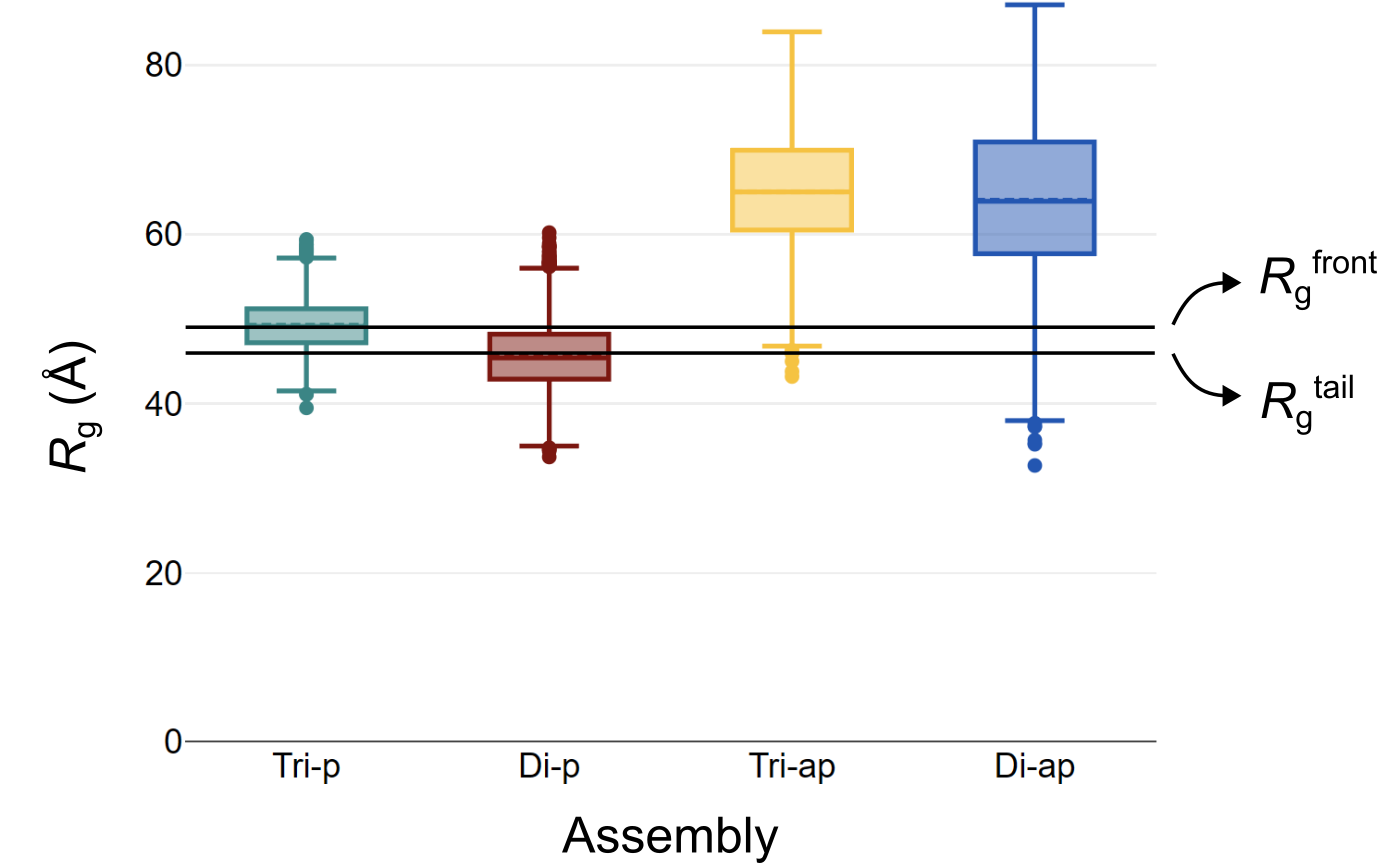

**Figure S4**. **Box plot of the radius of gyration (*R*_g_) from MBP-Cdc12CC+8 assemblies generated by RANCH.** A total of 5,000 models were made for each assembly (tri-p, parallel trimer; di-p, parallel dimer; tri-ap, antiparallel trimer; di-ap, antiparallel dimer). Real space *R*_g_’s from the peak front and the peak tail datasets (horizontal black lines) match well with the mean and median of those from tri-p and di-p models, respectively. Antiparallel assemblies were not considered for NNLSJOE analysis due to the large difference between their *R*_g_’s and the experimental *R*_g_’s.

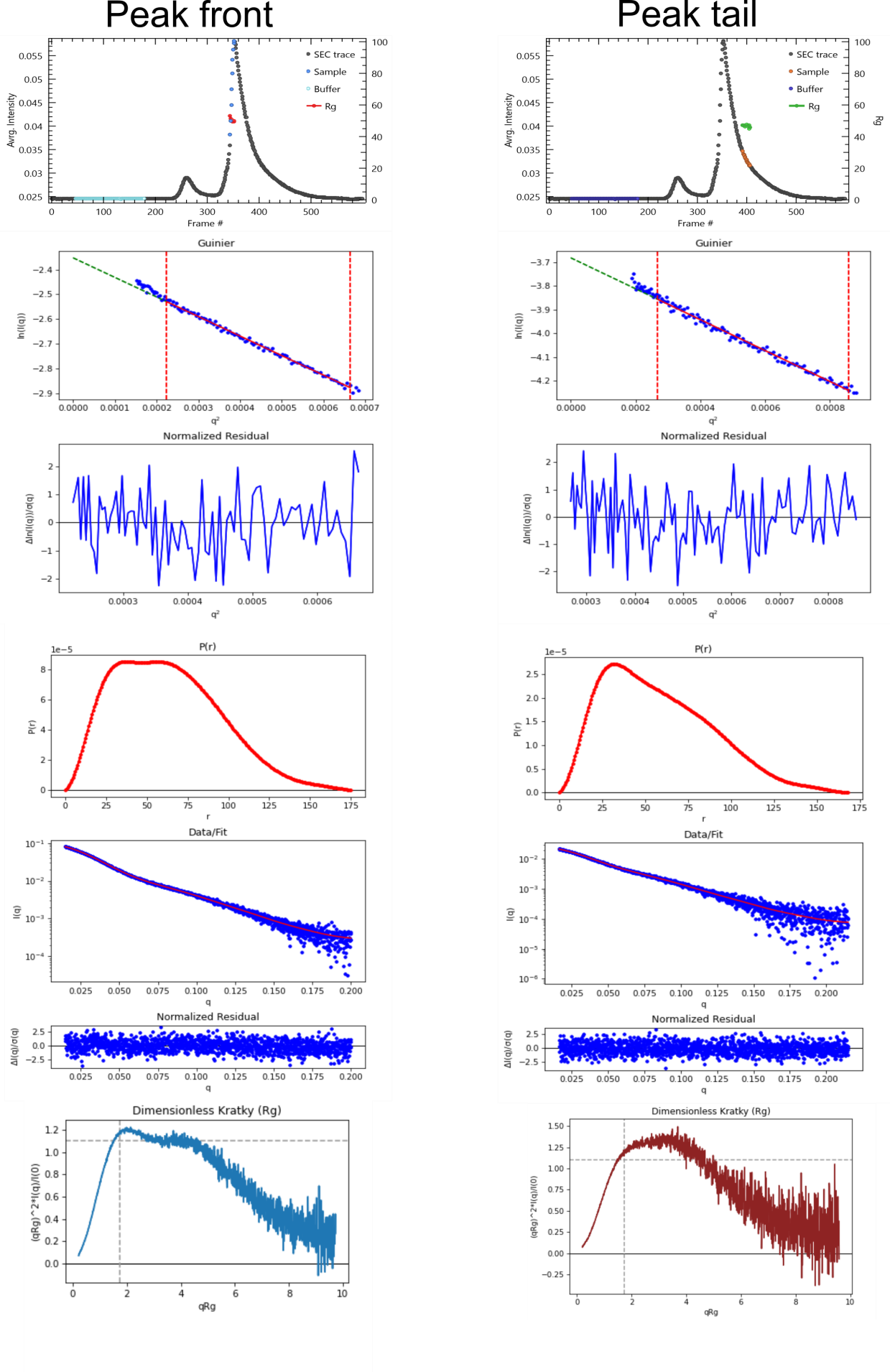

**Figure S5. SAXS analyses of MBP-Cdc12CC+8 SEC fractions.** SEC profiles, Guinier fit and residuals, *P*(*r*) distribution, fit and normalized residuals and Dimensionless Kratky plots are shown. Significant differences can be seen in the *P*(*r*) and Kratky plots.

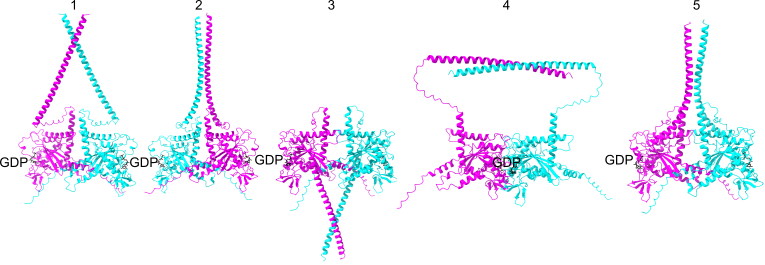

**Figure S6. Canonical septin-septin interfaces between GTPase domains in AlphaFold3 predictions of Cdc12•GDP homodimers.** Numbers indicate the confidence ranking of the models. “GDP” labels the nucleotide bound in the nucleotide-binding pocket, which is buried in the septin G interface. The septin-septin NC interface is located on the opposite side of the GTPase domain. pAE matrices are in Figure S1. Models have been deposited in ModelArchive, accession code ma-5hhc4.

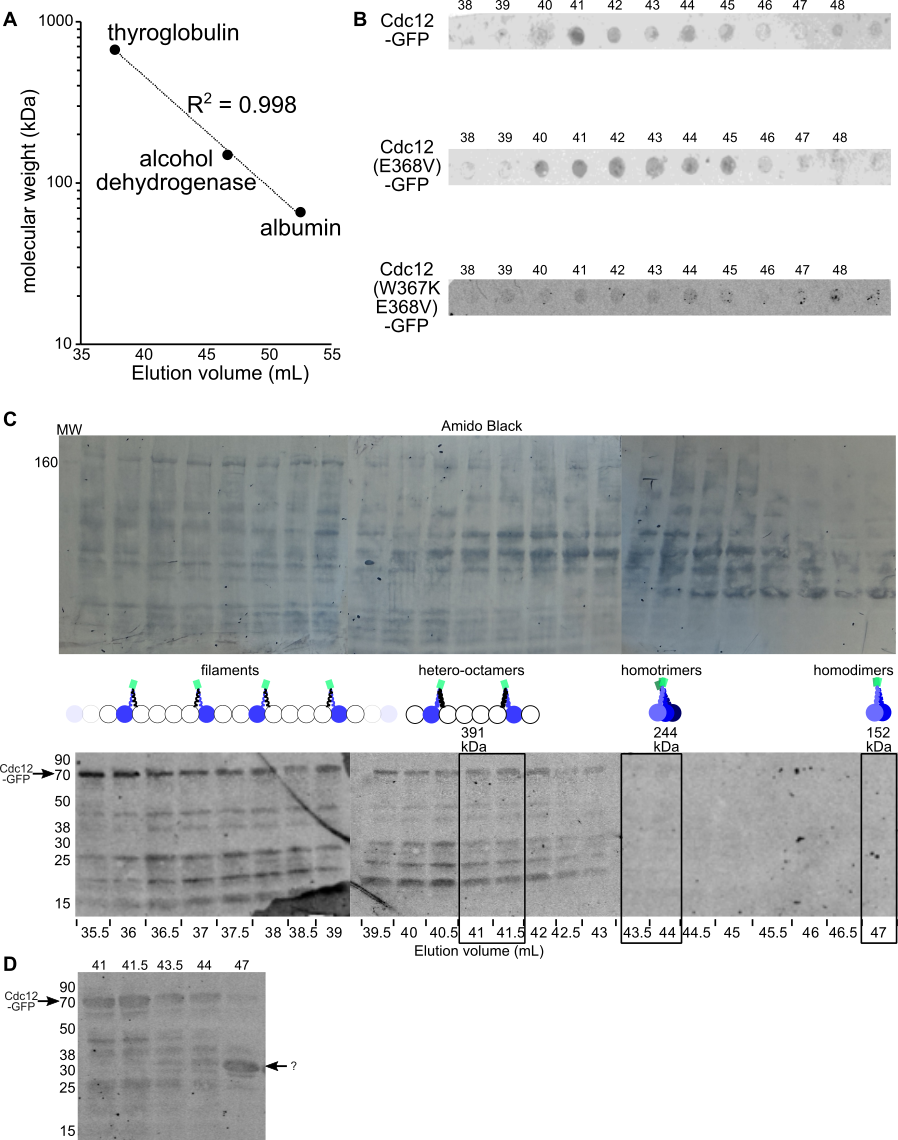

**Figure S7. Size exclusion chromatography of yeast lysates overexpressing Cdc12-GFP or mutants thereof.** (A) A mix of the indicated proteins dissolved in PBS was separated at 0.5 mL/min. Plot shows the elution volumes at each peak versus molecular weight. The R^2^ value shows the coefficient of variation of the three points fit to a line. (B) The dot blots of fractions used to quantify GFP signal for fractionated lysates of cells overexpressing the indicated Cdc12-GFP fusion. Elution volume increases left to right. (C) The experiment analyzing cells overexpressing Cdc12-GFP was repeated but 0.5-mL fractions were collected, precipitated with TCA, and resolved by SDS-PAGE prior to immunoblotting with the anti-GFP antibody. 40% of the protein volume from 24 fractions representing the indicated elution volumes was resolved on three gels. Shown at top are the three membranes stained for total protein with Amido Black after the antibody detection was complete. Below are the scanned membranes detecting anti-GFP with infrared-labeled secondary antibody. The positions of selected molecular weight standards (LICORbio #928-60000) are indicated at left; “MW” indicates this lane in the leftmost Amido Black-stained blot. (D) The remaining 60% of the five fractions boxed in (C) were resolved on a separate gel and GFP was detected in the same way. The corresponding elution volumes are indicated above each lane. The question mark indicates a band of unknown origin, likely a result of proteolysis of full-length Cdc12-GFP occurring after fractionation but before TCA precipitation.

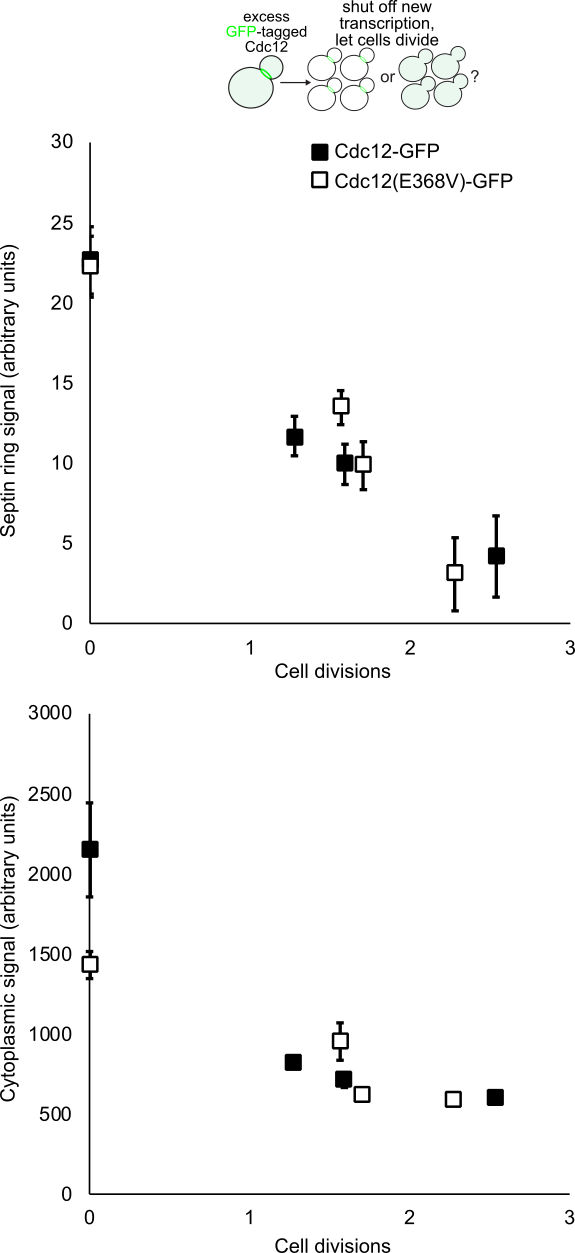

**Figure S8. Post-translational assembly into septin filaments by Cdc12-GFP and Cdc12(E368V)-GFP.** A pool of excess Cdc12-GFP or Cdc12(E368V)-GFP was generated in wild-type haploid cells (strain BY4741) carrying plasmid pMVB2 or BD7C264E by culturing cells in plasmid-selective medium containing 0.1% galactose and 1.9% raffinose, then new expression of the GFP-tagged Cdc12 was repressed by changing to medium with 2% glucose. Cell concentration per mL was monitored with a hemacytometer at various time points as the glucose cultures grew, and the number of cell divisions was calculated. At the same time points, GFP fluorescence was measured in 24-38 cells per time point per genotype via microscopy using a line scan of bud necks (septin filaments) or a circular area of the cytoplasm. Each point indicates the mean and error bars are standard error of the mean. Cytoplasmic signal is expected to decrease exponentially as the excess Cdc12-GFP is diluted via cell division; if some of the excess septins are able to assemble into septin hetero-octamers and filaments, then the septin ring signal is expected to decrease in a non-exponential manner (Hassell *et al*, 2022).

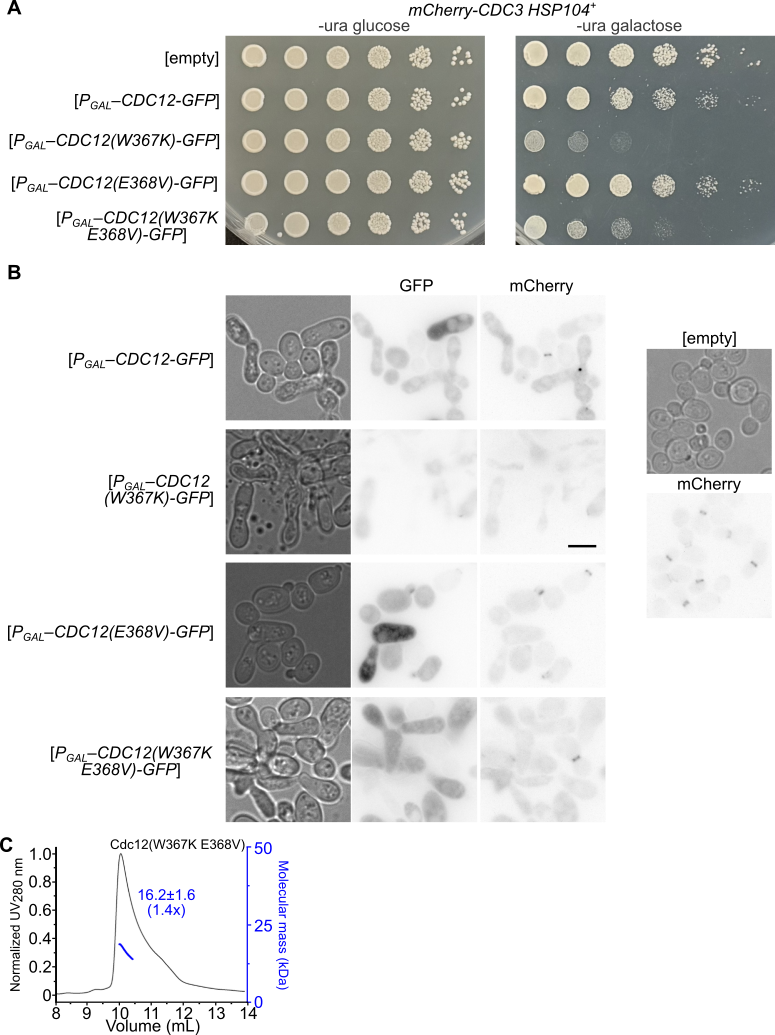

**Figure S9. Mutations altering Cdc12 CTD oligomerization state exacerbate dominant perturbation of septin function caused by overexpression of GFP-tagged Cdc12 in cells with mCherry-tagged Cdc3.** (A) Yeast strain H07151 carrying plasmid pRS316 (“empty”), pMVB2 (“*P_GAL_–CDC12-GFP*”), C6F1B99E (“*P_GAL_–CDC12(W367K)-GFP*”), (BD7C264E (“*P_GAL_–CDC12(E368V)-GFP*”), or 9F2B6E98 (“*P_GAL_–CDC12(W367K E368V)-GFP*”) was serially diluted and spotted on solid medium selective for the plasmids and either glucose or galactose as the carbon source. Plates were incubated at 30˚C for 2 days prior to imaging. (B) Cells were scraped from the galactose plates in (A) and imaged with transmitted light or the GFP (60-msec exposure) or mCherry (250-msec) cube. The cells with pRS316 (“empty”) at right were prepared and imaged the same way but in a separate experiment. Scale bar, 5 µm. (C) As in Fig. 1 D, but with Cdc12 residues 316-407 harboring both the W367K and E368V mutations.

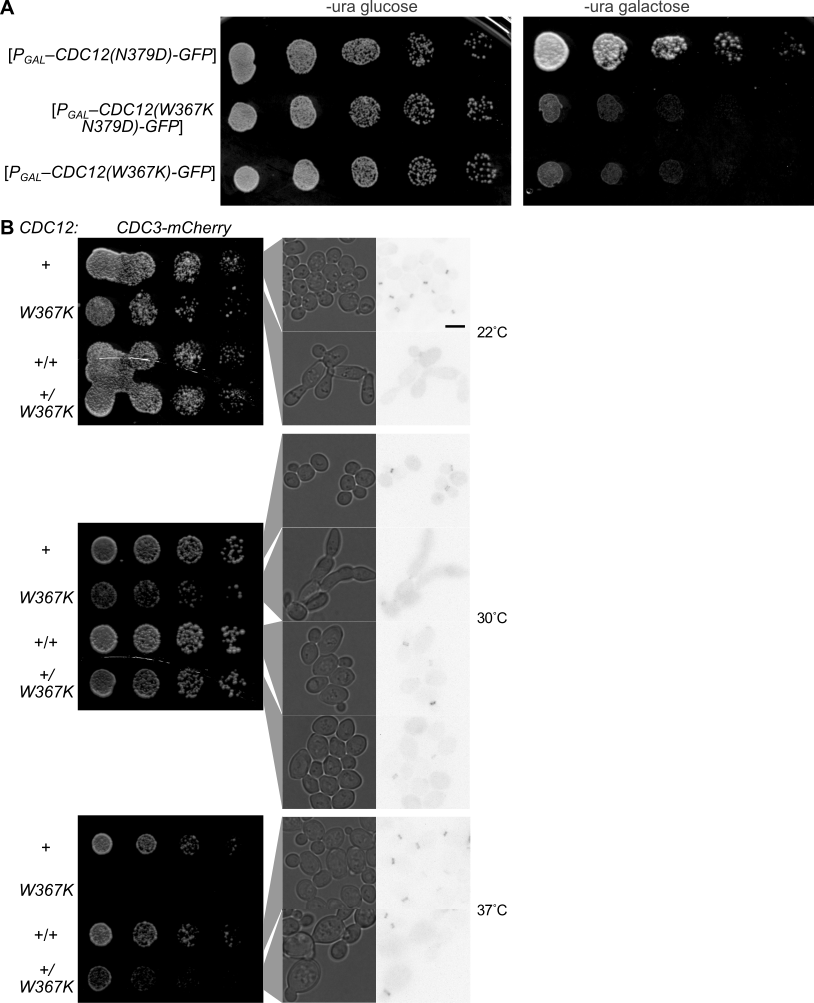

**Figure S10**. **The unanticipated mutation N379D discovered in the Cdc12-GFP-overexpressing plasmid is not responsible for the dominant lethal effects of the W367K mutation.** (A) Dilution series as in Fig. 4 A, plasmids were pMVB2 (“*P_GAL_–CDC12(N379D)-GFP*”) C6F1B99E (“*P_GAL_–CDC12(W367K N379D)-GFP*”), or 10633A79 (“*P_GAL_–CDC12(W367K)-GFP*”). (B) Dilution series as in (A) and microscopy as in Fig. S9 B but with strains lacking plasmids and in which one or no copies of the endogenous *CDC12* gene was mutated to introduce the W367K mutation, spotted on YPD agar and incubated at the indicated temperatures for two (30˚C and 37˚C) or three (22˚C) days. Haploid strains were H07151 (“+”) and 4H9K2X7P (“*W367K*”) and heterozygous diploid strains were made by mating these haploids (“+/+” and (“+/*W367K*”), respectively) with BY4742. Scale bar, 5 µm.

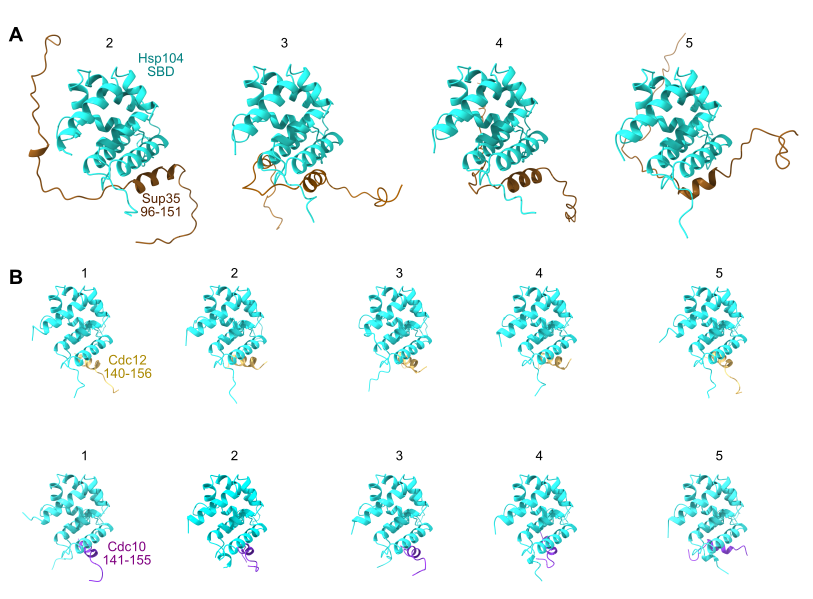

**Figure S11. AlphaFold3 predictions of Hsp104–substrate binding.** As in Fig. 6A, but showing the substrate-binding domain (SBD) of Hsp104 bound to the indicated peptides from Sup35 (deposited in ModelArchive, accession code ma-vj5d5), Cdc12 (ma-k1rre), or Cdc10 (ma-5n6qq). pAE matrices are in Figure S1.

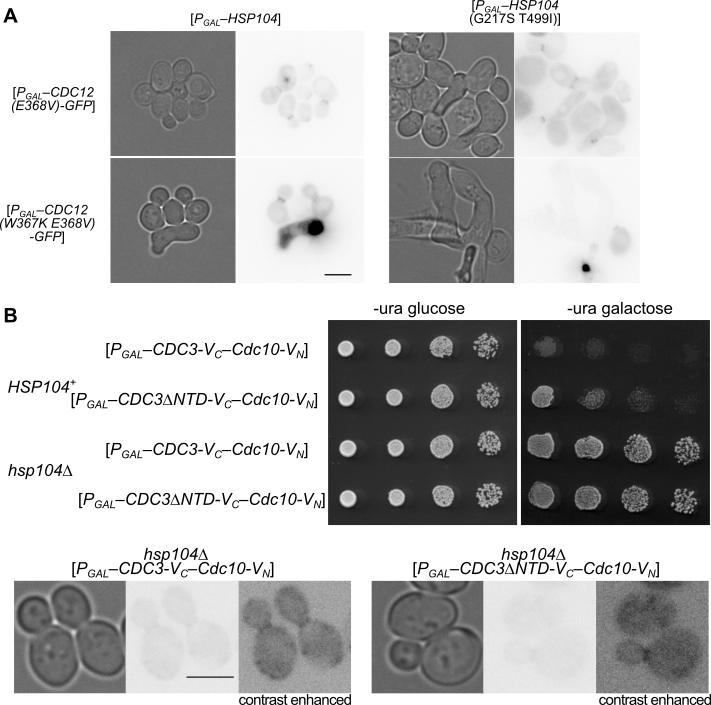

**Figure S12. Colony growth, cellular morphology, and/or septin localization upon overexpression of septins and/or Hsp104.** (A) As in Fig. 4 C, but with cells of strain BY4741 scraped from the galactose plates in Fig. 6 C. GFP images had 60-msec exposures. Scale bar is 5 µm. Plasmids were 5679 (“*P_GAL_–HSP104*”), 5707 (“*P_GAL_–HSP104(G217S T499I)*”), (BD7C264E (“*P_GAL_–CDC12(E368V)-GFP*”), and 9F2B6E98 (“*P_GAL_–CDC12(W367K E368V)-GFP*”)

(B) Top, as in Fig. 9 C but with plasmids E00432 (*P_GAL_–CDC3-V_C_–CDC10-V_N_*”), and E00435 (“*P_GAL_–CDC3∆NTD-V_C_–CDC10-V_N_*”). Bottom, cells scraped from the galactose plates were imaged with transmitted light and the GFP cube (1-sec exposures). As indicated, images with increased contrast are also provided. Scale bar, 5 µm.

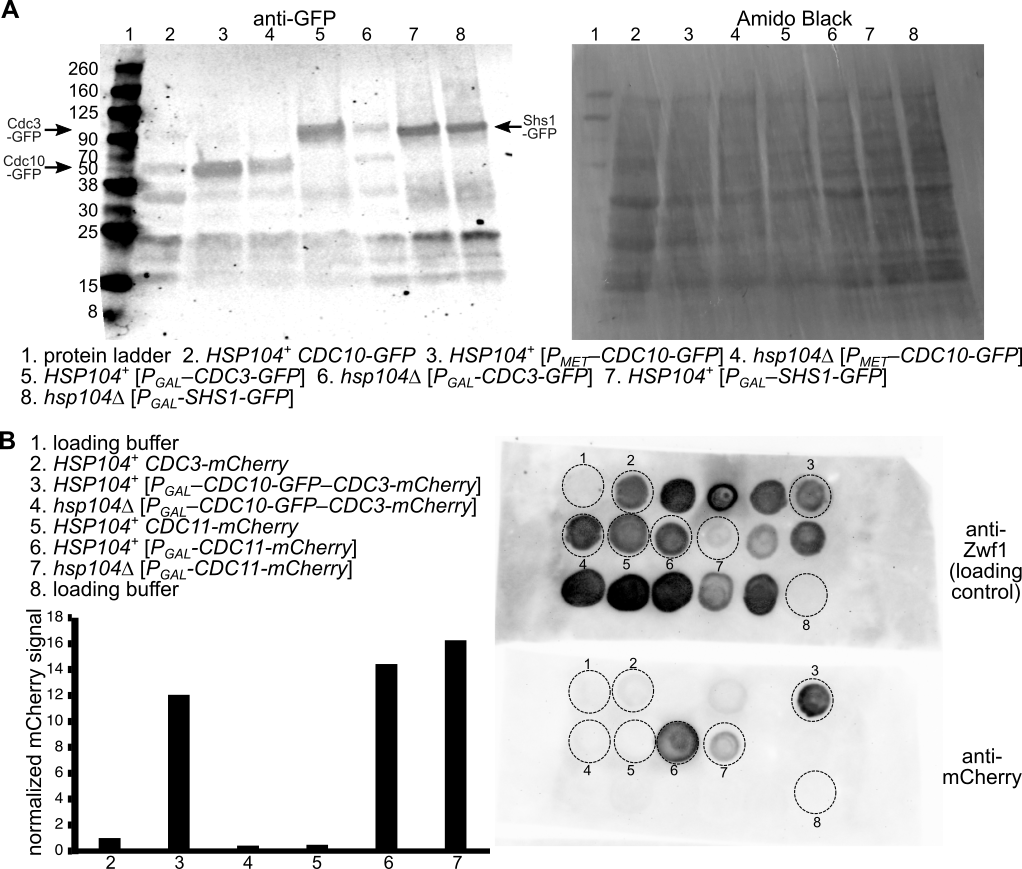

**Figure S13. Immunoblotting of levels of tagged septins.** (A) Total protein was extracted from liquid yeast cultures of the indicated genotypes grown to mid-to-late-log phase, and resolved by SDS-PAGE prior to immunoblotting with anti-GFP antibody (Sigma-Aldrich #11814460001) and infrared-labeled secondary antibody (Life Technologies Corporation #W10815). After infrared detection, the membrane was stained for total protein with Amido Black (right). Protein ladder, LICORbio #928-60000. Strains were H06798, and BY4742 or JTY4013 carrying plasmids pMETp-CDC10-GFP, pMVB1, or pGF-IVL-286, and cultures were grown in YPD (for H06798) or synthetic dropout medium with 2% galactose and lacking leucine (for Shs1-GFP) or uracil and methionine (for Cdc10-GFP and Cdc3-GFP). (B) As in (A), but protein extracts were spotted directly on membranes, and the membranes exposed to anti-mCherry or, as a loading control, anti-Zwf1 antibodies, followed by detection with peroxidase-conjugated secondary antibodies. Chemiluminescent signal was detected with an imaging system and quantified as described in the Methods. Plot shows normalized mCherry signal for the spots indicated with the key. Strains were H07151, GFY-160, and BY4741 or JTY4013 carrying plasmids YCpU-Pgal-Cdc3-mCherry-Cdc10-GFP or pGF-IVL-287 and growth was in synthetic dropout medium with 2% galactose lacking uracil or leucine, as appropriate for plasmid selection.

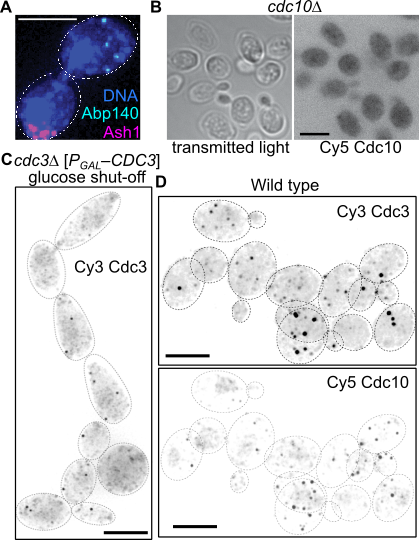

**Figure S14. Experimental controls for smiFISH detection of septin mRNAs.** (A) Representative overlay/merge image of smiFISH detection in wild-type (strain BY4743) cells of the mother-cell-localized Abp140 mRNA and the bud-tip-localized Ash1 mRNA, plus DAPI staining of the DNA. Dashed lines outline the cell walls. Scale bar, 2.5 µm. (B) SmiFISH with Cy5-labeled Cdc10 probe set, showing transmitted light and Cy5 fluorescence images of negative control (*cdc10*∆) strain JTY4010). Scale bar, 5 µm. (C) Negative control Cdc3 smiFISH. As in (B) but with Cy3-labeled Cdc3 probe set, Cy3 fluorescence detection, and *cdc3*∆ strain MMY0296 carrying plasmid pMVB169 that was cultured in glucose-containing medium to shut off *CDC3* transcription and deplete Cdc3 mRNAs. Faint spots in these cells are interpreted as non-specific signal. Scale bar, 5 µm. (D) As in (A), wild-type (strain BY4743) cells were fixed, permeabilized, and labeled with DAPI and the same Cy5-Cdc10 and Cy3-Cdc3 probe sets used in (B) and (C).

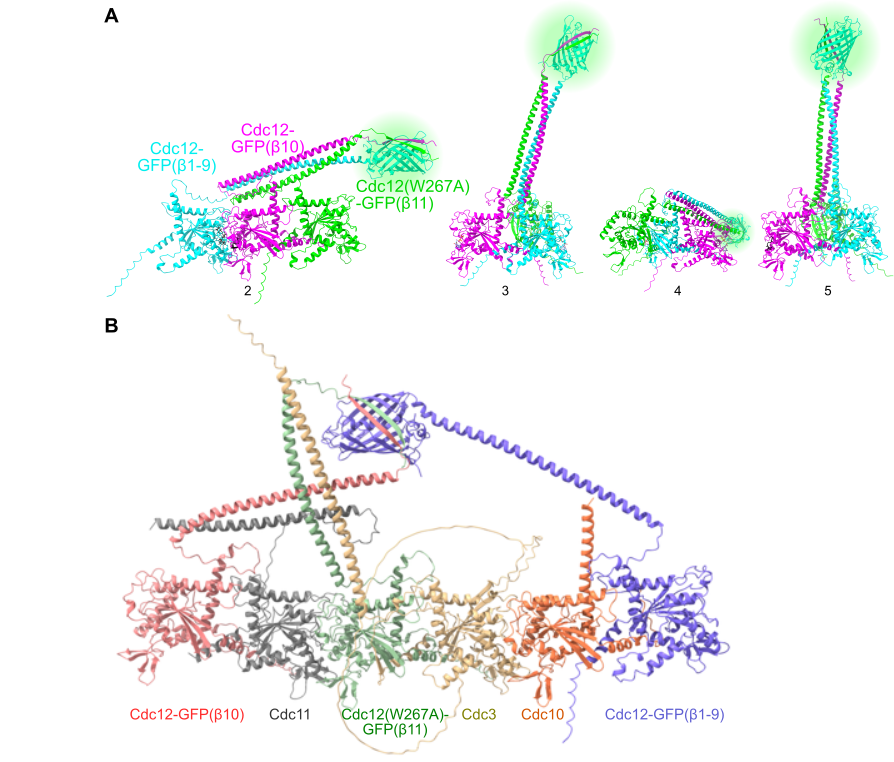

**Figure S15. AlphaFold3 predictions of Cdc12 TriFC complexes.** (A) As in Fig. 10 E, showing the other four top-ranked models. pAE matrices are in Figure S1. Models have been deposited in ModelArchive, accession code ma-s4slk. (B) The top-ranked prediction when Cdc3•GTP, Cdc10•GDP, and Cdc11•GDP were added to the prediction. pAE matrix is in Fig. S1. Model has been deposited in ModelArchive, accession code ma-37ka2.
